# Ecological Differentiation Among Globally Distributed Lineages of the Rice Blast Fungus *Pyricularia oryzae*

**DOI:** 10.1101/2020.06.02.129296

**Authors:** Maud Thierry, Florian Charriat, Joëlle Milazzo, Henri Adreit, Sébastien Ravel, Sandrine Cros-Arteil, Sonia Borron, Violaine Sella, Thomas Kroj, Renaud Ioos, Elisabeth Fournier, Didier Tharreau, Pierre Gladieux

## Abstract

Many invasive fungal species coexist as multiple lineages on the same host, but the factors underlying the origin and maintenance of population structure remain largely unknown. Here, we analyzed genetic and phenotypic diversity in isolates of the rice blast fungus (*Pyricularia oryzae*) covering a broad geographical range. We showed that the four lineages of *P. oryzae* were found in areas with different prevailing environmental conditions and types of rice grown, indicating niche separation. Pathogenicity tests revealed that specialization to rice subspecies contributed to niche separation between lineages, and differences in repertoires of putative virulence effectors were consistent with differences in host range. Experimental crosses revealed that female sterility and early post-mating genetic incompatibilities acted as strong barriers to gene flow between these lineages. Our results demonstrate that the spread of a pathogen across heterogeneous habitats and divergent populations of a crop species can lead to niche separation and reproductive isolation between distinct invasive lineages.

## INTRODUCTION

Invasive fungal plant pathogens constitute a major threat to ecosystem health and agricultural production [1–3]. Three decades of molecular plant pathology have revealed a seemingly inexhaustible stream of disease outbreaks or significant changes in the geographic location of diseases and host ranges of pathogens. In addition to the burden of emerging fungal infections, most major crops or domesticated trees are colonized by pathogens that are so widespread and have been present for so long, that they do not necessarily come to mind when one thinks about invasive fungi [4, 5]. If fungal pathogens are to spread successfully over their host species distribution, they must overcome a series of barriers to invasion related to dispersal, host availability, competition with other microbes, and abiotic conditions [6]. An understanding of the factors underlying the success of invasive fungal pathogens is essential to protect world agriculture against future pandemics [7].

Invasive fungal plant pathogens display a range of population structures, reflecting their highly diverse life-history strategies and invasion histories [6, 8, 9]. Demographic events (e.g. population bottlenecks, secondary contact) and natural selection during spread across heterogeneous environments may be associated with intricate changes in demography and life history traits, or may cause such changes. Shifting to a new host is a primary life history trait change in invasive fungal pathogens, and a major route for the emergence of disease. Changes in host range are facilitated by certain features, such as mating within or on their hosts and strong selective pressure on a limited number of genes [10, 11]. Reproduction within, or on plant hosts favors assortative mating with respect to host use, leading to a strong association between adaptation to the host and reproductive isolation [12, 13] and the differentiation of pathogen populations adapted to different host populations, varieties or species [14–17]. Such changes in pathogen host range can result from immune-escape mutations. Typically, they occur in a small number of genes encoding small secreted proteins called effectors, which enable microbes to influence the outcome of host-pathogen interactions for their own benefit, and which may be subject to surveillance by the plant immune system [18–20]. Another frequent change in the life history traits of invasive fungal pathogens is the loss of sexual reproduction [21, 22]. Invasive fungal pathogens may develop a clonal population structure over their entire introduced range or over parts of that range ([6] and references therein). The immediate causes of a the loss of sexual reproduction include a lack of compatible mating partners in the introduced range (i.e. a lack of compatible mating types) [8], hybridization and hybrid sterility [23, 24], and a lack of compatible alternate hosts for sexual reproduction [25]. Mating type loss may be driven by the extreme population bottlenecks associated with the colonization of a new host or new area. Shifts to asexual reproduction may also result from selection against reproduction [6], particularly for pathogens of domesticated crops and trees, due to the availability of large, homogeneous host populations, releasing constraints that maintain sexual reproduction in the short term [26]. These conditions may favor the asexual spread of new lineages carrying adaptive allelic combinations requiring shelter from recombination [27]. Finally, changes in life history traits can also result from variations in environmental conditions, including temperature in particular, during the course of invasive spread [28, 29]. Deciphering the complex interplay between changes in population structure and life history strategies during the spread of invasive fungi requires deep sampling the and integration of genetic, phenotypic, and environmental information throughout the native and invasive ranges.

*Pyricularia oryzae* (Ascomycota; syn. *Magnaporthe oryzae*) is a widespread model plant pathogen displaying population subdivision. *Pyricularia oryzae* is best known as the pathogen causing rice blast, which remains a major disease of Asian rice (*Oryza sativa*), but it is also a major constraint on the production of wheat (*Triticum*), finger millet (*Eleusine coracana*) and Italian millet (*Setaria italica*), a significant disease of ryegrass and an emerging pathogen of maize (*Zea maydis*) [30–35]. An in-depth genomic analysis of a large worldwide collection of *P. oryzae* isolates from 12 host plants revealed the existence of 10 distinct *P. oryzae* lineages, each associated mostly with a single main cereal crop or grass host [36]. Sequence divergence between lineages was low, of the order of about 1% [36], and analyses of gene flow and admixture provided evidence that the different host-adapted lineages were connected by relatively recent genetic exchanges, and therefore corresponded to a single phylogenomic species [37, 38]. The reproductive biology of *P. oryzae* is typical of plant pathogenic Ascomycetes. *Pyricularia oryzae* has a haplontic life cycle. Thus, the multicellular state is haploid, and the breeding system is heterothallic, with mating occurring only between haploid individuals of opposite mating types. *Pyricularia oryzae* is hermaphrodite, but successful mating requires that at least one of the partners to be capable of producing female structures (i.e. “female-fertile”).

The lineage of *P. oryzae* that infects rice is the most widely studied at the population level, due to its agricultural importance. Analyses of populations from commercial fields and trap nurseries based on molecular markers and mating assays have consistently shown that non-Asian populations of the rice-infecting lineage harbor a single mating-type and are clonal, with sexual reproduction never observed in the field [39, 40]. Populations with female-fertile strains, a coexistence of both mating types and signatures of recombination have been observed exclusively in Asia [39–43]. More recent population genomic studies of the rice-infecting lineage have revealed genetic subdivision into four main lineages, estimated to have diverged approximately 1000 years ago and displaying contrasting modes of reproduction [44–46]. Only one of the four lineages that infects rice (lineage 1, prevailing in East and Southeast Asia) displays a genome-wide signal of recombination, a balanced ratio of mating type alleles and a high frequency of fertile females, consistent with sexual reproduction [41]. All the other lineages (lineages 2, 3 and 4, present on several continents) have clonal population structures, highly biased frequencies of mating types and rare fertile females, suggesting that reproduction in these lineages is strictly asexual [41, 46]. The recombining lineage has a nucleotide diversity that is four times greater than that of the clonal lineages [46]. In theory, gene flow remains possible between the majority of the lineages, because there are rare fertile strains and some lineages have fixed opposite mating types. However, while resequencing studies have revealed signatures of recent shared ancestry between clonal lineages and the recombinant lineage from South-West Asia, this is not the case between the different clonal lineages despite their wide and overlapping distributions, suggesting a lack of recent gene flow between lineages of opposite mating types [46]. The loss of female fertility is probably a key factor in the emergence of clonal lineages, inducing a shift towards strictly asexual reproduction, and protecting populations adapting to new conditions from maladaptive gene flow from ancestral, recombining populations [10]. This raises questions about the nature of the factors underlying the emergence of different rice-infecting lineages in *P. oryzae*, and contributing to their maintenance, because, according to the competitive exclusion principle, different lineages should not be able to co-exist on the same host [47–49]. Previous studies have helped to cement the status of *P. oryzae* as a model for studies of the population biology of fungal pathogens. However, most efforts to understand the population structure of the pathogen have been unable to provide a large-scale overview of the distribution of rice-infecting lineages and of the underlying phenotypic differences, because the number of genetic markers was limited [40, 41], the number of isolates was relatively small [44, 46] and because few phenotypic traits were scored [40, 41, 44].

We provide here a detailed genomic and phenotypic overview of natural variation in rice-infecting *P. oryzae* isolates sampled across all rice-growing areas of the world. With a view to inferring and understanding the global population structure of rice-infecting *P. oryzae*, we analyzed genetic variation, using genotyping data with a high resolution in terms of genomic markers and geographic-coverage, and measured phenotypic variation for a representative subset of the sample set. We also characterized the genetic variability and content of repertoires of effector proteins using whole-genome resequencing data for a representative set of isolates. This allowed us to address the following specific questions: (i) How many different lineages of *P. oryzae* infect rice? (ii) Do lineages of *P. oryzae* display footprints of recombination? (iii) Do lineages of *P. oryzae* present evidence of recent admixture? (iv) Are the different lineages sexually competent and compatible? (v) Are the different lineages differentially adapted to host and temperature? (vi) Can we find differences in the number, identity and molecular evolution of putative effectors between lineages? (vii) Can we find differences in the geographic and climatic distribution of lineages?

## RESULTS

### The rice blast pathogen comprises one recombining, admixed lineage and three clonal, non-admixed lineages

We characterized the global genetic structure of the rice blast pathogen, using an Illumina genotyping beadchip to score 5,657 SNPs distributed throughout the genome of 886 *P. oryzae* isolates collected from cultivated Asian rice in 51 countries (Supplementary file 1). The final dataset included 3,686 SNPs after the removal of positions with missing data, which identify 264 distinct multilocus genotypes in the 886 isolates. The genotyping error rate was below 0.62% (Supplementary file 1). Clustering analyses based on sparse nonnegative matrix factorization algorithms, as implemented in the SNMF program (Frichot et al., 2014), revealed four clusters, hereafter referred to as “lineages” (Figure 1C). The model with *K*=4 clusters was identified as the best model on the basis of the cross-entropy criterion. Models with *K*>4 induced only a small decrease in cross-entropy, suggesting that *K*=4 captured the deepest subdivisions in the dataset (Figure 1 - figure supplement 1). The neighbor phylogenetic network inferred with Splitstree also supported the subdivision into four widely distributed lineages, with long branches separating three peripheral lineages branching off within a central lineage (Figure 1A; 1B). Principal Component Analysis (PCA) revealed four groups coinciding with the four lineages identified with phylogenetic network and clustering analyses. A comparison with previous findings revealed that the central lineage in the Neighbor-net network and PCA corresponds to the combination of recombining lineage 1 with lineages 5 and 6, represented by two and one individuals, respectively, in a previous study [46] (Supplementary file 2). This central lineage is referred to hereafter as lineage 1. The three peripheral lineages in the Neighbor-net network and PCA corresponded to the previously described lineages 2, 3 and 4 [46]. Lineages 2 and 3 are similar to lineages B and C, respectively, identified with microsatellite data in a previous study [41](Supplementary file 2). Genetic differentiation between the four lineages was strong and significant (Weir and Cockerham’s *F_ST_*> 0.54), indicating strong barriers to gene flow between the lineages. All genotypes from lineages 2, 3 and 4 had high proportions of membership *q* in a single cluster (*q* > 0.89), whereas shared ancestry with the other three lineages was detected in lineage 1, with 32% of the genotypes having *q* > 0.10 in lineages 2, 3 and 4 (Figure 1C). Admixture may account for the lower *F_ST_* observed in comparisons between lineage 1 and the other lineages.

**Figure 1.**
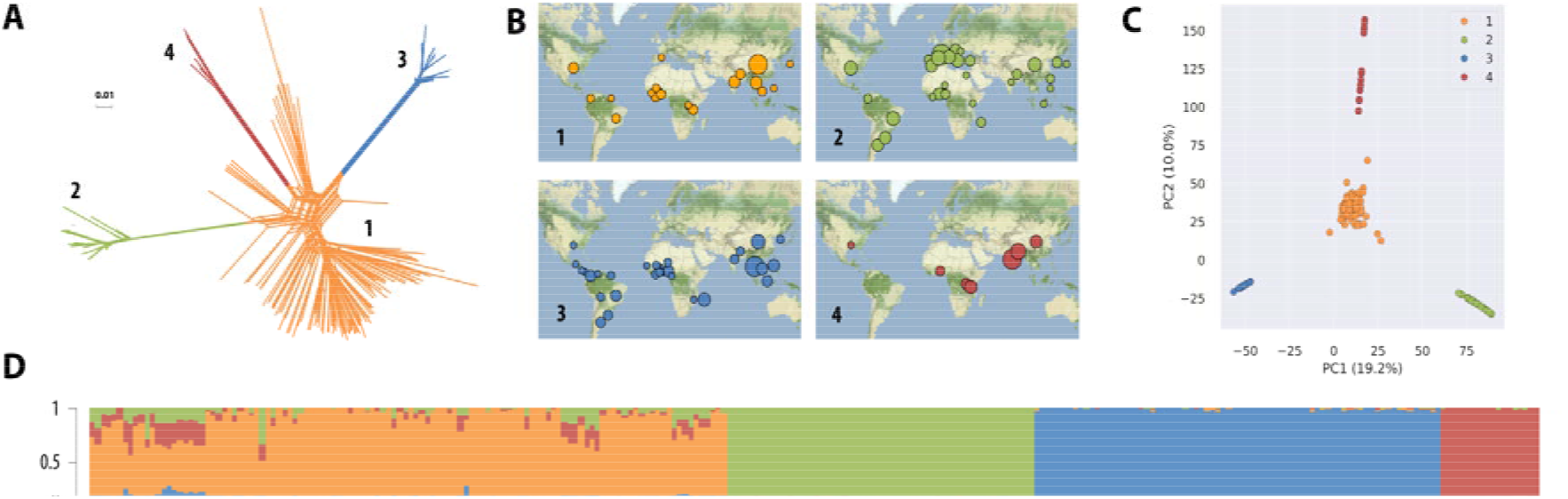
Rice-infecting *P. oryzae* populations are divided into four major lineages. Population subdivision was inferred from 264 distinct *P. oryzae* genotypes, representing 886 isolates, and the four lineages were represented in different colors. **(A)** Neighbor-net phylogenetic network estimated with Splitstree; reticulations indicate phylogenetic conflicts caused by homoplasy. **(B)** Geographic distribution of the four lineages identified with Splitstree (A) and sNMF (D), with disk area proportional to sample size. **(C)** Principal Component Analysis with colors indicating lineages identified with Splitstree (A) and sNMF (D). **(D)** Ancestry proportions in *K*=4 clusters, as estimated with sNMF software; each multilocus genotype is represented by a vertical bar divided into four segments indicating membership in *K*=4 clusters.

We detected no footprints of recombination in lineages 2, 3 and 4 with three different tests (pairwise homoplasy index [50], maximum χ^2^ [51] and neighbour similarity score [52]) and a null hypothesis of clonality (Table 1), confirming previous findings obtained with smaller datasets [41, 44-46]. The alignment of genotypes from lineages 2, 3 and 4 in the PCA (Figure 1C), their lack of linkage disequilibrium decay with increasing physical distance (Supplementary file 3), and their low proportions of homoplastic mutations (Table 1), provided additional evidence for an absence of recombination in these lineages, contrasting with lineage 1.

**Table 1.** Tests for recombination (null hypothesis: clonality), distribution of mating types, fertile females, and proportion of homoplastic polymorphisms. Analyses were conducted for four lineages, and for clusters within lineage 1.

| Lineage/Cluster | PHI (p-value) | Max $\chi^2$ (p-value) | NSS (p-value) | Homoplastic sites (%) | Mat1.1/Mat1.2 | Female-fertile isolates (%) |
| --- | --- | --- | --- | --- | --- | --- |
| 1 | <0.0001 | <0.0001 | <0.0001 | 1388/2214 (63) | 53/47 | 43 |
| *Baoshan* | <0.0001 | <0.0001 | <0.0001 | NC | 69/31 | 67 |
| *International* | <0.0001 | <0.0001 | <0.0001 | NC | 55/45 | 4 |
| *Laos* | <0.0001 | <0.0001 | <0.0001 | NC | 68/32 | 37 |
| *Yule* | <0.0001 | <0.0001 | <0.0001 | NC | 40/60 | 79 |
| 2 | 0.25 | 0.32 | 0.72 | 1/391 (0) | 97/3 | 0 |
| 3 | 0.18 | 0.35 | 0.88 | 0/408 (0) | 3/97 | 7 |
| 4 | 0.22 | 0.30 | 0.51 | 0 (218) | 97/3 | 0 |

Using whole-genome data for isolates from lineages of *P. oryzae* infecting other grasses and cereals, we showed that our set of Infinium SNPs could be used to identify isolates not belonging to the rice-infecting lineage, although it was unable to differentiate between the different host-specific lineages of *P. oryzae* (Appendix 1). Clustering analysis identified to two isolates from West Africa (BN0019 and BF0072, clonal group 18 [Supplementary file 1]) corresponding to a lineage different from the rice lineage and possibly resulting from ancient admixture between other lineages (Appendix 1). However, on inclusion of these divergent isolates in our dataset, we detected no fixed differences with respect to the rest of the isolates from lineage 1, and these divergent isolates did not form a separate group in clustering analyses. They are not, therefore, likely to have affected on our main conclusions on population subdivision and mode of reproduction.

**Figure 1 – figure supplement 1.**
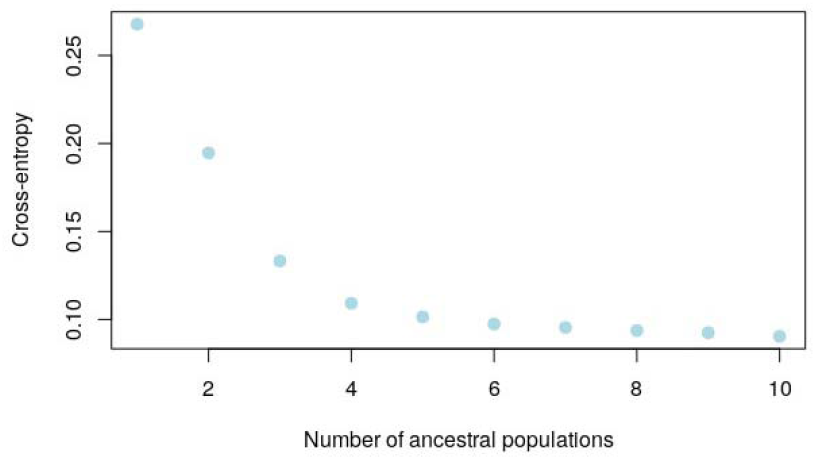
Cross-entropy (CE) as a function of the number of clusters K modeled in sNMF analyses of population subdivision. Figure 1 – source data 1. Multilocus genotypes for neighbor-net phylogenetic inference, principal component analysis and recombination analyses. Figure 1 – source data 2. Ancestry proportions in K=4 sNMF clusters.

### Geographic structure and admixture within recombining lineage 1

Lineage 1 was the only lineage for which there was population genetic evidence of recombination (Table 1). Most of the lineage 1 isolates were collected in Asia (79%), but the lineage was present on all continents in which the pathogen was sampled (Europe: one isolate; North America: 10, Central and South America: *7*, Africa: 22) (Figure 2). Clustering analysis on this single lineage with SNMF detected four clusters with different geographic distributions (Figure 2). Estimates of differentiation were lower between the clusters within lineage 1 (*F_ST_* < 0.49) than between the main lineages (*F_ST_* > 0.54), consistent with a longer history of restricted gene flow between the main lineages and a more recent history of admixture between clusters within lineage 1 (Supplementary file 3; Figure 1C).

**Figure 2.**
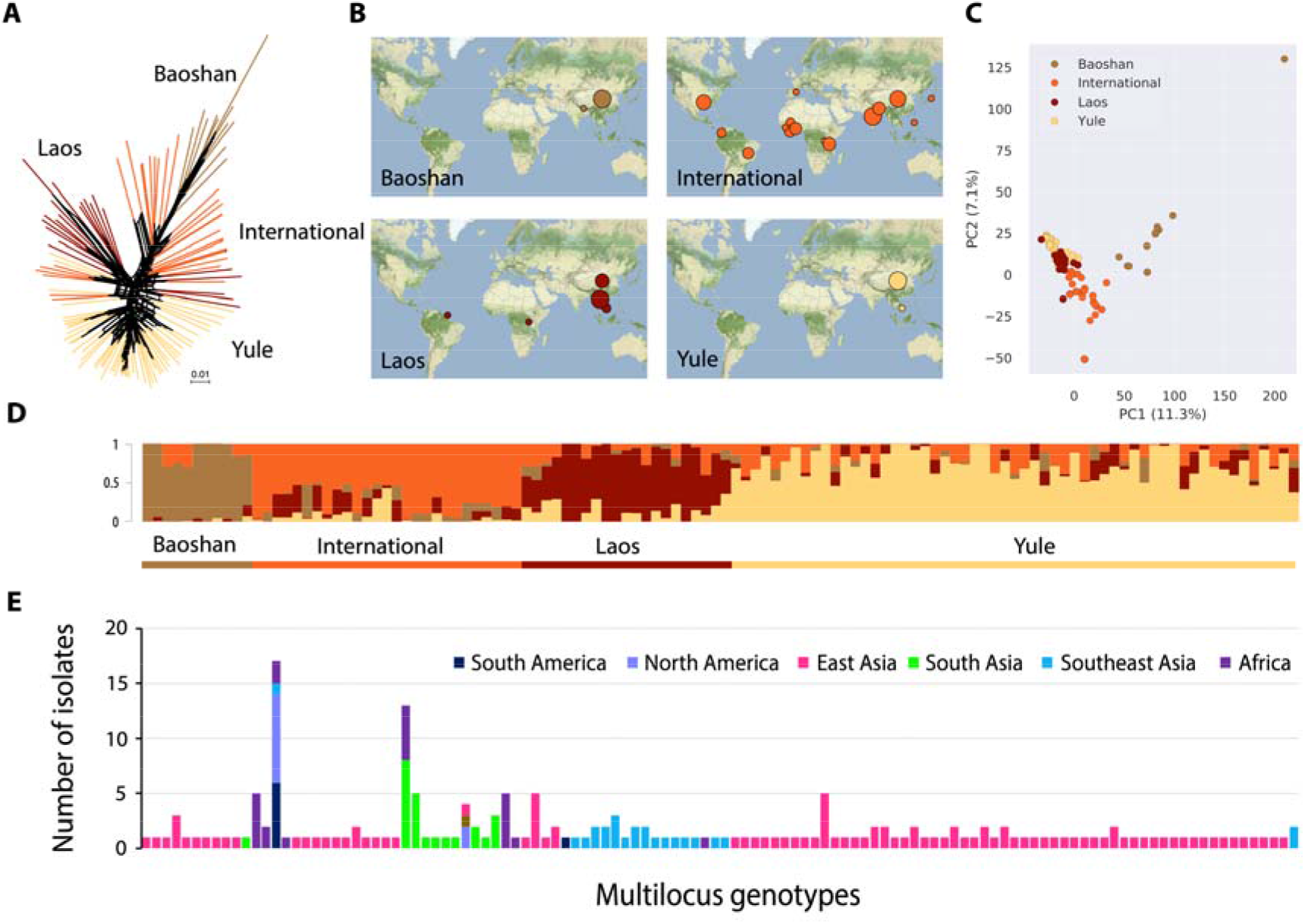
Population subdivision in lineage 1. **(A)** Neighbor-net phylogenetic network estimated with Splitstree; reticulations indicate phylogenetic conflicts caused by homoplasy. **(B)** Principal Component Analysis with colors indicating clusters identified with Splitstree (A) and sNMF (D). **(C)** Geographic distribution of the four clusters identified with sNMF (D), with disk area proportional to number of isolates. **(D)** Ancestry proportions in four clusters, as estimated with sNMF; each multilocus genotype is represented by a vertical bar divided into four segments, indicating membership in *K*=4 clusters. **(E)** Number of isolates and their geographic origin for each multilocus genotype of lineage 1. Panels D and E share *x*-axis (each vertical bar represents a different multilocus genotype). Figure 2 – source data 1. Multilocus genotypes for neighbor-net phylogenetic inference. Figure 2 – source data 2. Ancestry proportions in K=4 sNMF clusters. Figure 2 – source data 3. Geographic distribution of clones (i.e. multilocus genotypes repeated multiple times).

Two clusters within lineage 1 (referred to hereafter as *Baoshan* and *Yule*) consisted mostly of isolates sampled from two different sites in the Yunnan province of China (Baoshan and Yule, respectively, located about 370 km apart). The third cluster consisted mostly of isolates from Laos and South China (referred to hereafter as *Laos*), and the fourth brought together 95% of the isolates from lineage 1 collected outside Asia (referred to hereafter as *International*). In Asia, the *International* cluster was found mostly in the Yunnan province of China, India and Nepal (Figure 2). PHI tests for recombination rejected the null hypothesis of clonality for all four clusters (Table 1). Admixture was widespread in lineage 1 (Figure 2), with most of the isolates (78%) displaying membership proportions with *q* > 0.10 for two or more clusters (Figure 2). Only five genotypes were detected in multiple countries (genotype ID [number of countries]: 2 [6], 18 [2], 58 [2], 98 [3] and 254 [3], corresponding to 41 isolates in total). All these genotypes belonged to the *International* cluster.

### Reproductive barriers caused by female sterility and postmating genetic incompatibilities

To elucidate how the clonal lineages have emerged from the more ancient recombining populations in lineage 1, we determined the capacity of the different lineages to engage in sexual reproduction. Using *in vitro* crosses with tester strains, we analyzed the distribution of mating types and determined the ability to produce female structures (i.e. female fertility). We found that lineages 2, 3 and 4 were composed almost exclusively of a single mating type: 97% of lineage 2 and 4 isolates tested carried the *Mat1.1* allele, while 97% of lineage 3 isolates carried the *Mat1.2* allele. Only a small proportion (0-7%) of isolates from these lineages showed female fertility (Table 1). The mating type ratio was more balanced in lineage 1 (52% of *Mat1.1*; Table 1), with *Mat1.1/Mat1.2* ratios ranging from 40/60 in the *Yule* cluster to 69/31 in the *Baoshan* cluster. Female fertility rates were also high in most of the clusters in lineage 1 (*Yule*: 79%; *Baoshan:* 67%; *Laos:* 37%; Table 1). Only in the *International* cluster female fertility was as low as those in the clonal lineages, with 4% fertile females (Table 1). These observations suggest that despite the generally low rate of female fertility, sexual reproduction between most lineages is possible, except between lineages 2 and 4, that have the same highly biased mating type ratio.

We further assessed the likelihood of sexual reproduction within and between lineages, by evaluating the formation of sexual structures (perithecia), the production of asci (i.e. meiotic octads within the perithecia) and the germination of ascospores (i.e. meiospores in meiotic octads) in crosses between randomly selected isolates of opposite mating types from all four lineages (Figure 3). This experiment revealed a marked heterogeneity in the rate of perithecium formation across lineages. Clusters in lineage 1 had the highest rates of perithecia formation, with isolates in the *Yule* cluster, in particular, forming perithecia in 93% of the crosses with isolates from the same cluster, and in more than 46% of crosses with isolates from other lineages (Figure 3A). Due to their highly biased mating type ratios, isolates from lineages 2, 3 and 4 could only be crossed with isolates from other lineages. The proportion of these crosses producing perithecia was highly variable and ranged, depending on the lineages involved, from 0% to 83% (Figure 3A). In crosses involving the *International* cluster of lineage 1, the rate of perithecium formation was similar as in crosses involving clonal lineages 2, 3 and 4. None of the crosses attempted between isolates from the *International* cluster led to perithecium formation (Figure 3A).

**Figure 3.**
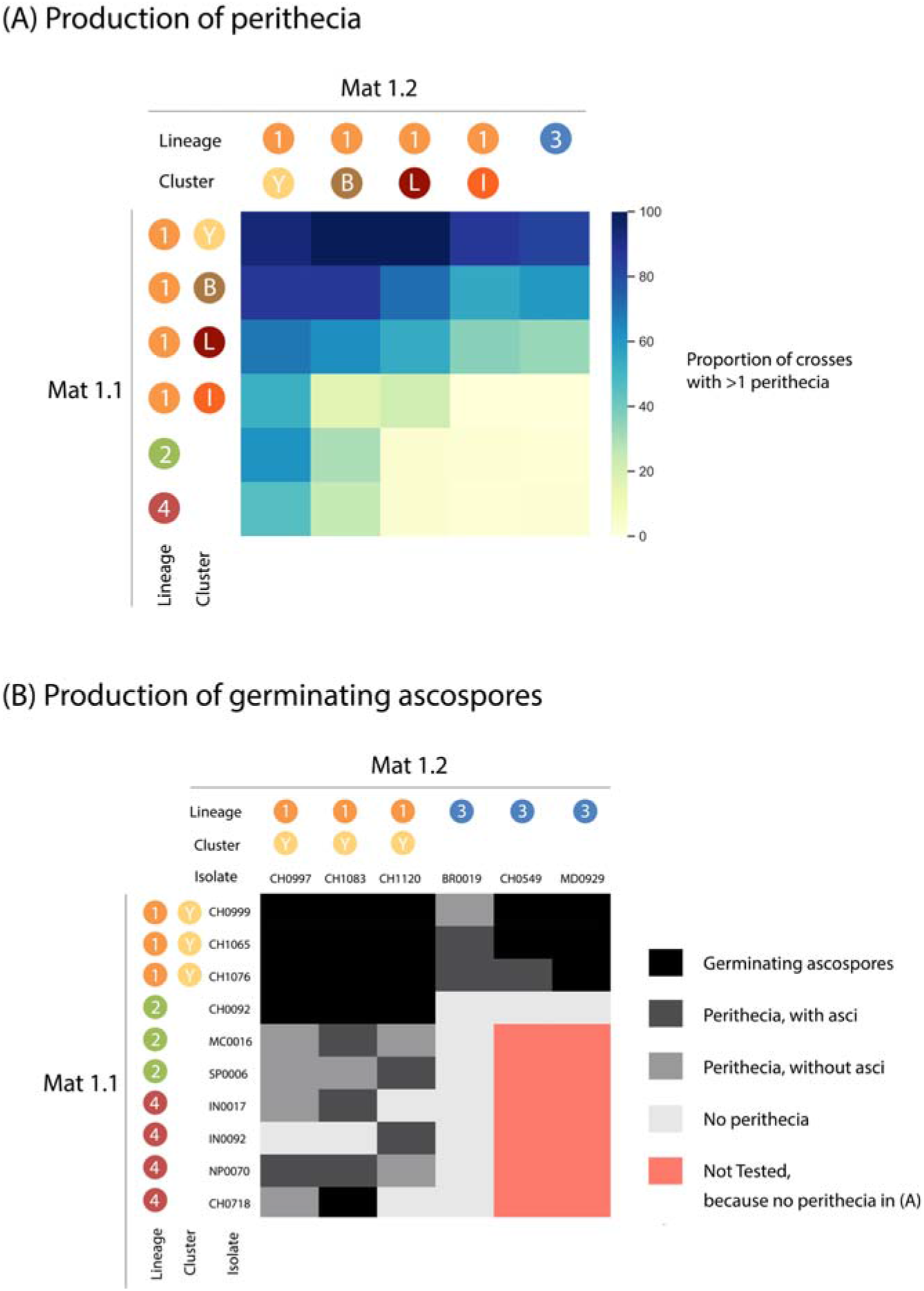
Success of crosses between lineages 2-4 and clusters within lineage 1 with (A) proportion of crosses producing at least one perithecium, (B) scoring of ascus formation and ascospore germination for a subset of crosses. Figure 3 – Source data 1 Production of perithecia Figure 3 – Source data 2 Production of germinating ascospores

Perithecium dissection for a subset of crosses involving some of the most fertile isolates revealed that most inter-lineage crosses produced perithecia that did not contain asci or contained asci with low ascospore germination rates (Figure 3B). While 100% of crosses between isolates of the *Yule* cluster produced numerous germinating ascospores, ascospore germination rates were only 33%, 56% and 7% in Yule x lineage 2, Yule x lineage 3 and Yule x lineage 4 crosses, respectively. Together, these results indicate that the three clonal lineages and the International cluster of lineage 1 are isolated from each other by strong pre- and early-postmating barriers, including breeding system isolation (differences in mating type and female sterility), and a combination of gametic incompatibility, zygotic mortality or hybrid non-viability. However, three of the clusters in lineage 1 had biological features consistent with an ability to reproduce sexually, suggesting possible sexual compatibility with clonal lineages 2, 3 and 4, and the International cluster.

### Specialization to temperature conditions and rice subspecies

Given the very broad distribution of *P. oryzae* and the strong environmental heterogeneity it encounters in terms of the nature of its hosts and the climate in which it thrives, we evaluated the variation of fitness over different types of rice host genotypes and temperature conditions. To test the hypothesis of adaptation to temperature, we measured growth rate and sporulation rate of 41 representative isolates cultured at different temperatures to test the hypothesis of adaptation to temperature (lineage 1 [yule cluster]: 11; lineage 2: 10; lineage3: 10; lineage 4: 10). For all lineages, mycelial growth rate increased with incubation temperature. This trend was more visible from 10°C to 15°C (increased mean mycelial growth of +2.22 mm/day) than from 25°C to 30°C (+0.05 mm/day) (growth curves and a full statistical treatment of the data are presented in Supplementary file 5; data are reported in Supplementary file 6). Fitting a linear mixed-effects model with incubation time, experimental replicate, and lineage of origin as fixed effects and isolate as a random effect, revealed a significant lineage effect at each incubation temperature [10°C: F(1,364)=7988, p<0.001; 15°C: F(1,419)=33161, p<0.001; 20°C: F(1,542)=40335, p<0.001; 25°C: F(1,413)=30156, p<0.001; 30°C: F(1,870)=52681, p<0.001]. Comparing least-squares means resulting from linear mixed-effects models at each temperature revealed a significantly lower growth rate of lineage 4 at 10°C relative to other lineages and a significantly higher growth rate of lineage 1 at 15°C, 20°C, 25°C and 30°C relative to other lineages. Almost no sporulation was detected at 10°C after 20 days of culture, with the few spores observed being not completely formed and divided by only one septum instead of two septa in mature conidia (sporulation curves and a full statistical treatment of the data are presented in Supplementary file 6; data are reported in Supplementary file 8). For all lineages, sporulation rates increased with temperature from 15 to 25 °C and dropped at 30°C, although isolates were cultured for only 7 days only at this temperature, versus 10 days at other temperatures, because by this stage, the mycelium had already reached the edges of the plates. Significant lineage effects were observed at 10°C, 15°C and 30°C (Kruskal-Wallis tests; 10°C: H(3)=9.61, p=0.022; 15°C: H(3)=16.8, p<0.001; 30°C: H(3)=8.96, p=0.030). Pairwise non-parametric comparisons of lineages based on Dunn’s test revealed significant differences between lineages 1 and 2 at 10°C, between lineage 1 and lineages 3 and 4 at 15°C, and between lineage 1 and 4 at 30°C. Together, measurements of mycelial growth and sporulation at different temperatures revealed differences in performance between lineages, but no clear pattern of temperature specialization.

We assessed the importance of adaptation to the host, by challenging 45 rice varieties representative of the five main genetic groups of Asian rice (i.e. the temperate japonica, tropical japonica, aus, aromatic and indica subgroups of *Oryza sativa*) with 70 isolates representative of the four lineages of *P. oryzae* and the four clusters within lineage 1 (Supplementary file 9). Interaction phenotypes were assessed qualitatively by scoring resistance (from full to partial resistance: scores 1 to 3) or disease symptoms (from weak to full susceptibility: scores 4 to 6) and analyzed by fitting a proportional-odds model and performing an analysis of variance. This analysis revealed significant differences between groups of isolates (χ^2^(6)=100, p<0.001), and between rice genetic groups (χ^2^(4)=161, p<0.001), and a significant interaction between these two variables (χ^2^(24)=97, p<0.001). The finding of a significant interaction between groups of isolates (lineages or clusters) and rice types indicates that the effect of the group of isolates on the proportion of compatible interactions differed between rice types, suggesting adaptation to the host. This is also supported by the specific behaviour of certain lineages. Isolates from lineage 2 had much lower symptom scores than the other lineages on all rice types except temperate japonica, and the isolates of the *Yule* cluster were particularly virulent on indica varieties (Figure 4A; Supplementary file 9). In comparisons of rice genetic groups, significantly higher symptom scores were observed on temperate japonica rice than on the other types of rice, whereas the varieties of the aromatic genetic group were significantly more resistant to rice blast (Figure 4B). Together, these experiments therefore revealed significant differences in host range between lineages. However, this specialization to the host was not strict, because there was an overlap between host ranges (Figure 4C-G).

**Figure 4:**
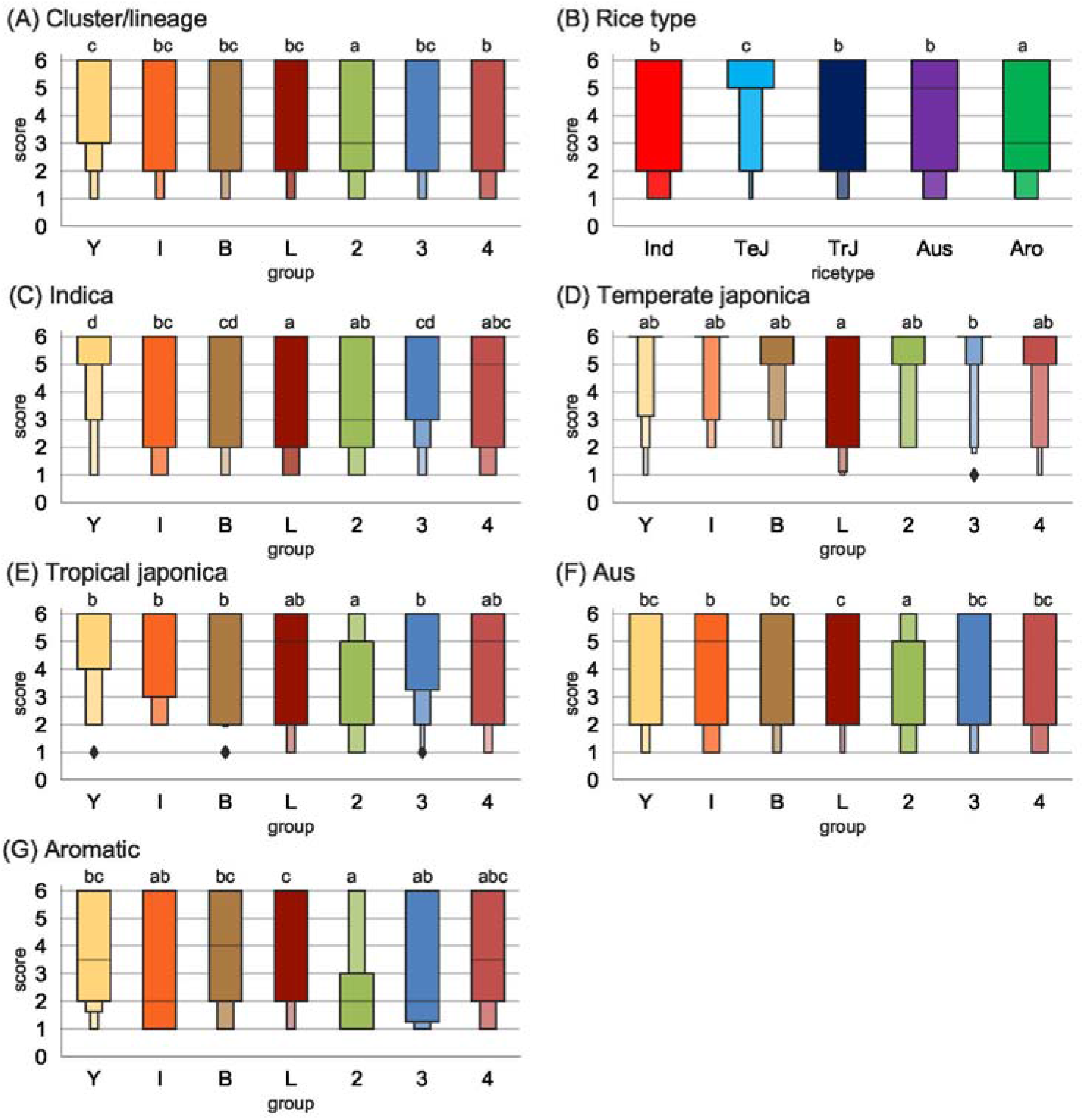
Compatibility between 70 *P. oryzae* isolates and 44 rice varieties, representing five types of rice. Compatibility was measured as symptom scores estimated using pathogenicity tests in controlled conditions. A: Symptom scores as a function of the lineage of origin of isolates, or cluster of origin for isolates from lineage 1; B: Symptom scores as a function of the type of rice; C-G: Symptom scores as a function of the lineage of origin of isolates, for each type of rice. Abbreviations: Y, *Yule*; I, *International*; B, *Baoshan*; L, *Laos*; 2, lineage 2; 3, lineage 3; 4, lineage 4; Ind, indica; TeJ, temperate japonica; TrJ, tropical japonica; Aro, aromatic. Small capitals indicate significant differences. All interactions between rice varieties and *P. oryzae* isolates were assessed in three independent experiments, and the highest of the three symptom scores was used in calculations. Boxen plots were drawn using function boxenplot() with Python package seaborn 0.11.1. Number of boxes was controlled using option k_depth=‘trustworthy’. Grey horizontal lines represent the median. Figure 4 – source data 1 Symptom scores for *P. oryzae* isolates inoculated onto rice varieties

### Differences in the number and genetic variability of putative effector genes between *P. oryzae* lineages

Effectors, defined as secreted proteins that modulate host responses, are the most prominent class of fungal virulence factors. They act by manipulating host cellular pathways and are thus key determinants of the host range and fitness on compatible hosts [53, 54]. In addition, they can be recognized by immune receptors and therefore determine host range. We therefore determined the differences in effector repertoires between *P. oryzae* lineages in terms of presence/absence and nucleotide polymorphisms, and analyzed whether this was a particularly dynamic fraction of the gene space. To this end, we used whole-genome sequencing data for 123 isolates, including 29 isolates sequenced in this study and 94 publicly available genomes [36, 44, 55], 33 of which were also genotyped with our Infinium genotyping beadchip (Supplementary file 11; Appendix 2). Clustering analysis indicated that this dataset covered the four lineages of *P. oryzae*, and three of the clusters of lineage 1 (Laos, International and Baoshan; Appendix 2; Supplementary file 11). Assembly lengths for the 123 sequenced isolates ranged from 36.7 to 39.6 Mb, with a mean value of 38.1 Mb (Supplementary file 11). The number of assembled contigs longer than 500 bp ranged from 1010 to 2438, and the longest contig was 1.1 Mb long (Supplementary file 11).

Gene prediction identified 11,684 to 12,745 genes per genome, including 1,813 to 2,070 genes predicted to encode effector proteins. We found a significant effect of the lineage of origin on both the mean number of putative effector genes (ANOVA, F=5.54, p=0.0014), and the mean number of non-effector genes (ANOVA, F=7.95, p<0.001). Multiple comparisons revealed that mean numbers of putative effectors and non-effector genes were only significantly different between the genomes of lineage 2 and lineages 3 (Tukey’s HSD test; p<0.001; Figure 5A; Figure 5B).

**Figure 5.**
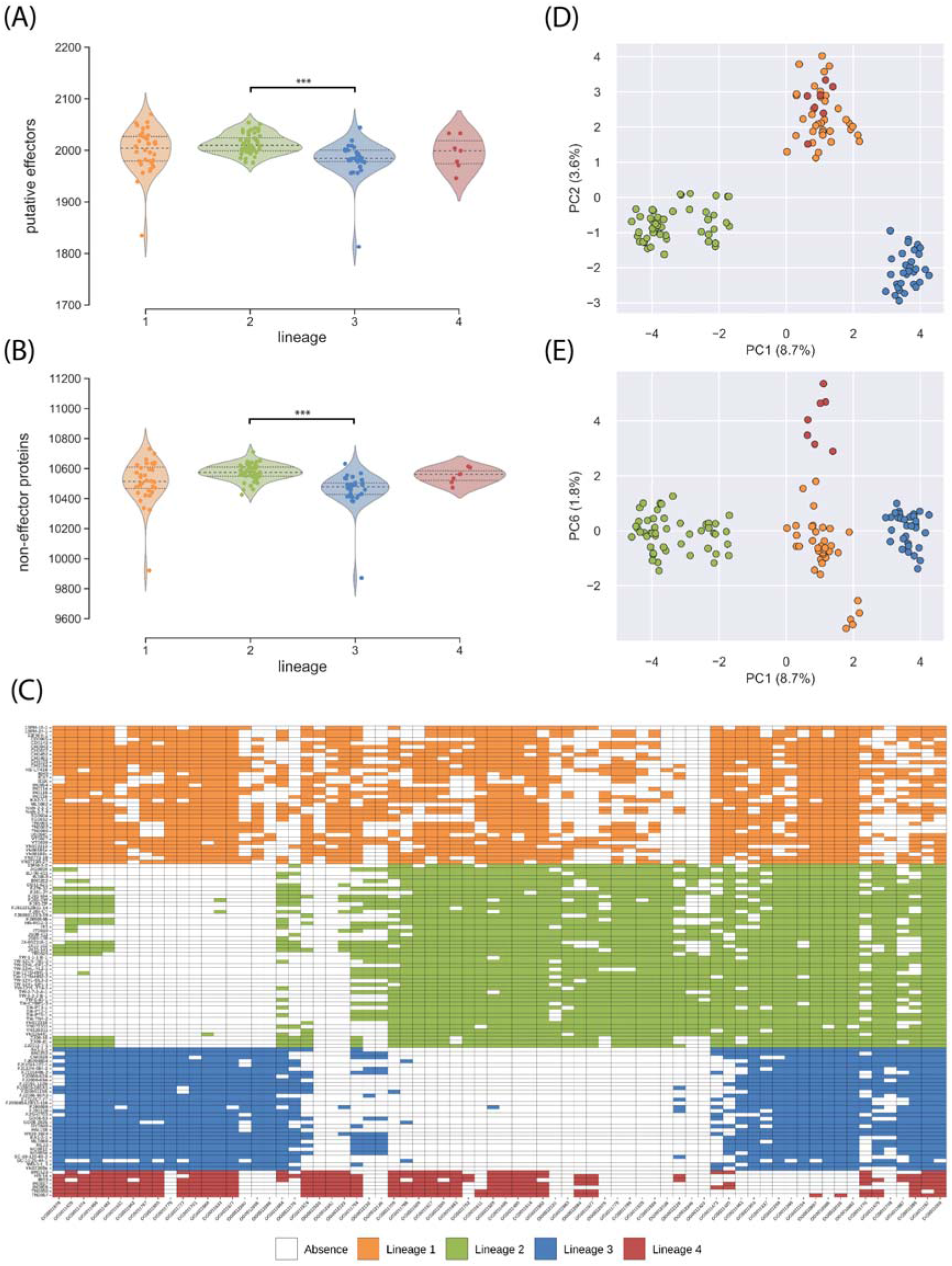
Pangenome analyses of 123 genomes of *P. oryzae*, representing four rice-infecting lineages. (A) and (B) Violin plots showing the number of putative effectors and non-effector proteins detected in each lineage, respectively; asterisks indicate significant differences in gene content (Tukey’s HSD test; p<0.001). (C) and (D) principal components PC1 against PC2, and PC1 against PC6, respectively, in a principal component analysis of presence/absence of accessory putative effectors and non-effector proteins; between parentheses is the proportion of variance represented by each principal component. (E) Presence and absence in lineage 1-4 of the 72 putative effectors with the highest contribution to principal components 1, 2 and 6, in a principal component analysis of presence/absence data. Figure 5 – source data 1 Size of core and accessory genomes for putative effectors and non-effector proteins Figure 5 – source data 2 Presence and absence of putative effectors in *P. oryzae* genomes Figure 5 – source data 3 Presence and absence of non-effector genes in *P. oryzae* genomes Figure 5 – source data 4 Orthology table, with one orthogroup per line and one isolate per column

To estimate the size of accessory and core genomes in the different lineages, we performed an orthology analysis that identified 14,573 groups of orthologous sequences (i.e. orthogroups), and we applied a rarefaction approach to the table of orthology relationships to estimate the size of accessory and core genomes in lineages while accounting for differences in sample size. For both putative effectors and the remaining of the gene space, we found that the number of core and accessory genes did not reach a plateau. Instead, they displayed an an almost linear relationship with the number of genomes resampled. This indicates that the number of core genes was probably much smaller, and the number of accessory genes substantially higher than estimated from 123 genomes (Supplementary file 12). With pseudosamples of size n=30 per lineage (i.e. excluding lineage 4, due to its small sample size), the gene content was highly variable within lineages with only 61-71% of all predicted effectors, and 68-73% of the remaining of the gene space being conserved in lineages 1-3 (Supplementary file 13). Despite extensive variation of the gene content within lineages, the clustering of isolates based on presence/absence variation for both putative effectors and the remaining of the gene space was highly similar to that based on 3,868 SNPs (Figure 5D; Figure 5E), indicating that variation in gene content and SNP allelic variation reflected similar genealogical processes.

For the identification of putative effectors potentially involved in the host specialization of rice-infecting lineages of *P. oryzae*, we first identified orthogroups with different patterns of presence/absence across lineages. Principal component analysis on presence/absence data identified 72 orthogroups of putative effectors accounting for 95% of the variance of the three principal components differentiating the four lineages (Figure 5C; Figure 5D; Figure 5E), and potentially contributing to the differences in host range between lineages.

Host specialization can also involve sequence divergence for effector proteins. We therefore also scanned the corresponding genes for signatures of diversifying selection, i.e. an excess of non-synonymous nucleotide diversity (ratio of non-synonymous to synonymous nucleotide diversity π_N_/π_S_ >1). We identified 185 orthogroups with π_N_/π_S_ > 1 in at least one lineage, including 164 orthogroups with π_N_/π_S_ > 1 in only one lineage (lineage 1: 131 orthogroups; lineage 2: 18; lineage 3: 10; lineage 4: 5), and twelve, seven and two orthogroups with π_N_/π_S_ > 1 in two, three or four lineages, respectively (Supplementary file 14). None of these orthogroups corresponded to effectors previously characterized as being involved in *P. oryzae* virulence on rice or other *Poaceae* hosts.

### Geographic and climatic differentiation in the distribution of *P. oryzae* lineages

Our tests of adaptation to host and temperature under controlled conditions showed that specialization was not strict, but it remained possible that fitness differences between lineages would be sufficient under natural conditions to induce separation in different ecological niches, and/or that our experimental conditions did not capture the full extent of the phenotypic differences. Here, we tested the hypothesis that lineages thrive in geographically and/or environmentally different conditions, which would provide indirect evidence for specialization to different habitats. We tested this hypothesis by collecting geographic and climatic data for all isolates, using the GPS position of the nearest city or the geographic center of their region of origin when the exact GPS position of isolates was not available (Supplementary file 1). At a larger scale, clonal lineages 2 and 3 were both widespread, with lineage 2 found on all continents, and lineage 3 on all continents except Europe. Lineage 2 was the only lineage sampled in Europe (with the exception of a single isolate from lineage 1), whereas lineage 3 was more widespread in intertropical regions. Lineage 4 was mostly found in South Asia (India, Bangladesh, Nepal), and in the USA and Africa (Benin, Tanzania). The recombining lineage, lineage 1, was found on all continents, but mostly in Asia (79% of all isolates and 94% of all genotypes). At a smaller scale, two, or even three different lineages were sampled in the same year in the same city in 11 different countries from all continents. However, in only one instance did two isolates from different lineages have identical GPS positions (isolates US0106 and US0107 sampled from the same field). Thus, lineages had different, but partially overlapping distributions, including some overlap with the distribution of the sexually reproducing lineage (lineage 1).

We then investigated whether differences in the geographic range of lineages were associated with differences in climatic variables. By plotting sampling locations onto a map of the major climate regions [56] we were able to identify clear differences in the climatic distributions of lineages 2 and 3, with lineage 2 mostly found in warm temperate climates and lineage 3 found in equatorial climates (Figure 6). We used the outlying mean index (OMI), which measures the distance between the mean habitat conditions used by a lineage and the mean habitat conditions used by the entire species, to test the hypothesis that different lineages are distributed in regions with different climates. Using information for 19 climatic variables (biomes) from the WorldClim bioclimatic variables database [57] for all sampling locations, we obtained statistically significant results, in a permutation test on OMI values, for lineages 2, 3 and 4 (permutation test: lineage 1: OMI=2.14, p=0.620; lineage 2: OMI=6.54, p<0.001; lineage 3: OMI=2.08, p<0.001; lineage 4: OMI=13.73, p=0.017). The OMI analysis, in which the first two axes accounted for 69% and 25% of the variability, respectively, revealed that lineage 2 was more frequent in regions with a high annual temperature range (biome 7) or a high degree of seasonality (biome 4); lineage 4 was associated with regions with high levels of seasonal precipitation (biomes 13, 16 and 18), and lineage 3 was more frequent in regions with high temperatures (biomes 1, 6, 10 and 11) and high levels of isothermality (biome 3), characteristic of tropical climes (Source code 2; Figure 6; Supplementary file 15).

**Figure 6.**
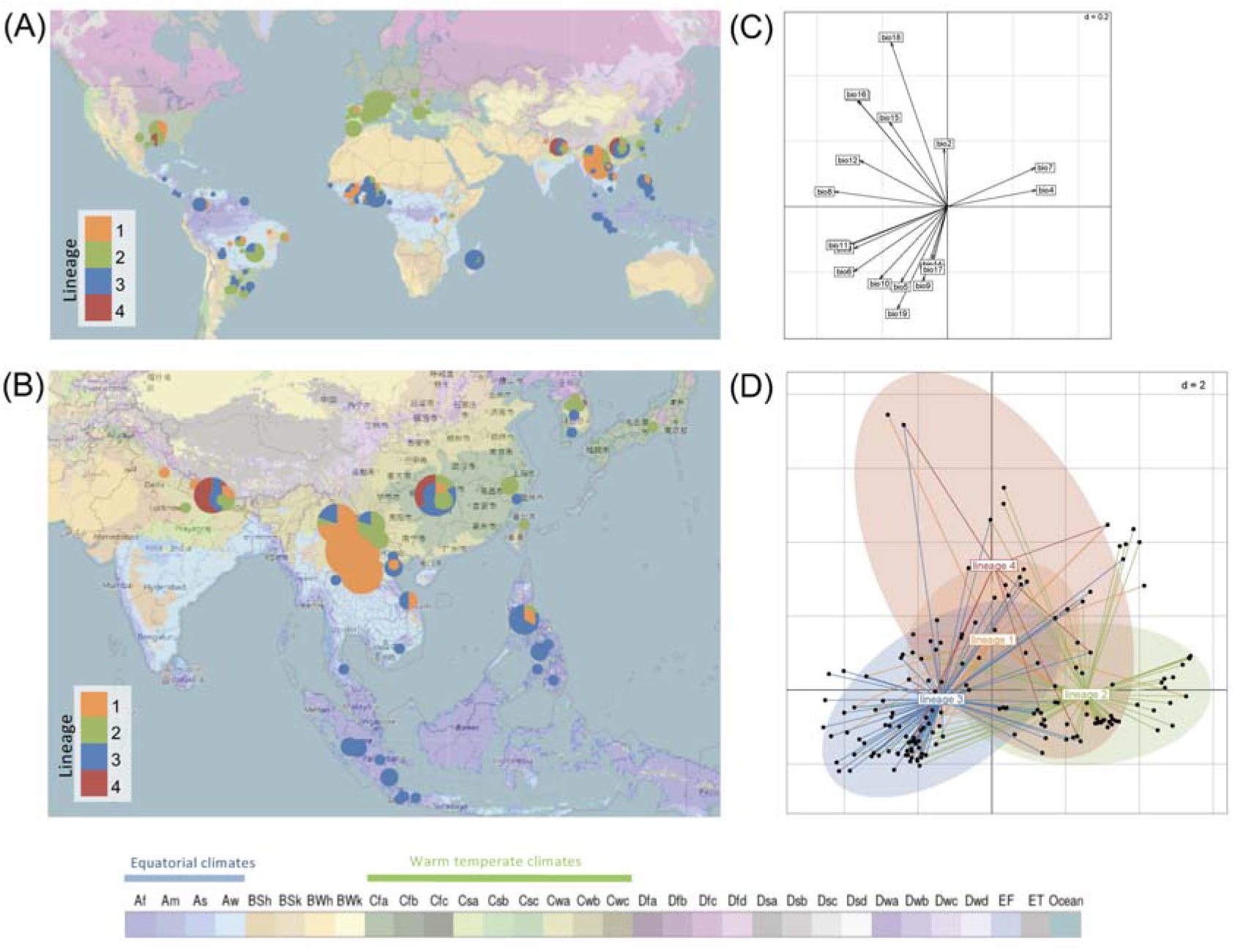
Geographic distribution of four lineages of *P. oryzae* and corresponding climatic data. (A, B) Pie charts representing the distribution of the four lineages in the world (A) and in South, East and Southeast Asia (B), keeping only isolates for which the sampling position was precisely known (i.e., for which the region, city or GPS position was documented). Background map is from OpenStreetMap (CC-BY-SA) and represents the updated Köppen-Geiger climate classification of main climates at world scale as described in [56]. (C, D) Outlying Mean Index analysis, testing the separation of ecological niches among lineages, with (C) canonical weights of the 19 environmental variables included in the analysis, and (D) site coordinates (dots) and realized niches of lineages represented as ellipses. Variable bio13 co-localizes with bio16, and variables bio1 and bio3 co-localize with bio11. The 11 temperate variables included in the analysis are listed in the Methods section. Figure 6 – source data 1 GPS position of isolates

## DISCUSSION

This study provides a detailed overview of the worldwide distribution of genetic and phenotypic diversity in *P. oryzae*, and of the eco-evolutionary factors underlying the emergence and maintenance of multiple divergent lineages in this widespread pathogen. We detected three clonal lineages and one recombining lineage with broad and largely overlapping distributions, corresponding to the four groups previously identified on the basis of fewer markers or fewer isolates [40, 41, 44-46]. Our work extends these previous findings by revealing that the lack of recombination and recent admixture in widespread lineages persists with deeper sampling, and by providing evidence for the subdivision of the recombining lineage into one international and three Asian clusters. This deep sampling of the natural genetic variation also reveals that, despite distributions displaying a broad overlap over large scales, rice blast lineages thrive in areas differing in terms of the prevailing environmental conditions and types of rice grown. We also provide experimental evidence that host-specialization, female sterility and early post-mating genetic incompatibilities act as strong barriers to gene flow between lineages. These results highlight that pathogen spread over heterogeneous habitats and over divergent populations of a crop species may create niche separation and barriers to gene flow between different invasive lineages.

### Niche separation and barriers to gene flow between clonal lineages

Many invasive plant pathogens consist of multiple divergent lineages coexisting on the same host, over large areas. This raises questions about the maintenance of different lineages in the face of potential gene flow and competitive exclusion. We show that female sterility and intrinsic genetic incompatibilities represented strong barriers to gene flow between the clonal lineages, potentially accounting for their maintenance over time, without fusion, despite the compatibility of mating types. These barriers to gene flow may have contributed to the establishment of lineage specialization because reproductive isolation facilitates adaptation by limiting the flow of detrimental ancestral alleles in populations adapting to a new habitat. This specialization accounts for the maintenance of the clonal lineages on the same host species, when faced with competitive exclusion.

Our analyses also show that, despite widely overlapping large-scale distributions, the different lineages are essentially found in different regions, with different climatic characteristics, when their distributions are observed at a finer scale. Lineage 1 was mostly sampled in Southeast Asia, lineage 4 predominated in India, and lineages 2 and 3 had global distributions. However, an analysis of climatic data indicated that lineage 2 predominated in temperate climates, where temperate japonica rice is grown, lineage 3 predominated in tropical climates, in which indica rice is grown, and lineage 4 predominated in regions with high levels of seasonal precipitation in which the indica and aromatic rice types are the principal rice types grown. Despite the finding of separation in different climatic regions, our experiments revealed no strong differences between lineages in terms of sporulation and mycelial growth on synthetic media at different temperatures. This suggests that, if adaptation to temperature occurs in this pathogen, it was not measured by our experiments, either because it involves traits other than sporulation and hyphal growth, or because the *in vitro* conditions were not suitable to demonstrate differences. The host range varied across lineages, but all host ranges overlapped, indicating that host specialization is not strict, or was not fully captured by our experiments. Nevertheless, adaptation to the host or small differences in temperature optima may, nonetheless, further increase niche separation and reduce gene flow between lineages, via pre-mating barriers, such as immigrant non-viability (i.e., lower rates of contact between potential mates due to the mortality of immigrants; [11, 58]), and post-mating barriers, such as ecological hybrid non-viability (i.e., lower survival of ill-adapted hybrid offspring).

### Pestification can foster lineage divergence

There is a direct connection between disease emergence and speciation, because the emergence of new pathogens can be facilitated, caused by or associated with strong divergent selection in organisms mating within or on their hosts, and by major changes in demographics and geographic isolation [12, 59-61]. There is also a connection between disease emergence and mode of reproduction, because disease emergence should be facilitated by multiple asexual cycles, corresponding to multiple cycles of selection for local adaptation without migration introducing locally deleterious immigrant alleles and recombination breaking down locally advantageous allelic combinations [12]. However, theoretically, in the long term, asexual organisms should accumulate a greater load of deleterious mutations and be less able to fix advantageous mutations, resulting in higher rates of extinction and lower rates of speciation [62, 63]. The pattern of population structure revealed in *P. oryzae* is seems to be intermediate between observations for invasive fungi with fully clonal population structures — such as the pathogens causing wheat brown rust [25], boxwood blight [64] or verticillium wilt [65, 66] — and invasive fungi with barriers maintaining invasive lineages that are permeable and allow hybridization and introgression to occur, such as the pathogens causing *Dothistroma* needle blight [67], Dutch elm disease [8], soybean anthracnose *Colletotrichum truncatum* [68], and coffee rust [69, 70]. The global structure of *P. oryzae* is more reminiscent of that of the pathogens causing chestnut blight [71, 72], wheat yellow rust [73, 74], and hop powdery mildew [75], for which clonal invasive lineages coexist with recombining lineages, although the proximal causes for the loss of sexual reproduction may differ between these models and rice blast. Another marked difference from most of the abovementioned pathogens is that the clonal lineages of rice blast are relatively older, having diverged about a thousand years ago [44–46], which corresponds to several thousand asexual generations. Recombining lineage 1 may, at some point, become a source of adaptive introgression, reinvigorating the genomes of clonal lineages to reduce the mutational load, introducing beneficial variation, and leading to marked changes in population structure. Thus, although our work shows that pestification can drive the emergence of reproductive barriers, we cannot predict whether these lineages will continue to exist in their current form for the foreseeable future.

### Loss of female sterility as the proximal cause of shifts to clonality

The loss of sexual reproduction, caused by demographic events or selection against sex, is a frequent change in the life history traits of invasive fungal pathogens [6, 76]. However, the underlying changes and driving factors remain largely unexplored. In *P. oryzae*, the release of constraints maintaining sex in the short term, such as the need for sexual structures for winter survival or long-distance dispersal, is probably not responsible for the loss of sex, because the likelihood of overwintering on seeds and the rate of human-mediated transportation would not be expected to differ between clonal lineages 2, 3 and 4 and recombining lineage 1. The protection of locally advantageous allelic combinations is a more plausible explanation for the loss of sex in lineages 2, 3 and 4, because, as shown here, these lineages are found in, and adapted to, different hosts and environmental conditions. This leaves open the question of the maintenance of sex in lineage 1 open, but it is possible the diversity of cultivated and wild rice diversity of rice is higher in the geographical range of this lineage [42]. It is difficult to draw firm conclusions about the evolutionary causes for selection against sexual reproduction, but the finding of an internationally distributed cluster in lineage 1 sheds light on the possible sequence of proximal events underlying the emergence of clonal lineages in this pathogen. The *International* cluster of lineage 1 displayed genetic footprints of recombination and sexual reproduction (in the form of an excess of homoplastic variants or relatively balanced mating types), but the signal of recombination detected here may be purely historical, because we also found that the clonal fraction and the frequency of sterile females were very high in this cluster. The loss of female fertility may, therefore, be an early and major cause of the shift towards asexual reproduction. Some genotypes assigned to the international cluster of lineage 1 may, in the future, come to resemble other clonal lineages in analyses of genealogical relationships, and form a long branch stemming from the central lineage 1, particularly for genotypes already displaying clonal propagation, such as genotypes 2, 18, 58, 98 and 254. However, according to the principle of competitive exclusion, competition between clonal lineages should only allow hegemonic expansion of the most competitive clonal groups, unless they separate into different ecological niches. The asexual spread of the International cluster in a separate ecological niche may ultimately lead to the fixation of a single mating type, as in clonal lineages 2, 3 and 4. This model could also apply to other fungal pathogens with similar life history strategies.

### Effector repertoires differ between lineages

Effector repertoires play a key role in pathogen specialization, in fungal pathogens in general [18], and *P. oryzae* in particular [20, 53, 77-79]. Previous work suggested that clonal lineages of *P. oryzae* possess smaller effector repertoires (Latorre *et al*. 2020) and that the clonal lineage associated with japonica rice (lineage 2) possesses more effectors, and, in particular, Avr-effectors, than other clonal lineages [45, 53]. Our analyzes on a larger set of putative effectors (ca. 2000 in our study, vs. 13 Avr-effectors in [53] and 178 known and candidate effectors in [45]) do not completely confirm this trend as we do not find a significantly larger effector complement in lineage 1 compared to other lineages. Lineage 2 associated with temperate japonica has significantly more putative effectors than lineage 3 associated with indica. However, the number of non-effector genes is also greater in lineage 2 than in lineage 3, and thus it remains possible that the larger effector complement of lineage 2 is a mere consequence of a larger number of genes. We also found that patterns of presence / absence variation mirrored patterns of population subdivision based on SNP allelic variation, and, thus, that the differential sorting of both nucleotide polymorphism and gene content across lineages reflects similar genealogical processes. Our multivariate analyses identified 72 effectors making the greatest contribution to the differentiation of the four lineages in terms of presence / absence. It is possible that the frequency differences for these 72 effectors were due to chance events during bottlenecks at the onset of lineage formation, or, alternatively, that their differential loss was important for the initial adaptation of lineages to new rice populations or subspecies. Interestingly, two of these 72 effectors, AvrRmg8 (=OG0011611) and PWL3 (=OG0011928), are host range determinants in wheat-infecting *P. oryzae* because they trigger immunity in wheat varieties carrying the resistance genes *Rmg8* [80] or *PWT3* [35, 81]. Nucleotide diversity at synonymous and non-synonymous sites also identified 185 effectors with signatures of diversifying selection in one or more lineages, possibly mediated by coevolutionary interactions with host molecules. In the future, it will be interesting to determine the molecular targets of these effectors and to decipher the relationship between their polymorphism and their mode of action.

### Lineage 1 is a threat to global rice production

Our results also indicate that lineage 1 may pose a major threat to rice production. Most of the effectors with signatures of diversifying selection were identified in lineage 1, highlighting the greater ability of this lineage to fix advantageous mutations rapidly. Eleven of the 12 most multivirulent isolates (lesion type > 2 on more than 40 varieties tested) belonged to lineage 1, and the *Yule* cluster, in particular, was highly pathogenic on indica varieties. The propagation of such highly pathogenic genotypes, belonging to a lineage that has both mating types and fertile females, should be monitored closely. This monitoring of lineage 1 is all the more critical because this lineage has an intermediate geoclimatic distribution that overlaps largely with the distributions of the other three lineages, and some of the attempted crosses between clonal lineage 2, 3 and 4 isolates and recombining lineage 1 isolates produced viable progeny. This finding confirms the possibility of gene flow into this lineage, as previously demonstrated on the basis of admixture mapping [45, 46]. It also raises the question of the threat posed by gene flow from the highly pathogenic lineage 1 to the other lineages.

### Concluding remarks

Our study of genetic and phenotypic variation within and between clonal lineages, and within the recombinant lineage of *P. oryzae* suggests a scenario for the emergence of widespread clonal lineages. The loss of female fertility may be a potent driver of the emergence of asexually reproducing groups of clones. The reproductive isolation generated by the loss of sex and the accumulation of mutations due to the absence of sexual purging would facilitate the specialization of some of these clonal groups, leading to the competitive exclusion of the least efficient clonal groups, and, finally, to the propagation of clonal lineages fixed for a single mating type. Our results, thus, demonstrate that the spread of a pathogen across heterogeneous habitats and divergent populations of a crop species can lead to niche separation and reproductive isolation between different invasive lineages in the pathogen.

## MATERIALS AND METHODS

### Biological material

We chose 886 *P. oryzae* isolates collected on Asian rice between 1954 and 2014 as representative of the global genetic diversity of the fungus. Isolates were selected on the basis of microsatellite data, to maximize the number of multilocus genotypes represented ([41] and unpublished data), or based on geographic origin in the absence of genotypic data, to maximize the number of countries represented in the dataset.

Sixty-eight isolates were selected for experimental measurements of reproductive success, adaptation to host, and growth and sporulation experiments at different temperatures (Supplementary file 1). This subset of isolates included 10 isolates from each of the three clonal lineages (lineages 2, 3 and 4), 27 isolates from the various clusters within lineage 1 [*Baoshan* (9 isolates), *International* (10), *Laos* (8), and *Yule* (11)].

Forty-four varieties were chosen as representative of the five main genetic subgroups of Asian rice [82]: indica (Chau, Chiem chanh, DA11, De abril, IR8, JC120, Pappaku), aus (Arc 10177, Baran boro, Black gora, DA8, Dholi boro, Dular, FR13 A, JC148, Jhona 26, Jhona 149, Kalamkati, T1, Tchampa, Tepi boro), temperate japonica (Aichi asahi, Kaw luyoeng, Leung pratew, Maratelli, Nep hoa vang, Nipponbare, Sariceltik, Som Cau 70A), tropical japonica (Azucena, Binulawan, Canella de ferro, Dholi boro, Gogo lempuk, Gotak gatik, Trembese) and aromatic (Arc 10497, Basmati lamo, Dom zard, Firooz, JC1, Kaukkyisaw, N12). The Maratelli (temperate japonica) and CO39 (indica) varieties were used as susceptible controls.

### Genotyping

*Pyricularia oryzae* isolates were genotyped at 5,657 genomic positions with an Illumina Infinium beadchip microarray carrying single-nucleotide polymorphisms identified in 49 previously characterized genomes of rice- and barley-infecting *P. oryzae* isolates [46]. The final dataset included 3,686 biallelic SNPs without missing data.

### Whole genome sequencing

We used a set of 123 whole-genome sequences to investigate differences in effector content between clusters and lineages, including 94 publicly available genomes and 29 new genomes that we selected to increase sample size for the International and Laos clusters of lineage 1, and for lineage 4 (Supplementary file 11). The 29 isolates were sequenced using Illumina HiSeq 3000. Isolates were grown on rice flour-agar medium for mycelium regeneration, then in liquid rice flour medium [83]. Genomic DNA extraction was carried out with >100 mg of fresh mycelium from liquid culture. Fresh mycelium dried on Miracloth paper was crushed in liquid nitrogen. Nucleic acids were subsequently extracted using an extraction buffer (2 % CTAB – 1.4 M NaCl – 0.1 M Tris-HCl pH 8 – 20 mM EDTA pH 8 added before use with a final concentration of 1 % Na_2_SO_3_), then purified with a chloroform:isoamyl alcohol (24:1) treatment, precipitated overnight in isopropanol, and rinsed with 70% ethanol. The extracted nucleic acids were further treated with RNase A (0.2 mg/mL final concentration) and purified with another chloroform:isoamyl alcohol (24:1) treatment followed by overnight precipitation in ethanol. The concentration of extracted genomic DNA was assessed on Qubit^®^ using the dsDNA HS Assay Kit. The purity of the extracted DNA was checked by verifying that the 260/280 and 260/230 absorbance ratios measured with a NanoDrop spectrophotometer were between 1.8 and 2.0. Preparation of TruSeq nano library preparation and HiSeq3000 sequencing (150 nucleotide reads, and 500 bp insert size) were performed at GeT-PlaGe (INRAE, Toulouse, France).

### Genome assembly, gene prediction, orthology analysis, effector prediction, and summary statistics of nucleotide variation

For the 123 sequenced isolates included in the dataset, low-quality reads were removed with Cutadapt software [84]. Reads were assembled with ABYSS 2.2.3 [85, 86], using different K-mer sizes. For each isolate, we chose the assembled sequence with the highest N50 for further analyses. Genes were predicted with Braker 2.1.5 [87, 88] using RNAseq data [55] as extrinsic evidence for model refinement. Genes were also predicted with Augustus 3.4.0 [89](training set=*Magnaporthe grisea*) and gene models that did not overlap with the gene models identified with Braker were added to the GFF file generated with the latter. Repeated regions were masked with RepeatMasker 4.1.0 (http://www.repeatmasker.org/). The completeness of genome assemblies and gene predictions was assessed with BUSCO [90], and completeness in BUSCO genes was greater than 93.7% (Supplementary file 11). Putative effector genes were identified as genes encoding proteins predicted to be secreted by at least two of three methods [SignalP 4.1 [91], TargetP [92] and Phobius [93]], with no predicted transmembrane domain based on TMHMM analysis [94], no predicted motif of retention in the endoplasmic reticulum based on PS-scan [95], and no CAZy annotation based on DBscan v7 [96]. Differences in the numbers of putative effectors and non-effectors between lineages (ANOVA and Tukey’s HSD test) were assessed with the scipy 1.6.0 package in python. Homology relationships betweenpredicted genes were established with OrthoFinder v2.4.0 [97]. Principal component analysis of presence/absence variation for putative effectors and non-effector proteins was performed with the R package prcomp. We used the get_pca_var() in prcomp to extract the results for variables (i.e. effectors) and to identify the effectors making the largest contributions to the principal components, defined as effectors with a contribution to loadings of PC1, PC2 and PC6 greater than 1%. For estimates of the numbers of core and accessory genes, we used a rarefaction approach to account for differences in sample size across lineages. For each pseudo-sample size, we estimated the size of the core and accessory genome in each lineage using a maximum of 2000 pseudosamples. Sequences for each orthogroup were aligned and cleaned with TranslatorX [98], using default parameters. The ratio of non-synonymous to synonymous diversity π_N_/π_S_ was calculated for each orthogroup, for lineages with a sample size of at least four, with Egglib 3 (https://www.egglib.org).

### Population subdivision and recombination

The dataset used for population genetic analysis had no missing data. Clones were therefore identified as isolates with identical genotypes at 3,686 sites. Among the 886 *P. oryzae* isolates genotyped, we identified 264 different multilocus genotypes (i.e., “clones”), which were used for the analysis of population subdivision. We used the SNMF program to infer individual ancestry coefficients in *K* ancestral populations. This program is optimized for the analysis of large datasets and does not assume Hardy-Weinberg equilibrium. It is, therefore, more appropriate to deal with inbred or clonal lineages [99]. We used Splitstree version 4 [100] to visualize relationships between genotypes in a phylogenetic network, with reticulations to represent the conflicting phylogenetic signals caused by homoplasy. PCA was performed as implemented in the Python package Scikit-allel v. 1.3.3. (https://zenodo.org/record/4759368#.YL4R5DaiHwQ). Weir and Cockerham’s F_ST_ was calculated with the R package hierfstat, by the WC84 method [101].

We tested the null hypothesis of clonality using the pairwise homoplasy index (PHI; [50]), maximum χ^2^ (Max χ^2^, [51]) and neighbour similarity score (NSS; [52]), as implemented in the Phipack program (https://www.maths.otago.ac.nz/~dbryant/software.html). Homoplastic sites have sequence identities that are not inherited from a common ancestor, but instead derived from independent events in different branches, such as recurrent mutations, sequencing errors or recombination. Homoplastic sites were identified by concatenating genotypes at all sites, inferring a genealogy with the GTR+GAMMA model in RAxML v.8.2.12 [102], and mapping mutations onto the genealogy with the ‘Trace All Characters’ function of Mesquite under maximum parsimony [103]. In the resulting matrix of ancestral states for all nodes, the number of mutations occurring at each site was counted and sites displaying multiple mutations across the genealogy were considered to be homoplastic. We used PopLDdecay [104] to measure linkage disequilibrium (r^2^) as a function of distance between pairs of SNPs. PopLDdecay was configured with a maximum distance between SNPs of 400 kbp, and a minor allele frequency of 0.005.

### Experimental measurement of reproductive success and female fertility

*Pyricularia oryzae* is a heterothallic fungus with two mating types (*Mat1.1* and *Mat1.2*). Sexual reproduction between strains of opposite mating type can be observed in laboratory conditions, and results in the production of ascospores within female sexual structures called perithecia [105]. Crosses were carried out on a rice flour agar medium (20 g rice flour, 2.5 g yeast extract, 15 g agar and 1 L water, supplemented with 500 000 IU penicillin G after autoclaving for 20 min at 120°C), as described by [105]. We assessed reproductive success by determining the production of perithecia produced after three weeks of culture at 20°C under continuous light. Each cross was performed twice and we determined the mean number of perithecia across repeats. We further assessed the presence of asci and germinating ascospores in perithecia for a subset of crosses, by excising perithecia with a scalpel and counting the number of germinated filaments for each individual ascus after overnight incubation on water agar. The subset of crosses included 10 Mat1.1 isolates (lineage 1: CH0999, CH1065, CH1076; lineage 2: CH0092, MC0016, SP0006; lineage 4: IN0017, IN0092, NP0070, CH0718) and 6 Mat1.2 isolates (lineage 1: CH0997; CH1083, CH1120; lineage 3: BR0019, CH0549, MD0929).

We measured female fertility for 210 isolates (listed in Supplementary file 1), by monitoring perithecium production in crosses involving the tester strains CH0997 (Mat1.2) and CH0999 (Mat1.1). On synthetic media, perithecia are formed at the zone of contact between parental mycelia. Isolates forming perithecia are described as female-fertile. Perithecia can be formed by the two interacting partners or by one partner only. If the two partners are female-fertile, two parallel lines of perithecia are observed at the contact zone between mycelia. If only one partner is female-fertile, only one line of perithecia is observed, at the edge of the mycelium of the fertile isolate.

### Pathogenicity tests

Compatibility between *P. oryzae* isolates and rice plants from the five main genetic subgroups of rice (indica, temperate japonica, tropical japonica, aus and aromatic) was measured in controlled conditions. We inoculated on 46 varieties (see section Biological Material), with 70 isolates. Inoculations were performed as described by [106]. Conidial suspensions (25 000 conidia.mL^-1^) in 0.5% gelatin were sprayed onto three-week-old rice seedlings (> 6 plants/variety). The inoculated plants were incubated for 16 hours in the dark at 27°C and 100% humidity and then for seven days with a day/night alternation (13 hours at 27°C/11 hours at 21°C), before the scoring of symptoms. Lesion type was rated from 1 to 6 [106] and the symptom type was assessed visually on leaves. Each interaction was assessed in three independent experiments. Symptoms scores are ordinal-scale variables (i.e. variables that can be ranked but are not evenly spaced). We therefore analyzed the data with a proportional-odds model with the clm() function implemented in the ordinal version 2018.8-25 in R, including rice type and pathogen lineage as main effects, and the rice type - pathogen lineage interaction. A comprehensive mixed model including all experimental replicates, with isolate as a random effect, was also tested, but did not display convergence. We therefore used the maximum of the three scores in calculations, as this is what most accurately represented the ability of an isolate to exploit a given host genotype. Analyses were also conducted with median values, and gave similar results (not shown). We used an analysis of deviance to evaluate whether effect and interaction terms were statistically significant. Pairwise comparisons of significant factors were performed after computing least-squares means with lsmeans version 2.30-0 in R, with Tukey adjustment. The R commands and their outputs are provided in Supplementary file 9.

### Mycelial growth and sporulation rate at different temperatures

Mycelial growth and sporulation rate were measured for 41 isolates at five different temperatures (10°C, 15°C, 20°C, 25°C and 30°C). For each isolate, Petri dishes containing PDA medium were inoculated with mycelial plugs placed at the center, and incubated in a growth chamber at a fixed temperature. Mycelium diameter was estimated as the mean of two measurements made along two perpendicular axes at different time points. At the end of the experiment, conidia were collected by adding 5 mL of water supplemented with 0.01% Tween 20 to the Petri dish and rubbing the surface of the mycelium. Conidia were counted with a hemocytometer. Three or four independent experiments were performed for each isolate, at each temperature. We had initially planned to carry out only three independent experiments. However, a fourth experiment was eventually performed for some temperature conditions because some isolates did not grow in the first experiment, and because some cultures were invaded by mites and had to be discarded in the second and third experiments.

Mycelium growth was analyzed independently for each temperature with linear mixed-effects models. The variable used in statistical analysis was the square root of the mean mycelial diameter measured along two perpendicular axes on a Petri plate. Post-hoc pairwise comparisons were performed after computing least-squares means with lsmeans version 2.30-0 in R, with Tukey adjustment. The R commands and their outputs are provided in Supplementary file 4.

Sporulation rates were analyzed with a Kruskal-Wallis test with rstatix version 0.7.0 in R. The variable used in statistical analysis was the median number of spores calculated across replicates. Post-hoc pairwise comparisons were performed with Dunn’s non-parametric multiple comparison test. The R commands and their outputs are provided in Supplementary file 7.

### Statistical analyses of climatic data

The outlying mean index (OMI), or marginality, is used to study niche separation and niche breadth. The OMI approach evaluates whether a particular lineage is mostly associated with some climatic conditions or can be sampled with the same probability throughout the range of the species. The approach is based on a multivariate analysis (here a PCA) of climatic data associated with the locations included in the dataset, regardless of the identity of the lineages sampled or the number of isolates sampled. An index of marginality or specialization (the outlying mean index) is then calculated for each lineage represented in the dataset. This index measures the distance between the average habitat of a given lineage and the average habitat conditions of the area studied (corresponding to the distribution of a hypothetical species uniformly distributed under all conditions). Lineages are then placed in environmental conditions that maximize their OMIs, and permutation tests are used to test whether the distribution of a lineage differs from random. OMI gives the same weight to all sampling locations, whether rich or poor in individuals or lineages, and is particularly suitable in cases in which sampling is not homogeneous. Environmental values, consisting of 19 biome values (WorldClim bioclimatic variables [57]), were retrieved for all sampling locations. The 11 temperature variables included in the analysis were (bio1) annual mean temperature, (bio2) mean diurnal range, (bio3) isothermality [100*(bio2/bio7)], (bio4) temperature seasonality [standard deviation*100], (bio5) maximum temperature of the warmest month, (bio6) minimum temperature of the coldest month, (bio7) annual temperature range [bio5-bio6], (bio8) mean temperature of the wettest quarter, (bio9) mean temperature of the driest quarter, (bio10) mean temperature of the warmest quarter, (bio11) mean temperature of the coldest quarter. The eight precipitation variables included in the analysis were (bio12) annual precipitation, (bio13) precipitation of the wettest month, (bio14) precipitation of the driest month, (bio15) precipitation seasonality (coefficient of variation), (bio16) precipitation of the wettest quarter, (bio17) precipitation of the driest quarter, (bio18) precipitation of the warmest quarter, (bio19) precipitation of the coldest quarter. The 11 climatic variables were normalized before analysis. A contingency table was generated to associate the number of isolates from each lineage with each sampling location. Only isolates from our dataset with a precisely known geographic origin (region, city or GPS position) were included in the analysis. A random permutation test with 10,000 permutations was used to assess the statistical significance of marginality for each lineage.

## ACKNOWLEDGMENTS

We thank Tatiana Giraud for useful suggestions. We thank all our colleagues who shared strains or participated in the collection of rice blast samples. This work was supported by Bayer Crop Science Singapore, an ANSES-CIRAD PhD fellowship, CGIAR Research Program on Rice, and Agence Nationale pour la Recherche (ANR) in the framework of the project ANR-18-CE20-0016.

## AUTHOR CONTRIBUTIONS

PG, EF, DT, RI and TK conceived and supervised the research. MT conducted experiments with JM, HT, SCA, SB and VS. MT, PG, FC and SR performed data analyses. PG and MT wrote the manuscript. PG, DT, RI and TK provided funding. All co-authors edited the manuscript.

## DATA AVAILABILITY

All raw sequencing data are deposited under accession codes PRJEB42377 and PRJEB46618.

The following data sets were generated:

1. European Nucleotide Archive M Thierry, F Charriat, J Milazzo, H Adreit, S Ravel, S Cros-Arteil, S Borron, V Sella, T Kroj, R Ioos, E Fournier, D Tharreau, P Gladieux ID PRJEB42377 Whole genome sequencing of Pyricularia oryzae fungi isolated from rice.
2. European Nucleotide Archive M Thierry, F Charriat, J Milazzo, H Adreit, S Ravel, S Cros-Arteil, S Borron, V Sella, T Kroj, R Ioos, E Fournier, D Tharreau, P Gladieux ID PRJEB46618 Whole genome sequencing of Pyricularia oryzae fungi from Benin and Burkina Faso
3. Zenodo M Thierry, F Charriat, J Milazzo, H Adreit, S Ravel, S Cros-Arteil, S Borron, V Sella, T Kroj, R Ioos, E Fournier, D Tharreau, P Gladieux Single-nucleotide polymorphisms, genome assemblies, genome annotations, and gene predictions of Pyricularia oryzae isolates from rice doi 10.5281/zenodo.4561581
4. Zenodo M Thierry, F Charriat, J Milazzo, H Adreit, S Ravel, S Cros-Arteil, S Borron, V Sella, T Kroj, R Ioos, E Fournier, D Tharreau, P Gladieux Single-nucleotide polymorphisms in isolates Pyricularia oryzae isolates from rice and other hosts doi: 10.5281/zenodo.5500197

The following previously published data sets were used:

1. European Nucleotide Archive A Pordel, S Ravel, F Charriat, P Gladieux, S Cros-Arteil, J Milazzo, H Adreit, M Javan-Nikkhah, A Mirzadi-Gohari, A Moumeni, D Tharreau ID PRJEB41186 Origin and evolutionary history of Pyricularia oryzae fungal pathogens infecting maize and barnyard grass in Iran
2. European Nucleotide Archive Z Zhong, M Chen, L Lin, Y Han, J Bao, W Tang, L Lin, Y Lin, R Somai, L Lu, W Zhang, J Chen, Y Hong, X Chen, B Wang, WC Shen, G Lu, J Norvienyeku, DJ Ebbole, Z Wang ID PRJNA354675 Population Genomic Analysis of the Rice Blast Fungus Reveals Specific Events Associated With Expansion of Three Main Clades
3. National Center for Biotechnology Information ID PRJNA320483 Genome-based Molecular Diagnostics for the Blast Fungus
4. National Center for Biotechnology Information ID PRJNA417903 Resequencing of a global collection of *Pyricularia oryzae* isolates

## ADDITIONAL FILES

**Source code 1.** R code used to prepare maps in Figure 6.

**Source code 2.** R code used for analyses based on the Outlying Mean Index, including Principal Component Analysis presented in Figure 6.

**Supplementary file 1.** Isolates of Pyricularia oryzae and analysis of genotyping technical replicates

**Supplementary file 2.** Assignment of genotypes to clusters identified in this study, in Gladieux et al. 2018b (A) and in Saleh et al. 2014 (B and C).

**Supplementary file 3.** Linkage disequilibrium as a function of physical distance.

**Supplementary file 4.** F_ST_ estimates between clusters within lineage 1.

**Supplementary file 5.** Analysis of mycelial growth rates.

**Supplementary file 6.** Mycelial growth rates.

**Supplementary file 7.** Analysis of sporulation rates.

**Supplementary file 8.** Sporulation rates.

**Supplementary file 9.** Matrix of compatibility between four lineages of *P. oryzae* isolates and five subspecies of rice plants, as determined by pathogenicity tests in controlled conditions. Numbers in the matrix represent the maximum symptom score across three replicate experiments. Numbers at the right and bottom margins represent the number of compatible interactions observed (symptom score >2) for varieties and isolates, respectively.

**Supplementary file 10.** Analysis of pathogenicity data.

**Supplementary file 11.** Sequenced isolates.

**Supplementary file 12.** Estimating the size of the core and accessory genome using a rarefaction approach. For each lineage, genomes were resampled in <=2000 combinations of N-1 genomes (N being the sample size). (A) Non-effector genes. (B) Putative effectors.

**Supplementary file 13.** Size of core and accessory genomes for non-effector proteins (A) and putative effectors (B) estimated using a rarefaction approach with a pseudo-sample size of n=30 genomes. Lineage 4 was not included due to small sample size.

**Supplementary file 14.** Summary statistics of non-synonymous and synonymous polymorphism in four lineages of *P. oryzae*. For each lineage, only orthogroups with sample size >=4 were included in calculations. Lineage: lineage of origin of genomes used in calculations for each orthogroup. Orthogroup: orthogroup ID. N: sample size. numNS: number of non-synonymous sites. numS: number of synonymous sites. SNS: number of non-synonymous segregating sites. SS: number of synonymous segregating sites. PiNS: non-synonymous nucleotide diversity. PiS: synonymous nucleotide diversity.

**Supplementary file 15.** ecological niches of the four major lineages (referred to as L1 to L4) considering each of the 19 biomes individually (referred to as Bio1 to Bio19). The x-axis represents the outlying mean index (OMI), which measures the distance between the mean habitat conditions used by a lineage and the mean habitat conditions used by the entire species, to test the hypothesis that different lineages are distributed in regions with different climates.

