## Appendix 1 for "Ecological Differentiation Among Globally Distributed Lineages of the Rice Blast Fungus *Pyricularia oryzae*"

*Pyricularia oryzae* is subdivided into multiple lineages on the basis of preferential association with a single host genus (Gladieux *et al.* 2018a). However, analyses of gene flow have also revealed that genetic exchanges have contributed to the make-up of these lineages, with one rice isolate (87-120) shown to lie outside the group of rice isolates and to harbor blocks of DNA with high levels of identity to the Eleusine- and Eragrostis-associated lineages, indicating recent admixture (Ebbole *et al.* 2021; Gladieux *et al.* 2018a). Here, we present analyses (i) illustrating the magnitude of sequence divergence within and between lineages, (ii) investigating the ability of the 3,686 positions included in our Illumina Infinium beadchip microarray to discriminate between host-associated lineages, (iii) characterizing two rice-infecting isolates from a lineage different from the rice-infecting lineage, but for which inclusion in the dataset had little effect on the conclusions drawn.

The following data sets were generated:
1. European Nucleotide Archive

M Thierry, F Charriat, J Milazzo, H Adreit, S Ravel, S Cros-Arteil, S Borron, V Sella, T Kroj, R Ioos, E Fournier, D Tharreau, P Gladieux

ID PRJEB46618 Whole genome sequencing of Pyricularia oryzae fungi from Benin and Burkina Faso

2. Zenodo

M Thierry, F Charriat, J Milazzo, H Adreit, S Ravel, S Cros-Arteil, S Borron, V Sella, T Kroj, R Ioos, E Fournier, D Tharreau, P Gladieux

Single-nucleotide polymorphisms in isolates Pyricularia oryzae isolates from rice and other hosts

doi: 10.5281/zenodo.5500197

The following previously published data sets were used:

1. European Nucleotide Archive

A Pordel, S Ravel, F Charriat, P Gladieux, S Cros-Arteil, J Milazzo, H Adreit, M Javan-Nikkhah, A Mirzadi-Gohari, A Moumeni, D Tharreau

ID PRJEB41186 Origin and evolutionary history of Pyricularia oryzae fungal pathogens infecting maize and barnyard grass in Iran

2. European Nucleotide Archive

Z Zhong, M Chen, L Lin, Y Han, J Bao, W Tang, L Lin, Y Lin, R Somai, L Lu, W Zhang, J Chen, Y Hong, X Chen, B Wang, WC Shen, G Lu, J Norvienyeku, DJ Ebbole, Z Wang

ID PRJNA354675 Population Genomic Analysis of the Rice Blast Fungus Reveals Specific Events Associated With Expansion of Three Main Clades

3. [National Center for Biotechnology Information](https://www.ncbi.nlm.nih.gov/)

### ID PRJNA320483 Genome-based Molecular Diagnostics for the Blast Fungus

4. National Center for Biotechnology Information

ID PRJNA417903 Resequencing of a global collection of *Pyricularia oryzae* isolates

1. **Population subdivision inferred from polymorphism at single-copy orthologs**

We illustrated the magnitude of sequence divergence between host-associated lineages and within the rice-associated lineage, by reproducing a published analysis (Gladieux *et al.* 2018a) based on the combination of whole-genome data (Table A in Gladieux *et al.* 2018a) and whole-genome data for 123 rice-infecting isolates from this study (Supplementary file 11). The assembled genomic sequences for 49 non-rice isolates from the previous study (Gladieux *et al.* 2018a) were masked and annotated with the same approach as described in the main text for the 123 rice-infecting isolates. Homology analysis with OrthoFinder v2.4.0 identified 5190 single-copy orthologs that were used for inferring population structure. After alignment and cleaning with TranslatorX (Abascal *et al.* 2010), using default parameters, the 5190 single-copy orthologs identified covered 6.1 Mb, and included 650,910 SNPs. Clustering analysis with sNMF (Frichot *et al.* 2014) revealed no recent shared ancestry between isolates collected on rice and isolates collected on other hosts, except for isolate 87-120, which had an ancestry in multiple non-rice lineages (Figure A).

A phylogenetic network based on concatenation of the 650,910 SNPs confirmed that the rice-infecting lineage, with the exception of 87-120, was clearly different from the lineages infecting other hosts (Figure B). The mean number of pairwise differences was 2.23e-4/bp (standard deviation 7.9e-5/bp) within the Rice lineage (excluding isolate 87-120), and ranged from 1.35e-3/bp (standard deviation 5.7e-5/bp) between the Rice and Setaria lineages to 4.38e-3/bp (standard deviation 3.3e-5/bp) between the Rice and Brachiaria2 lineages.

2. **Distinguishing between host-associated lineages of *P. oryzae* with the 3686-positions Infinium beadchip**

Our 3,686-position Illumina Infinium beadchip microarray was originally designed with resequencing data for use exclusively with the rice-associated lineage (Gladieux *et al.* 2018b). We therefore investigated the ability of the set of 3,686 SNPs to recover the phylogenetic relationships demonstrated with single-copy orthologous sequences (Figure B). We mapped sequencing reads for rice and non-rice isolates (Table A; Supplementary file 11) onto the reference genome 70-15 with BWA-mem (Li 2013) and then called and filtered SNPs with Bcftools (Bonfield *et al.* 2021; Danecek *et al.* 2021) (command: bcftools filter -e "INFO/MQ < 50 || INFO/DP < 2000 || INFO/DP > 12000 || QUAL <200" | bcftools view -I -V indels). We extracted the 3,686 positions included in the Infinium beadchip from the filtered VCF with Bedtools intersect (Quinlan 2014) and merged the resulting dataset with our clone-corrected dataset of 264 multilocus genotypes.

The phylogenetic network based on the concatenation of the 3686 polymorphic sites (Figure C) revealed that the Infinium SNPs did not recover the relationships between lineages within *P. oryzae* as they appeared in the network reconstructed from SNPs in single-copy orthologs (Figure B). All non-rice isolates clustered together, and were connected to lineage 1 of rice-infecting isolates by branches that were shorter than the branches connecting lineage 1 and lineages 2 to 4. Isolate 87-120 – which was not genotyped with the Infinium method — was on the same branch as other rice isolates, but was connected to lineage 1 by the longest terminal branch. Only two rice-infecting isolates, BN0019 and BF0072 (International cluster of lineage 1, clonal group 18 [Supplementary file 1], represented by isolate BN0019 in Figure C) branched with non-rice isolates, and can be regarded as spillover pathogens from non-rice lineages or members of a new, previously uncharacterized, lineage of possible hybrid origin.

3. **Characterization of isolates BN0019 and BF0072**

We investigated whether the Infinium-genotyped isolates BN0019 and BF0072 were interlineage hybrids or spillover genotypes, by Illumina-sequencing these two isolates (Supplementary file 11), assembling their genomes with ABySS (Simpson *et al.* 2009), masking repeats, predicting genes and analyzing orthology as described in section 1 of this appendix. Clustering by sNMF and phylogenetic network reconstruction (Figure D and E) indicated that neither of these isolates were spillover pathogens, and that they instead belonged to a distinct lineage potentially resulting from ancient admixture between other lineages. The clonal genotypes BN0019 and BF0072 originated from Benin and Burkina Faso, suggesting that this lineage may represent a perennial, potentially widespread, lineage.

4. **Concluding remarks**

The analysis presented in this appendix shows that our set of Infinium SNPs can identify isolates collected from rice that do not belong to the rice lineage, but cannot distinguish between the various non-rice lineages. By merging whole-genome data from non-rice lineages with our Infinium dataset, we identified two clonal genotypes from Benin and Burkina Faso (isolates BN0019 and BF0072) belonging to a lineage branching between the Rice- and Setaria-infecting lineages. However, the inclusion of these divergent isolates within our dataset is unlikely to have affected the main conclusions of our population genetic analyses with respect to population subdivision and reproductive mode, because these isolates displayed no show fixed differences relative to the rest of isolates from lineage 1 and formed a distinct group in clustering analyses (Figure 2).

Table A. non-rice isolates from Gladieux *et al.* 2018a included in the analysis of population subdivision within *P. oryzae* isolates from multiple hosts. Lineage are named following Gladieux *et al.* 2018a.

| Isolate | Host genus | Lineage |
| --- | --- | --- |
| Br58 | *Avena* | Lolium |
| Bd8401 | *Brachiaria* | Brachiaria1 |
| Bm88324 | *Brachiaria* | Brachiaria2 |
| P28 | *Bromus* | Lolium |
| P29 | *Bromus* | Triticum |
| B51 | *Eleusine* | Eleusine1 |
| Br62 | *Eleusine* | Eleusine1 |
| EI9411 | *Eleusine* | Eleusine2 |
| G22 | *Eleusine* | Eleusine2 |
| PH42 | *Eleusine* | Eleusine1 |
| Z2-1 | *Eleusine* | Eleusine2 |
| G17 | *Eragrostis* | Eragrostis |
| Pg1213-22 | *Festuca* | Lolium |
| TH0016 | *Hordeum* | Oryza |
| CHRF | *Lolium* | Lolium |
| CHW | *Lolium* | Lolium |
| FH | *Lolium* | Lolium |
| HO | *Lolium* | Lolium |
| LpKY97 | *Lolium* | Lolium |
| PgKY | *Lolium* | Lolium |
| PGPA | *Lolium* | Lolium |
| PL2-1 | *Lolium* | Lolium |
| PL3-1 | *Lolium* | Lolium |
| 87-120 | *Oryza* | Unassigned |
| Arcadia | *Setaria* | Setaria |
| GFSI1-7-2 | *Setaria* | Setaria |
| US0071 | *Setaria* | Setaria |
| SSFL02 | *Stenotaphrum* | Stenotaphrum |
| SSFL14-3 | *Stenotaphrum* | Stenotaphrum |
| B2 | *Triticum* | Triticum |
| B71 | *Triticum* | Triticum |
| BdBar | *Triticum* | Triticum |
| BdMeh | *Triticum* | Triticum |
| BR0032 | *Triticum* | Triticum |
| Br130 | *Triticum* | Triticum |
| Br48 | *Triticum* | Triticum |
| Br7 | *Triticum* | Triticum |
| Br80 | *Triticum* | Triticum |
| P3 | *Triticum* | Triticum |
| PY0925 | *Triticum* | Triticum |
| PY36-1 | *Triticum* | Triticum |
| PY5010 | *Triticum* | Lolium |
| PY5033 | *Triticum* | Triticum |
| PY6017 | *Triticum* | Triticum |
| PY6045 | *Triticum* | Triticum |
| PY86-1 | *Triticum* | Lolium |
| T25 | *Triticum* | Triticum |
| WBKY11 | *Triticum* | Lolium |
| WHTQ | *Triticum* | Triticum |

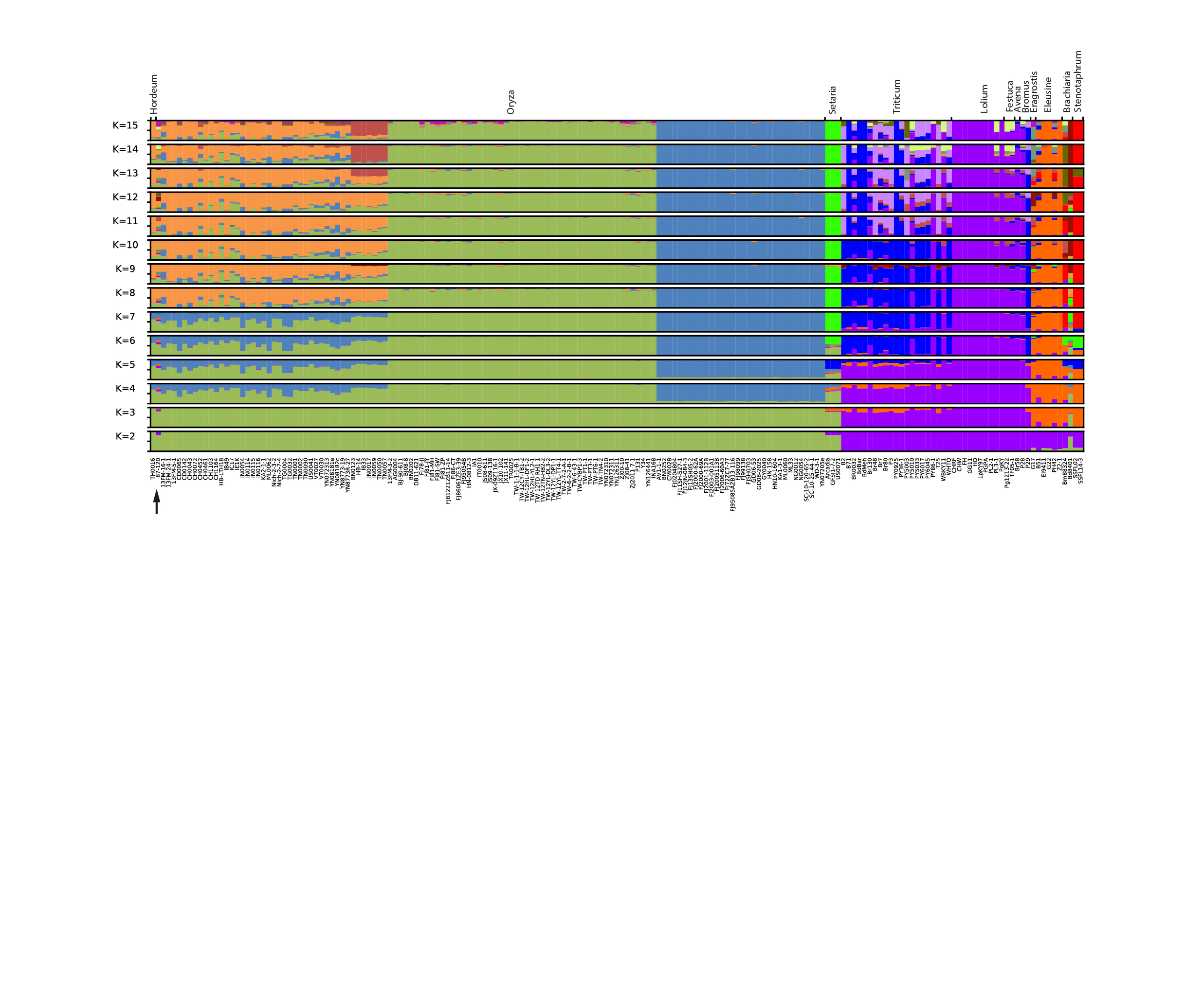

Figure A. Ancestry proportions in *K*=2 to *K*=15 clusters, as estimated with sNMF software for 177 isolates of *P. oryzae* from 11 host genera (Table A, Supplementary file 11), based on 650,910 polymorphisms identified in a set of 5190 single-copy orthologs. Each multilocus genotype is represented by a vertical bar divided into four segments indicating membership in *K* clusters. For each *K*, ten replicate runs were carried out, processed with Clumpak (Kopelman *et al.* 2015), and ancestry coefficients from the major mode are presented. Host of origins of isolates are shown at the top of the figure. Black arrow identifies isolate 87-120, collected on rice, which shares ancestry with non-rice lineages.

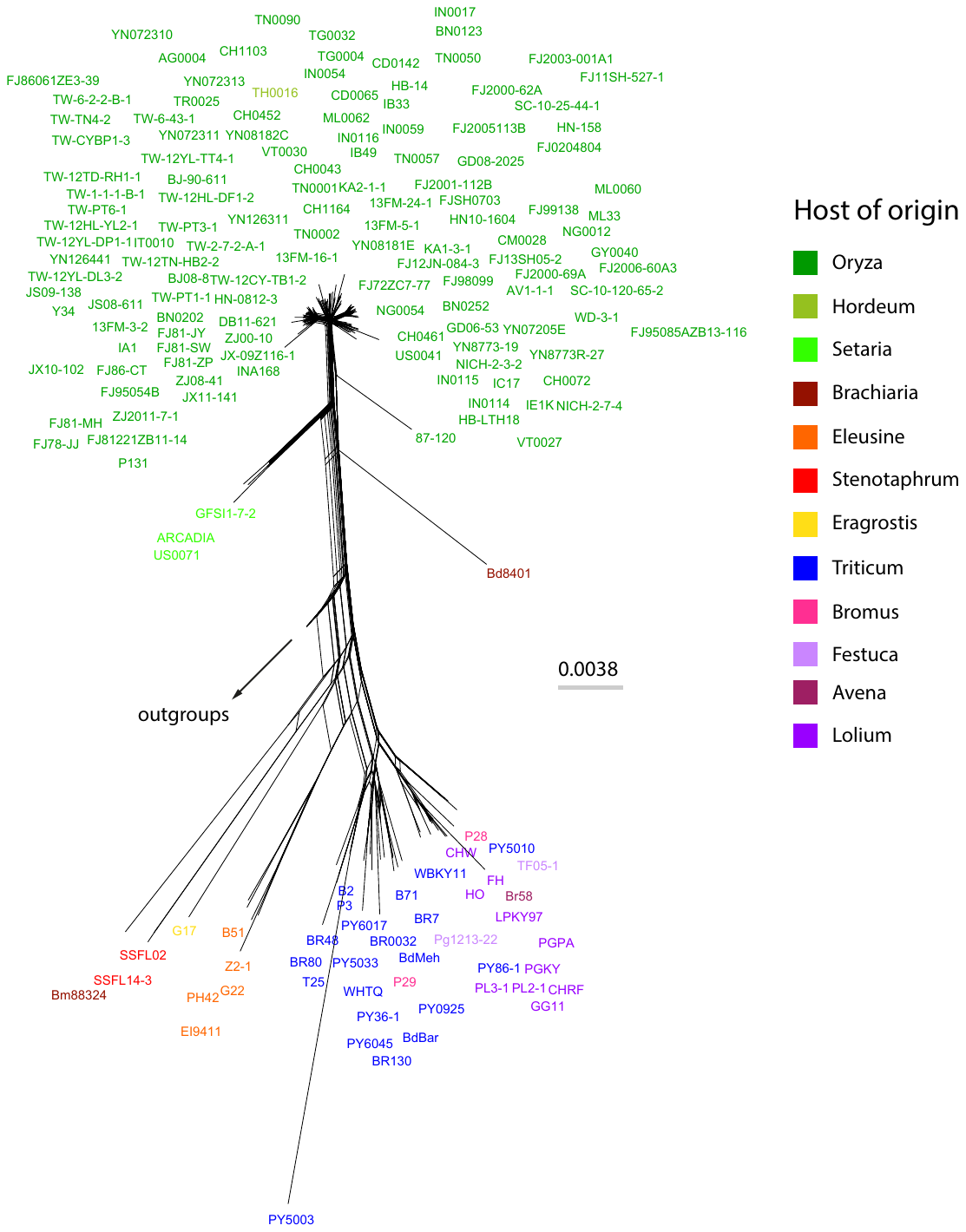

Figure B. Neighbor-net phylogenetic network estimated with Splitstree for 177 isolates of *P. oryzae* from 11 host genera (Table A, Supplementary file 11), based on 650,910 polymorphisms identified in a set of 5190 single-copy orthologs. Isolates BR29 (*P. grisea*) and PM1 (*P. pennisetigena*) were used as outgroups (not shown because connected by a long branch).

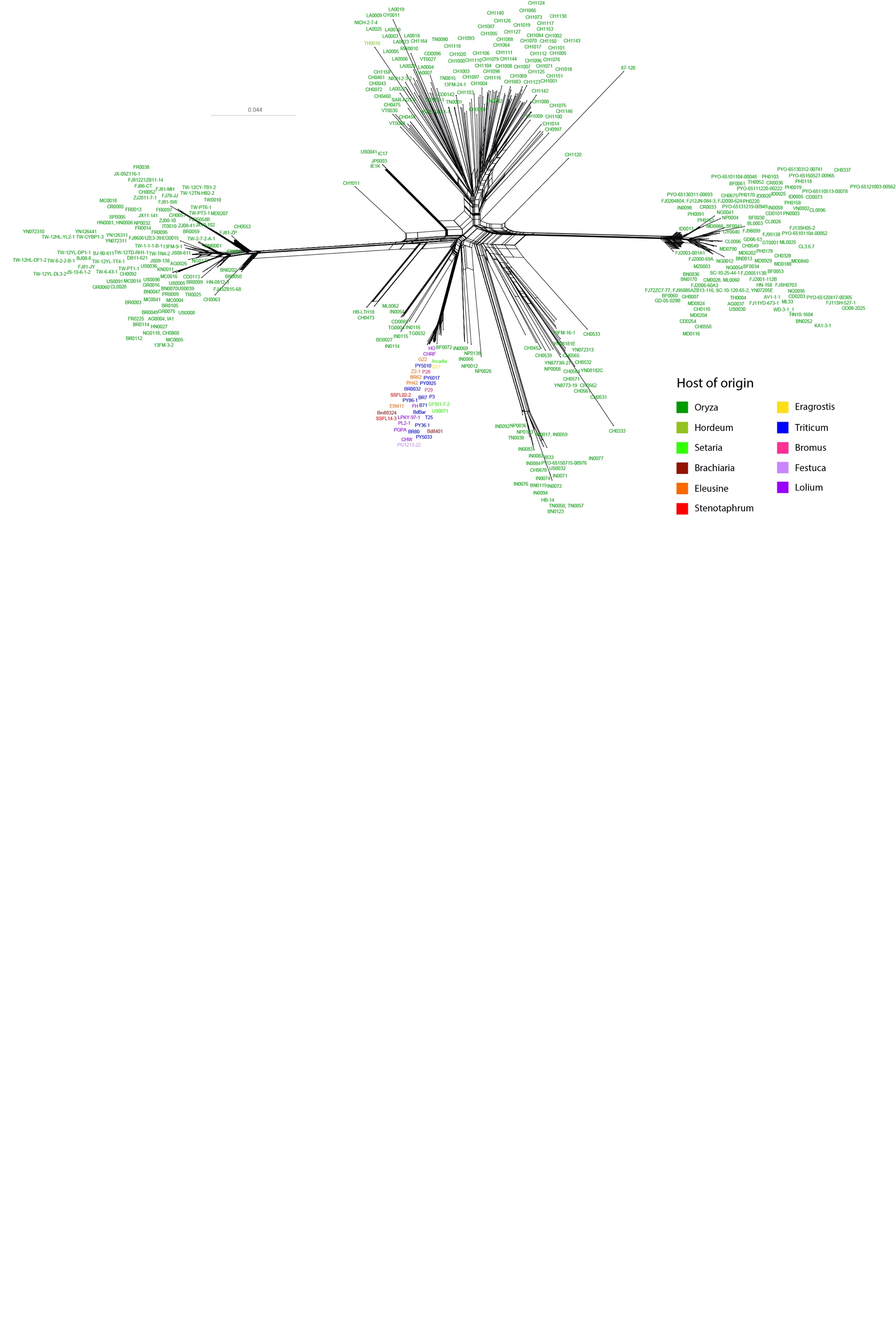

Figure C. Neighbor-net phylogenetic network estimated with Splitstree for 415 isolates of *P. oryzae* from 10 host genera (Table A, Supplementary file 11), based on the concatenation of 3686 single nucleotide polymorphisms. Thirty-eight of the 49 non-rice isolates used in Figure A and B are included in this analysis.

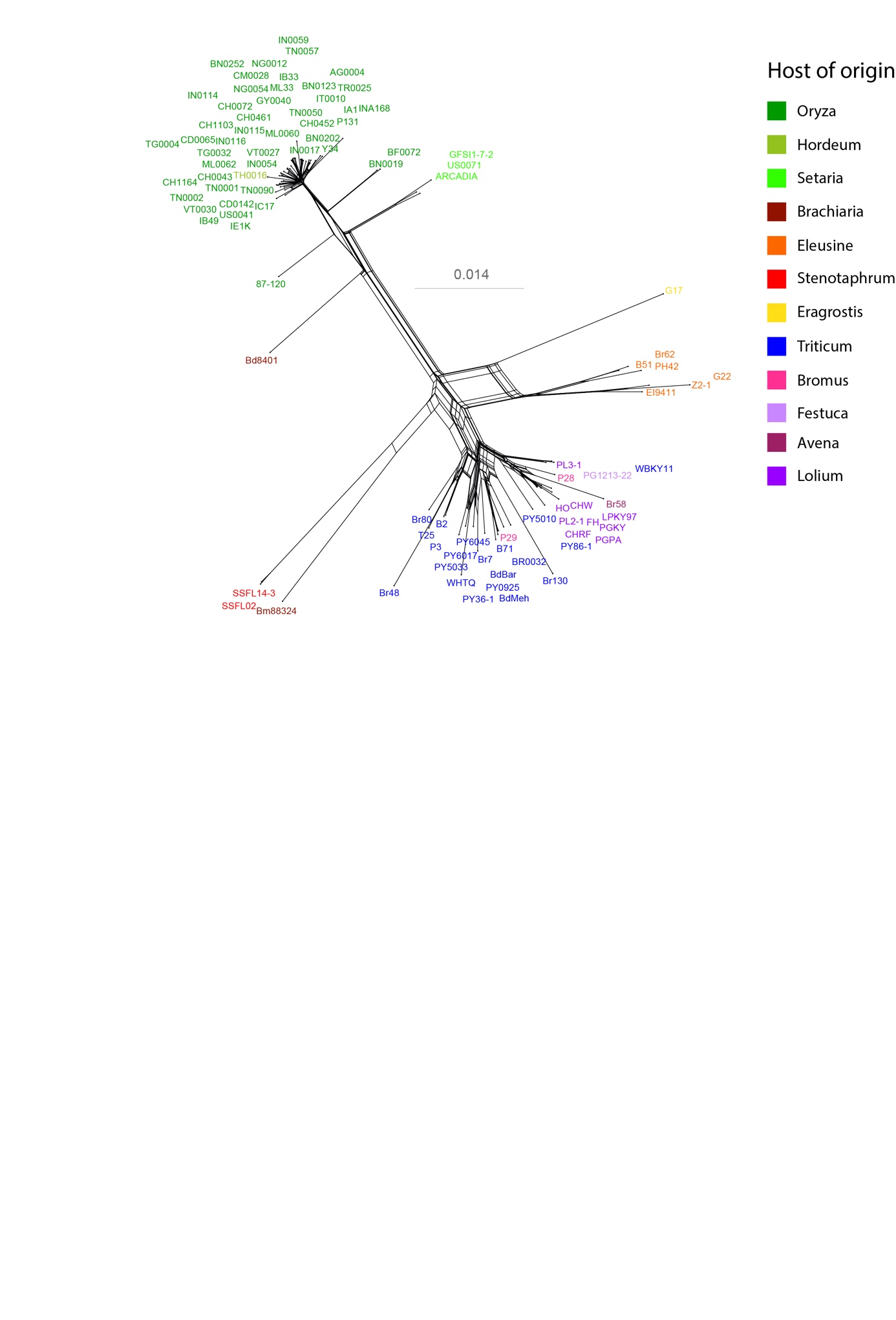

Figure D. Neighbor-net phylogenetic network estimated with Splitstree for 96 isolates of *P. oryzae* from 11 host genera (Table A, Supplementary file 11), based on 503,889 polymorphisms identified in a set of 5075 single-copy orthologs. Fourty-eight of the 123 rice isolates used in Figures A, B and C are included in this analysis.

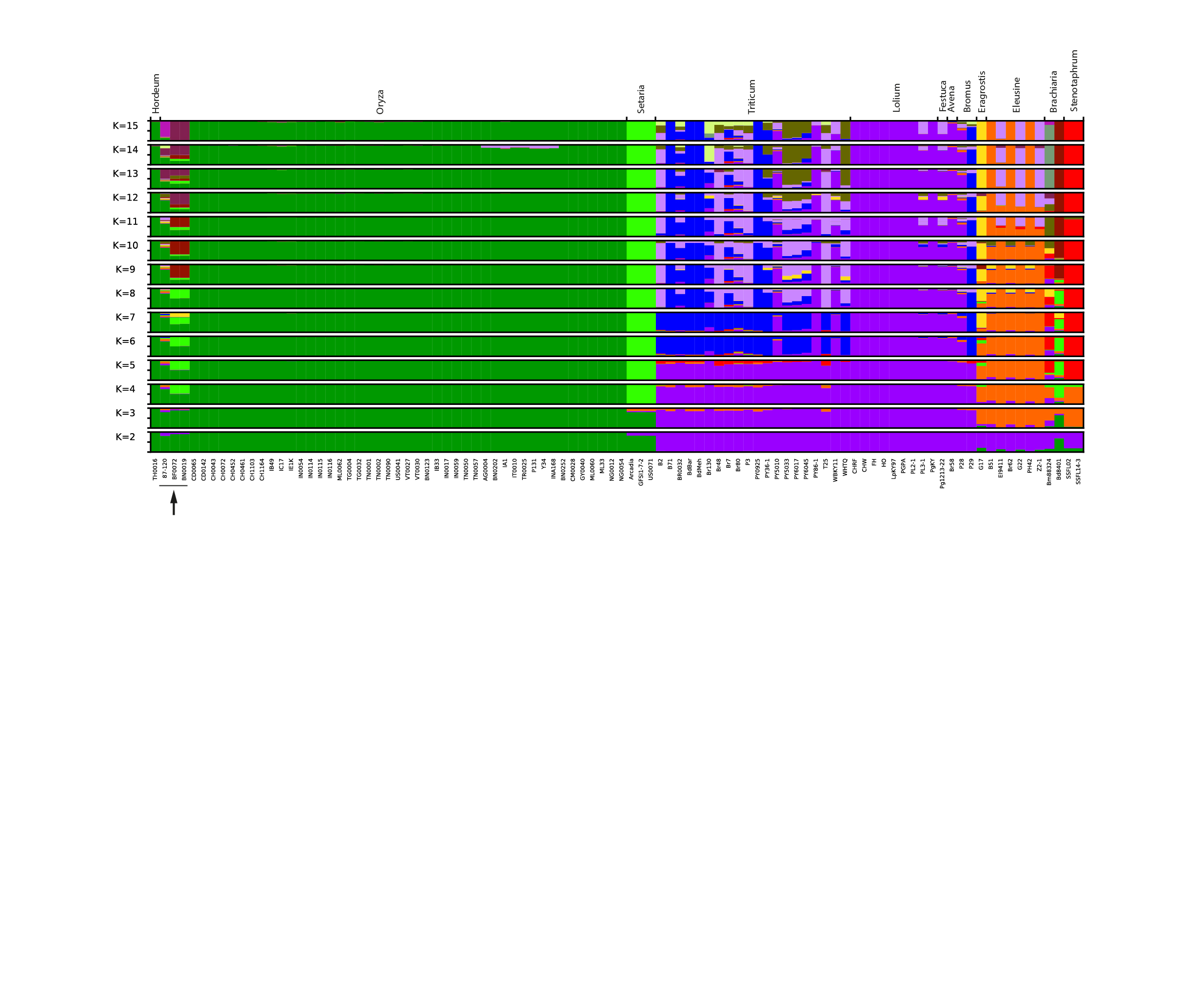

Figure E. Ancestry proportions in *K*=2 to *K*=15 clusters, as estimated with sNMF software for 96 isolates of *P. oryzae* from 11 host genera (Table A, Supplementary file 11), based on 503,889 polymorphisms identified in 5075 single-copy orthologs. Each multilocus genotype is represented by a vertical bar divided into four segments indicating membership in *K* clusters. For each *K*, ten replicate runs were carried out, processed with Clumpak (Kopelman *et al.* 2015), and ancestry coefficients from the major mode are presented. Host of origins of isolates are shown at the top of the figure. Black arrow identifies isolates 87-120, BN0019 and BF0072, collected on rice, which share ancestry with non-rice lineages. Fourty-eight of the 123 rice isolates included in Figures A, B and C are included in this analysis.

Abascal F, Zardoya R, Telford MJ (2010) TranslatorX: multiple alignment of nucleotide sequences guided by amino acid translations. *Nucleic Acids Research* **38**, W7-W13.

Bonfield JK, Marshall J, Danecek P*, et al.* (2021) HTSlib: C library for reading/writing high-throughput sequencing data. LID - 10.1093/gigascience/giab007 [doi] LID - giab007.

Danecek P, Bonfield JK, Liddle J*, et al.* (2021) Twelve years of SAMtools and BCFtools. LID - 10.1093/gigascience/giab008 [doi] LID - giab008.

Ebbole DJ, Chen M, Zhong Z*, et al.* (2021) Evolution and Regulation of a Large Effector Family of Pyricularia oryzae. *Molecular Plant-Microbe Interactions* **34**, 255-269.

Frichot E, Mathieu F, Trouillon T, Bouchard G, François O (2014) Fast and efficient estimation of individual ancestry coefficients. *Genetics* **196**, 973-983.

Gladieux P, Condon B, Ravel S*, et al.* (2018a) Gene Flow between Divergent Cereal- and Grass-Specific Lineages of the Rice Blast Fungus Magnaporthe oryzae. *mBio* **9**.

Gladieux P, Ravel S, Rieux A*, et al.* (2018b) Coexistence of multiple endemic and pandemic lineages of the rice blast pathogen. *mBio* **In Press**.

Kopelman NM, Mayzel J, Jakobsson M, Rosenberg NA, Mayrose I (2015) Clumpak: a program for identifying clustering modes and packaging population structure inferences across K. *Molecular Ecology Resources* **15**, 1179-1191.

Li H (2013) Aligning sequence reads, clone sequences and assembly contigs with BWA-MEM. *arXiv preprint arXiv:1303.3997*.

Quinlan AR (2014) BEDTools: The Swiss-Army Tool for Genome Feature Analysis. *Current protocols in bioinformatics* **47**, 11.12.11-11.12.34.

Simpson JT, Wong K, Jackman SD*, et al.* (2009) ABySS: a parallel assembler for short read sequence data. *Genome Research* **19**, 1117-1123.
