## Appendix 2 for "Ecological Differentiation Among Globally Distributed Lineages of the Rice Blast Fungus *Pyricularia oryzae*"

**Read mapping and SNP calling**

Reads for 123 isolates were aligned against the 70-15 reference genome (version MG8, <https://fungi.ensembl.org/Magnaporthe_oryzae/>) using BWA 0.7.17 and Samtools 1.9. SNP calling was carried out using mpileup in Bcftools 1.9 with option --max-depth 500, and SNPs with mapping quality MQ<=30 or sequencing depth DP<=5 were excluded.

**Assignment of sequenced isolates**

Sequenced isolates were assigned to lineages and clusters identified in *P. oryzae* using genotype data at positions matching the coordinates of the 3,868 SNPs genotyped using our Infinium BeadChip (file Infinium-and-sequencing_SNPs.txt (doi:10.5281/zenodo.4561581)).

Sequenced isolates were assigned to lineages 1-4 by building a neighbor-net network in Splitstree 4 (Huson & Bryant 2006). Identity of the lineages represented in the neighbor-net network was determined using previous assignement work by Latorre et al. (2020) and assignment information for the 33 isolates that were both submitted to Infinium genotyping and whole-genome sequencing (Figure A). Thirty-six isolates were assigned to lineage 1, 48 to lineage 2, 32 to lineage 3, and seven to lineage 4 (Supplementary table 10).

Sequenced isolates assigned to lineage 1 were further assigned to clusters Baoshan, Yule, International and Laos by merging tables of SNPs for sequenced and genotyped isolates from lineage 1 (isolates with genotyping data only: 168; isolates with sequencing data only: 15; isolates with both sequencing and genotyping data: 21; total: 204; file Infinium-and-sequencing_SNPs_lineage1.txt (doi:10.5281/zenodo.4561581)). We used this approach instead of relying on isolates from lineage 1 that were both submitted to Infinium genotyping and whole-genome sequencing because none of these 21 isolates belong to the Baoshan and Yule clusters (Supplementary table 10). Only one representative multilocus genotype was kept for multilocus genotypes repeated multiple times in the dataset (i.e., we “clone-corrected” the dataset; see line “clonal_groups” in file Infinium-and-sequencing_SNPs_lineage1.txt (doi:10.5281/zenodo.4561581)) and population subdivision was inferred using sNMF (Frichot et al., 2014). Multilocus genotypes were assigned to the cluster in which the proportion of ancestry was the highest. Of the 36 isolates from lineage 1 with sequencing data, six were assigned to the Baoshan cluster, 24 to the International cluster, six to the Laos cluster, and none to the Yule cluster (Figure B; Supplementary table 10).

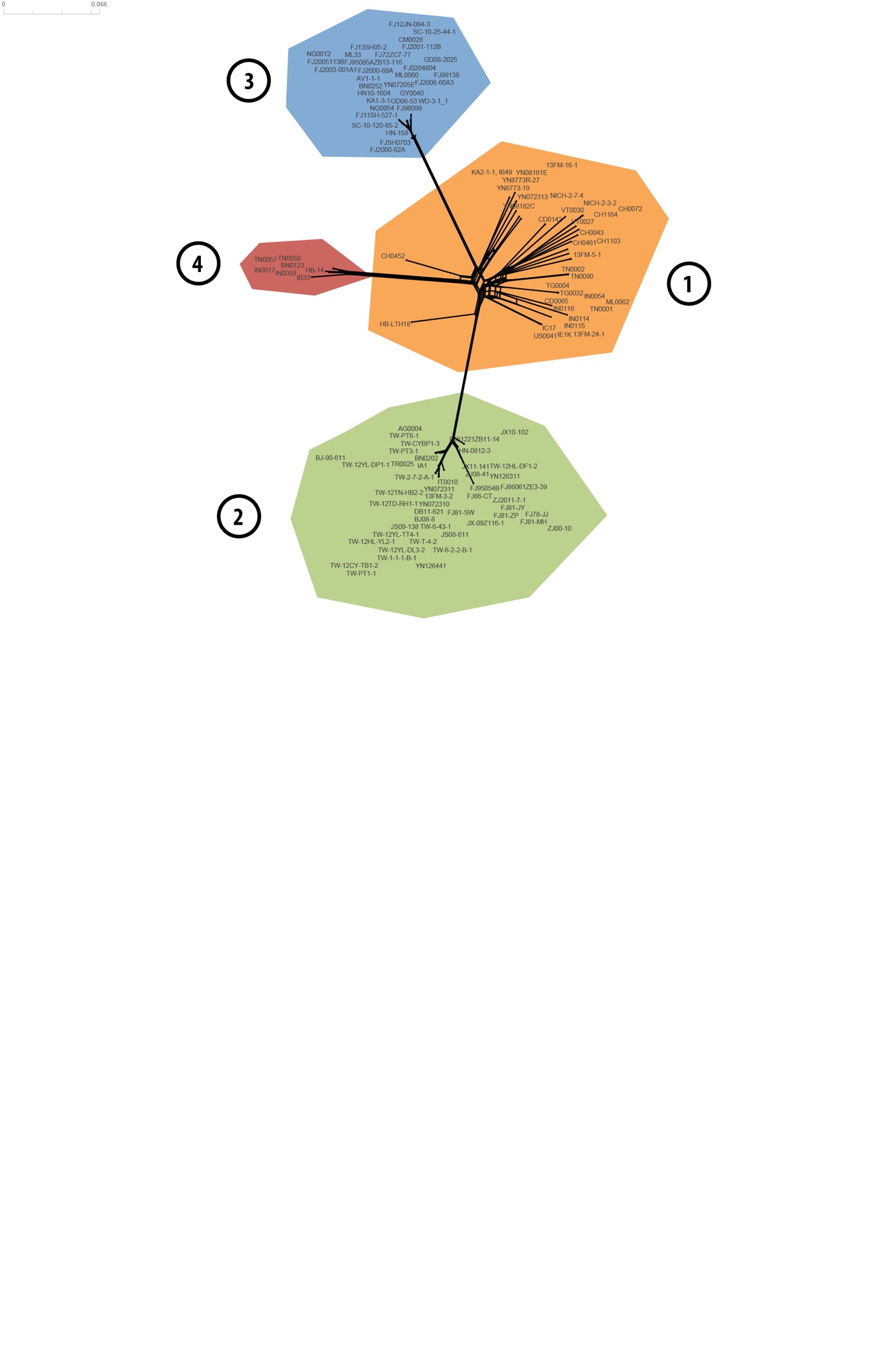

Figure A. Neighbor-net phylogenetic network estimated with Splitstree based on 3,686 SNPs, and showing assignment to four lineages of 123 isolates with whole-genome sequencing data.

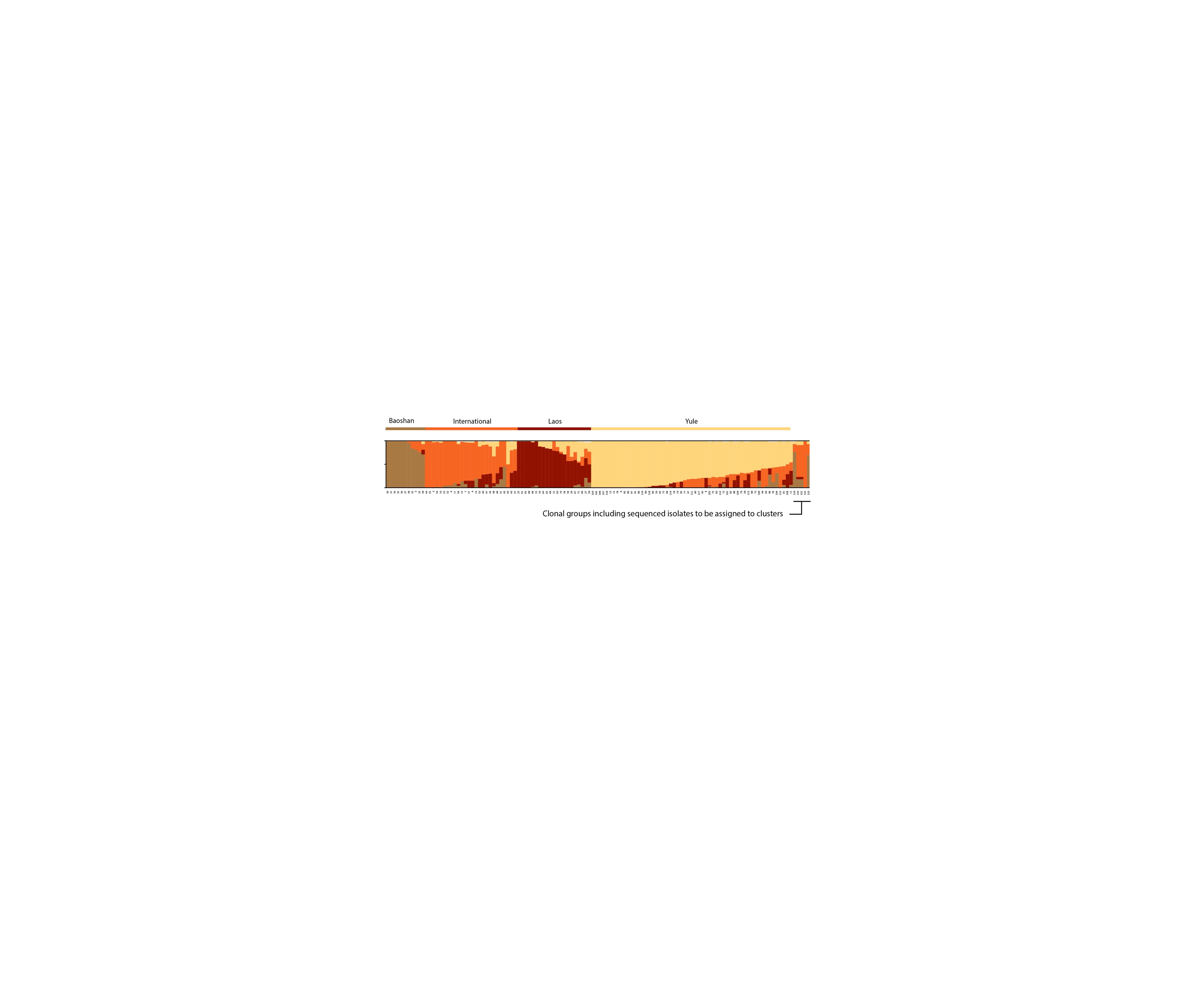

Figure B. Ancestry proportions in four clusters, as estimated with sNMF based on 3,686 SNPs, for 204 isolates with genotyping data, sequencing data, or both. The goal of this analysis was to assign sequenced isolates to clusters. Dataset was clone-corrected and only one representative isolate of each clonal group is represented; hence, it is “clonal groups” that were assigned to clusters, and not strictly speaking “sequenced isolates”. The composition of clonal groups is described in file Infinium-and-sequencing_SNPs.txt (doi:10.5281/zenodo.4561581). Each of the 127 multilocus genotype is represented by a vertical bar divided into four segments, indicating membership in *K*=4 clusters. Cluster names are shown at the top.
