## Supplementary file 3 for "Ecological Differentiation Among Globally Distributed Lineages of the Rice Blast Fungus *Pyricularia oryzae*"

(A)

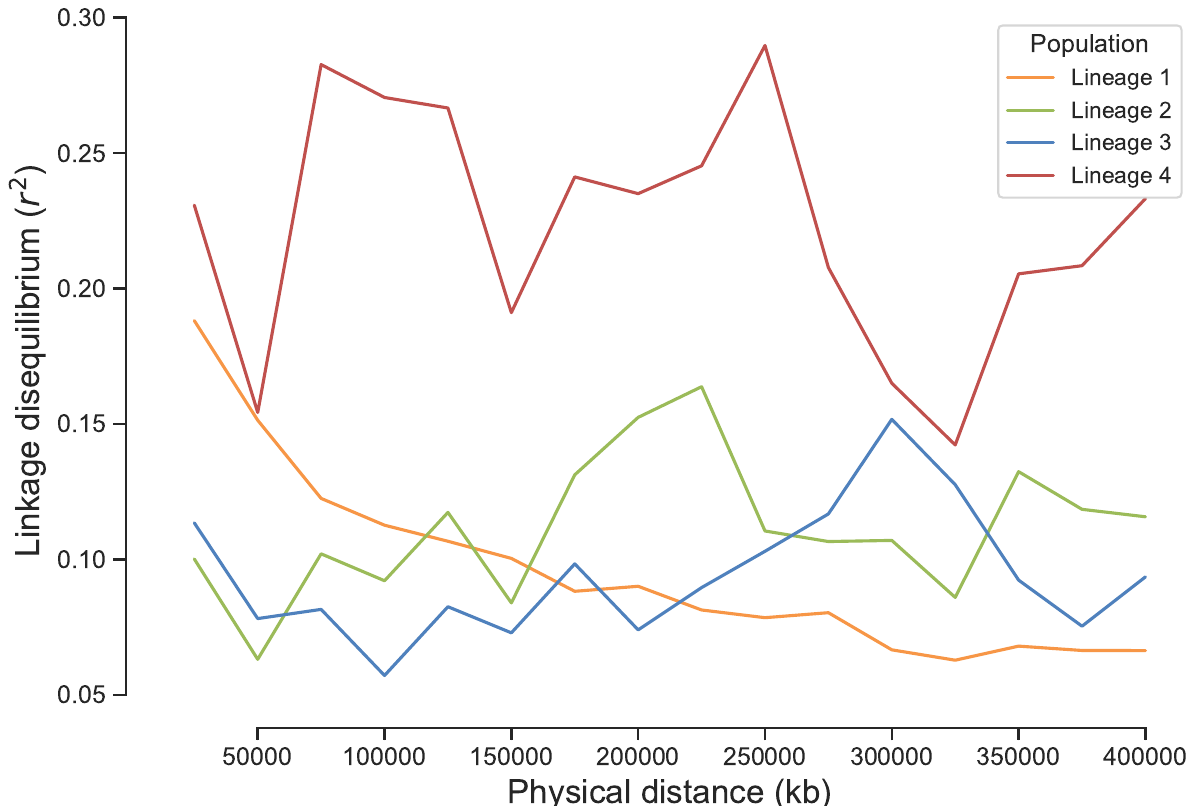

(B)

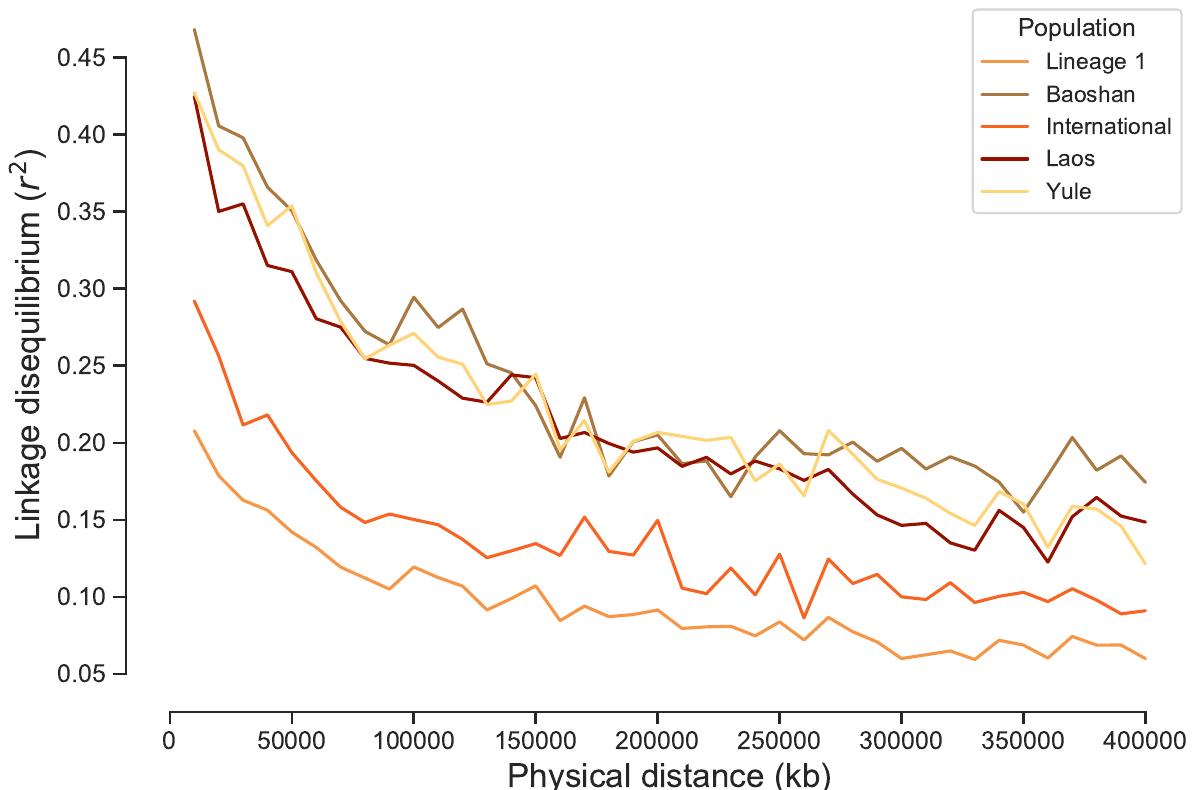

Supplementary file 3. Linkage disequilibrium (*r^2^*) as a function of physical distance (kb) in four lineages of P. oryzae (A) and in clusters within lineage 1 (B).
