## Supplementary file 4 for "Ecological Differentiation Among Globally Distributed Lineages of the Rice Blast Fungus *Pyricularia oryzae*"

Supplementary file 4. F_ST_ estimates between clusters within lineage 1

|  | International | Laos | Baoshan |
| --- | --- | --- | --- |
| Laos | 0.155 |  |  |
| Baoshan | 0.285 | 0.410 |  |
| Yule | 0.219 | 0.180 | 0.487 |
