## Supplementary file 5 for "Ecological Differentiation Among Globally Distributed Lineages of the Rice Blast Fungus *Pyricularia oryzae*"

Supplementary file 5: Analysis of mycelial growth rates

We measured growth rate of 41 representative isolates cultured at different temperatures to test the hypothesis of adaptation to temperature. Main results are shown in Figure SF5-1.

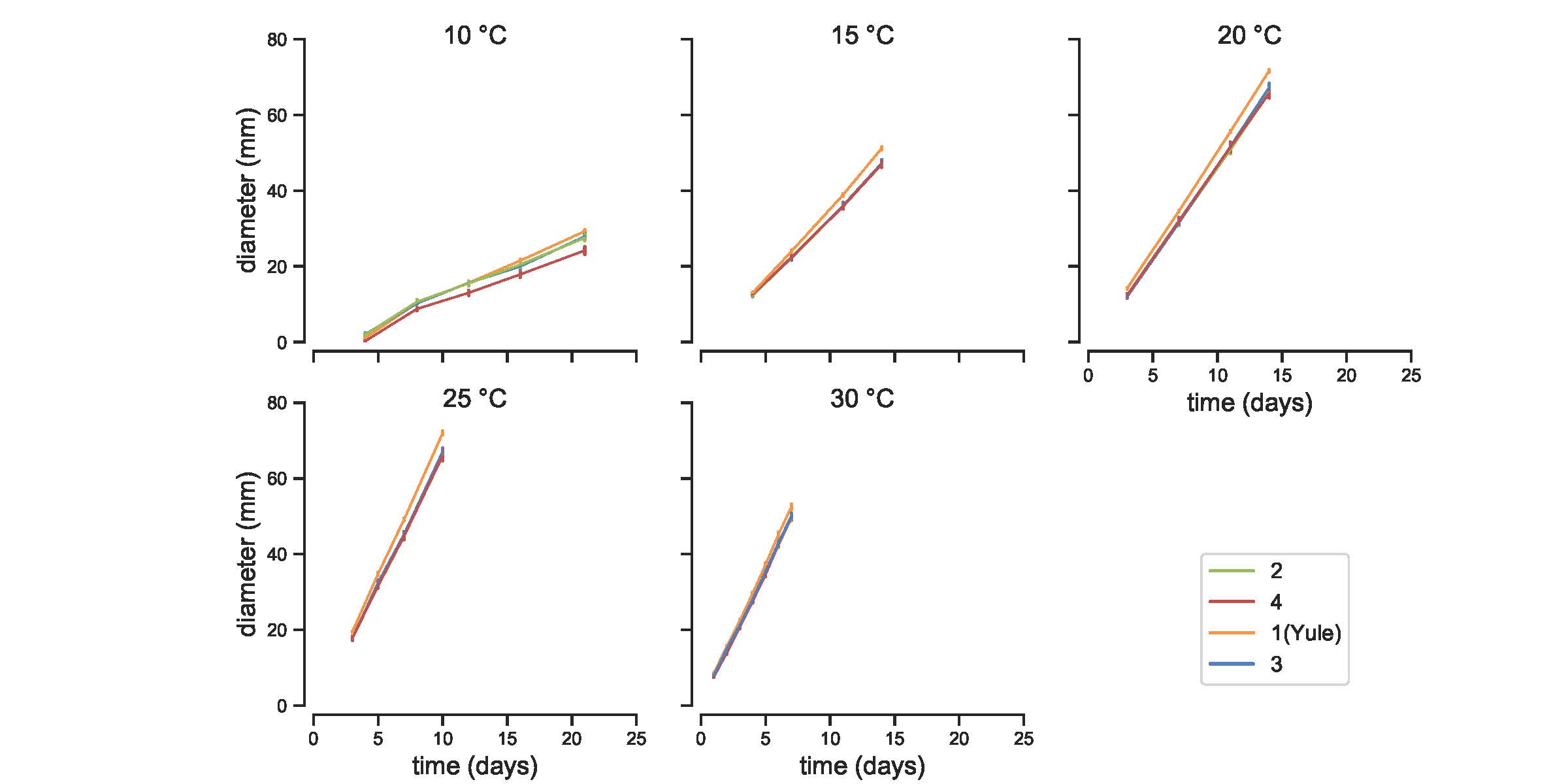

Figure SF5-1. Growth curves for lineages 2-4 and cluster *Yule* within lineage 1 of *P. oryzae* at five incubation temperatures. Error bars represent the standard error across three to four biological replicates.

In what follows, we detail the statistical processing of data:

```
#R packages (R version 4.0.3)
library(lme4)
library(ggplot2)
library(lmerTest)
library(multcomp)
library(lsmeans)
library(RVAideMemoire)
#load and process data
temp_data <- read.table("Supplementary_file_5.txt", sep="\t",header=T)
temp_data$Temperature<-as.factor(temp_data$Temperature)
temp_data$isolate<-as.factor(temp_data$isolate)
temp_data$lineage<-as.factor(temp_data$lineage)
temp_data$replicate<-as.factor(temp_data$replicate)

#T=10°C data summary
T10<- temp_data[ which(temp_data$Temperature==10 & temp_data$diameter>0), ]
T10<-droplevels(T10)

```

#T=10°C plot
ggplot(T10, aes( time,diameter, color=lineage)) +
 stat_summary(fun.data=mean_se, geom="pointrange") + scale_color_manual(values=c("#FFD479", "#9BBB59", "#4F81BD","#C0504D"))

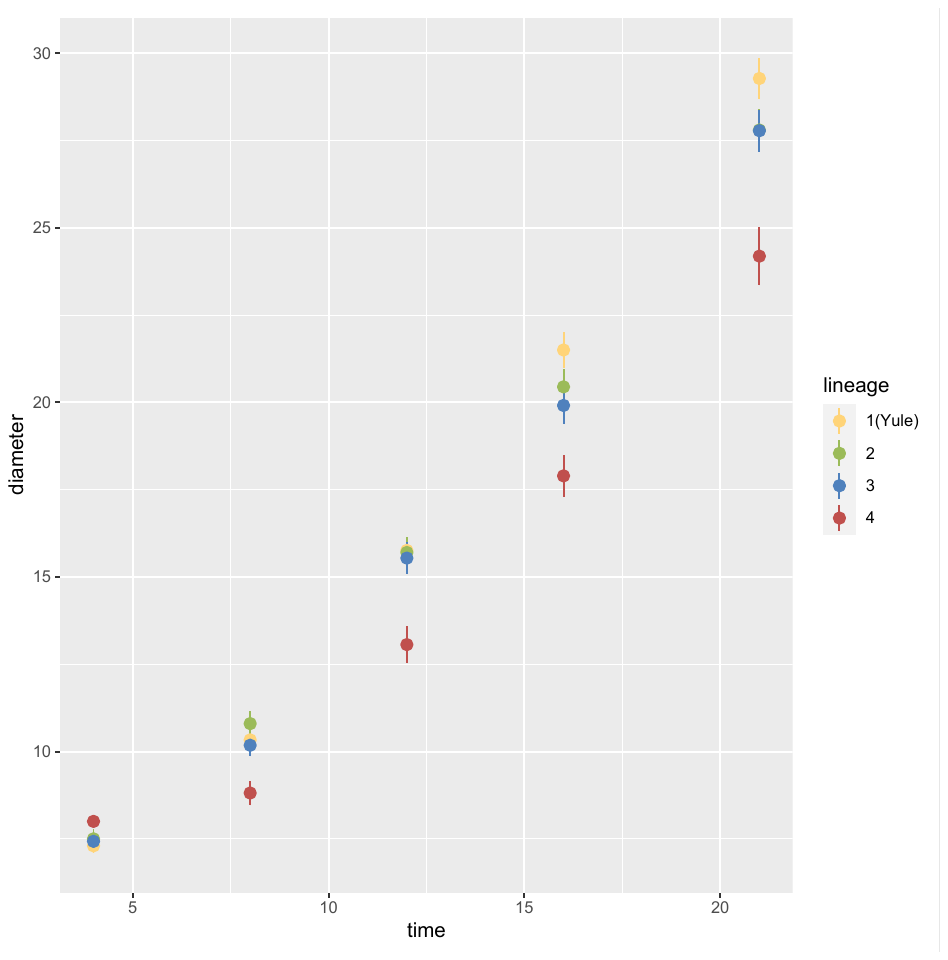

```
#T=10°C: linear mixed model
m.10 <- lmer(sqrt(diameter) ~ time + replicate+ lineage +(1 | isolate), data=T10, REML=T)

#T=10°C: plot residuals
plotresid(m.10)
## Registered S3 methods overwritten by 'car':
##   method                          from
##   influence.merMod                lme4
##   cooks.distance.influence.merMod lme4
##   dfbeta.influence.merMod         lme4
##   dfbetas.influence.merMod        lme4
```

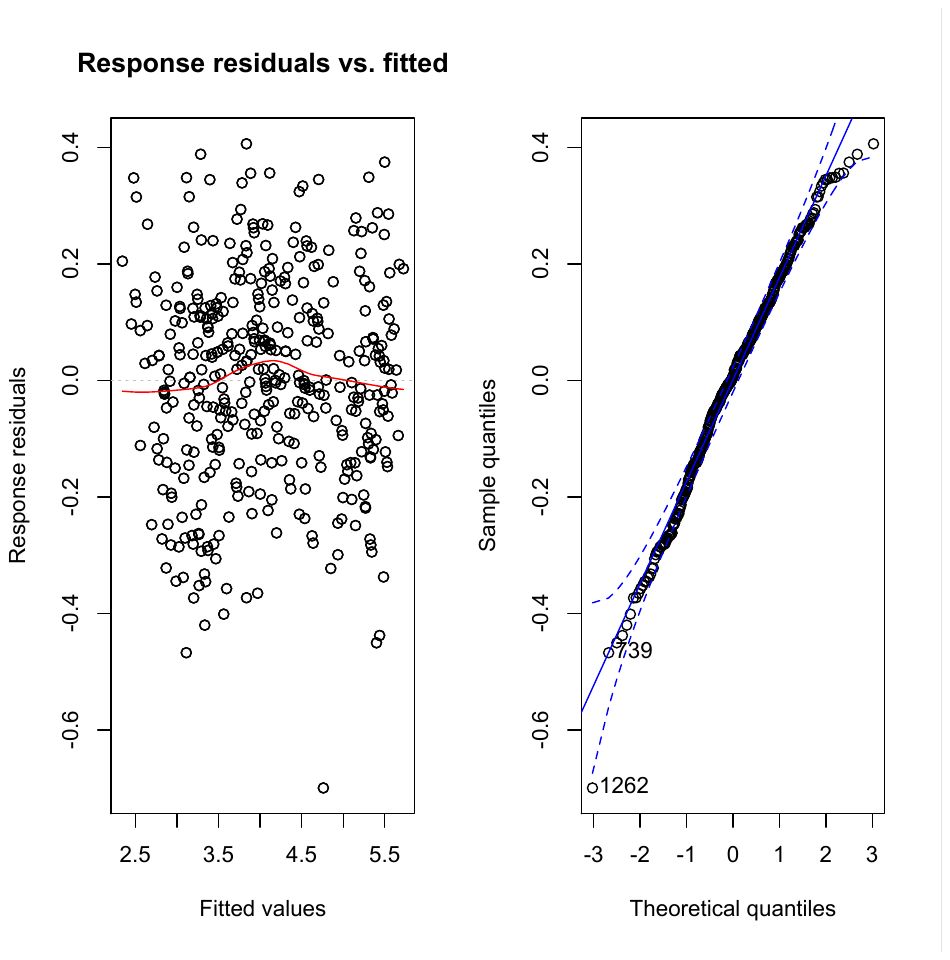

```
#T=10°C: ANOVA
anova( m.10)
## Type III Analysis of Variance Table with Satterthwaite's method
##                   Sum Sq Mean Sq NumDF  DenDF   F value    Pr(>F)
## time 276.203 276.203     1 364.39 7987.5139 < 2.2e-16 ***
## replicate         3.317   1.658     2 367.46   47.9561 < 2.2e-16 ***
## lineage             0.528   0.176     3  35.21    5.0867  0.004972 **
## ---
## Signif. codes:  0 '***' 0.001 '**' 0.01 '*' 0.05 '.' 0.1 ' ' 1
summary(m.10)
## Linear mixed model fit by REML. t-tests use Satterthwaite's method [
## lmerModLmerTest]
## Formula: sqrt(diameter) ~ time + replicate + lineage +
##     (1 | isolate)
##    Data: T10
##
## REML criterion at convergence: -79.3
##
## Scaled residuals:
##     Min      1Q  Median      3Q     Max
## -3.7622 -0.6272  0.0342  0.6412  2.1850
##
## Random effects:
##  Groups   Name        Variance Std.Dev.
##  isolate   (Intercept) 0.04725  0.2174
##  Residual             0.03458  0.1860
## Number of obs: 406, groups:  isolate, 39
##
## Fixed effects:
##                    Estimate Std. Error         df t value Pr(>|t|)
## (Intercept)        2.079740   0.073437  48.001647  28.320  < 2e-16 ***
## time   0.158814   0.001777 364.390128  89.373  < 2e-16 ***
## replicate2       -0.166588   0.021601 365.873700  -7.712 1.19e-13 ***
## replicate3        0.049812   0.025184 368.435269   1.978 0.048685 *
## lineage2           -0.056168   0.101040  34.939604  -0.556 0.581824
## lineage3           -0.063920   0.101069  34.978664  -0.632 0.531213
## lineage4           -0.356141   0.098702  35.598575  -3.608 0.000939 ***
## ---
## Signif. codes:  0 '***' 0.001 '**' 0.01 '*' 0.05 '.' 0.1 ' ' 1
##
## Correlation of Fixed Effects:
##             (Intr) tmps__ rpttn2 rpttn3 ligne2 ligne3
## tmps_d_cltr -0.344
## replicate2 -0.150 -0.003
## replicate3 -0.186  0.166  0.445
## lineage2     -0.621  0.003  0.002 -0.004
## lineage3     -0.619  0.002 -0.004 -0.016  0.451
## lineage4     -0.634 -0.003 -0.008  0.013  0.461  0.461
#T=10°C: model fit
ggplot(T10, aes( time,sqrt(diameter), color=lineage)) +
  stat_summary(fun.data=mean_se, geom="pointrange")+
  stat_summary(aes(y=fitted(m.10)),fun=mean,geom="line")
## Warning: Removed 1 rows containing missing values (geom_segment).
```

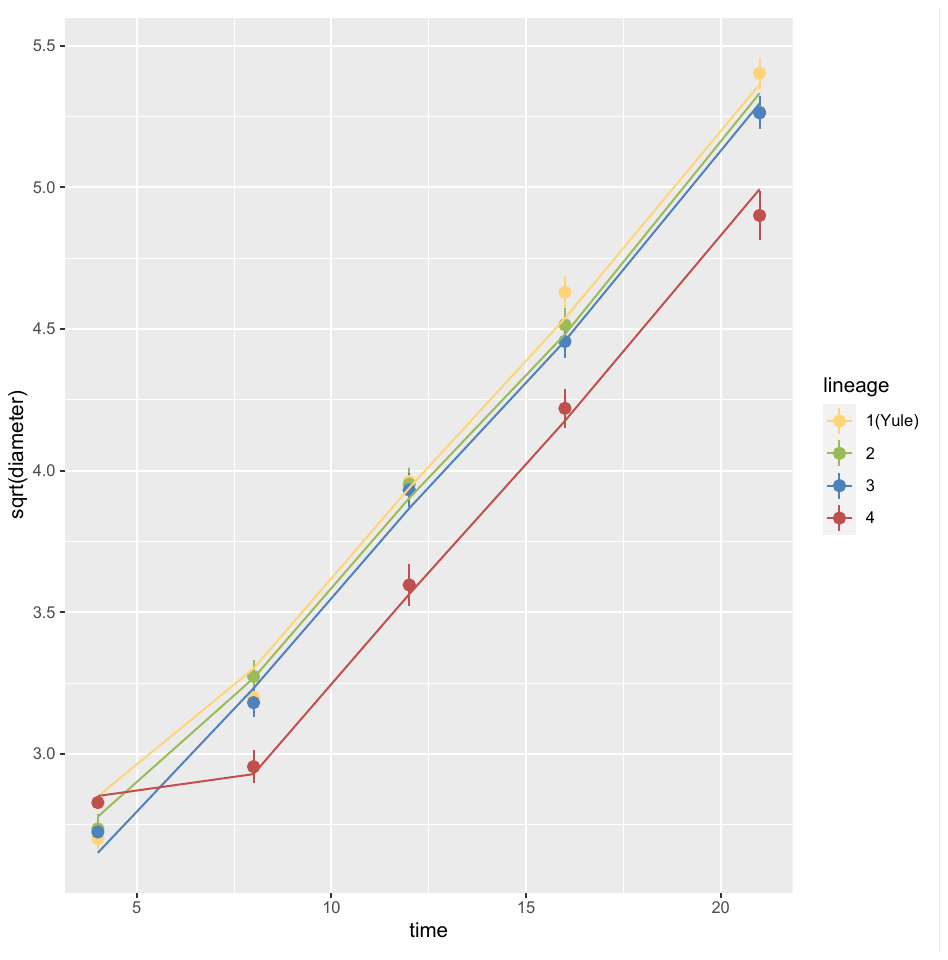

```
#T=10°C : least-squares means
LSM<-lsmeans(m.10,pairwise~lineage)
LSM
## $lsmeans
##  lineage lsmean     SE   df lower.CL upper.CL
##  1(Yule)   4.20 0.0679 34.9     4.06     4.34
##  2        4.15 0.0749 34.6     3.99     4.30
##  3        4.14 0.0749 34.6     3.99     4.29
##  4        3.85 0.0718 36.0     3.70     3.99
##
## Results are averaged over the levels of: replicate
## Degrees-of-freedom method: kenward-roger
## Results are given on the sqrt (not the response) scale.
## Confidence level used: 0.95
##
## $contrasts
##  contrast   estimate     SE   df t.ratio p.value
##  1(Yule) - 2  0.05617 0.1010 34.7 0.556   0.9443
##  1(Yule) - 3  0.06392 0.1011 34.8 0.632   0.9209
##  1(Yule) - 4  0.35614 0.0987 35.4 3.608   0.0050
##  2 - 3       0.00775 0.1059 34.6 0.073   0.9999
##  2 - 4       0.29997 0.1037 35.2 2.894   0.0315
##  3 - 4       0.29222 0.1037 35.2 2.818   0.0377
##
## Results are averaged over the levels of: replicate
## Note: contrasts are still on the sqrt scale
## Degrees-of-freedom method: kenward-roger
## P value adjustment: tukey method for comparing a family of 4 estimates
cld(lsmeans(m.10,~lineage))
## Note: Use 'contrast(regrid(object), ...)' to obtain contrasts of back-transformed estimates
##  lineage lsmean     SE   df lower.CL upper.CL .group
##  4        3.85 0.0718 36.0     3.70     3.99  1
##  3        4.14 0.0749 34.6     3.99     4.29   2
##  2        4.15 0.0749 34.6     3.99     4.30   2
##  1(Yule)   4.20 0.0679 34.9     4.06     4.34   2
##
## Results are averaged over the levels of: replicate
## Degrees-of-freedom method: kenward-roger
## Results are given on the sqrt (not the response) scale.
## Confidence level used: 0.95
## Note: contrasts are still on the sqrt scale
## P value adjustment: tukey method for comparing a family of 4 estimates
## significance level used: alpha = 0.05
#T=15°C data summary
T15<- temp_data[ which(temp_data$Temperature==15 & temp_data$diameter>0), ]
T15<-droplevels(T15)

```

#T=15°C plot
ggplot(T15, aes( time,diameter, color=lineage)) +
 stat_summary(fun.data=mean_se, geom="pointrange") *+ scale_color_manual(values=c("#FFD479", "#9BBB59", "#4F81BD","#C0504D"))*

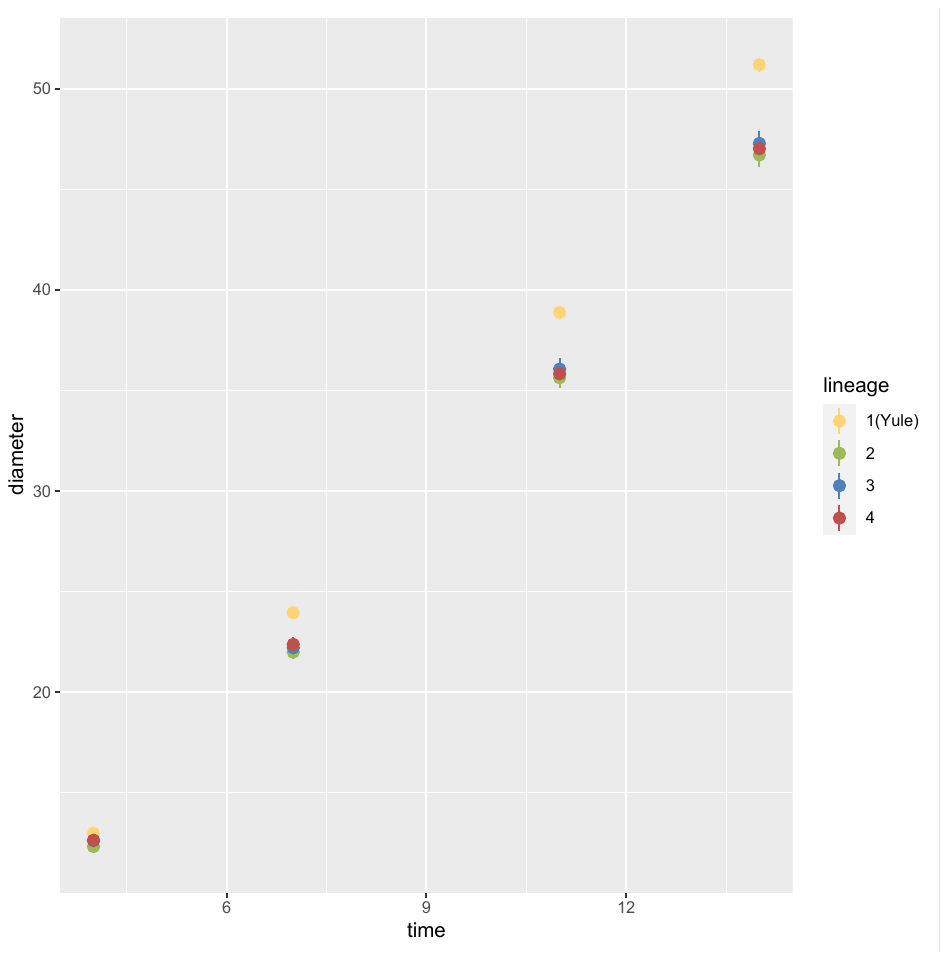

```
#T=15°C: linear mixed model
m.15 <- lmer(diameter ~ time + replicate+ lineage +(1 | isolate), data=T15, REML=T)

#T=15°C: plot residuals
plotresid(m.15)
```

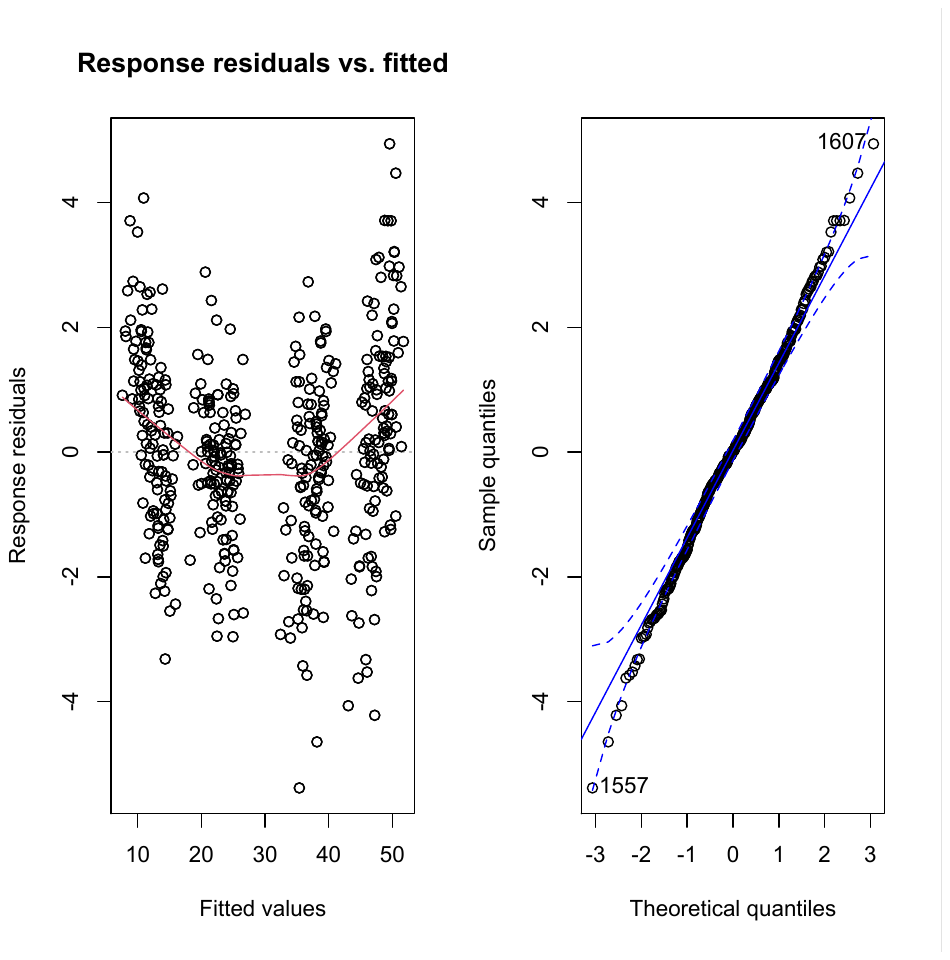

```
#T=15°C: ANOVA
anova( m.15)
## Type III Analysis of Variance Table with Satterthwaite's method
##                  Sum Sq Mean Sq NumDF  DenDF    F value    Pr(>F)
## time  84476   84476     1 418.95 33161.3730 < 2.2e-16 ***
## replicate          117      59     2 420.06    23.0190 3.265e-10 ***
## lineage               35      12     3  35.90     4.6039  0.007944 **
## ---
## Signif. codes:  0 '***' 0.001 '**' 0.01 '*' 0.05 '.' 0.1 ' ' 1
summary(m.15)
## Linear mixed model fit by REML. t-tests use Satterthwaite's method [
## lmerModLmerTest]
## Formula: diameter ~ time + replicate + lineage + (1 |
##     isolate)
##    Data: T15
##
## REML criterion at convergence: 1848.4
##
## Scaled residuals:
##     Min      1Q  Median      3Q     Max
## -3.3730 -0.5755  0.0075  0.6065  3.0952
##
## Random effects:
##  Groups   Name        Variance Std.Dev.
##  isolate   (Intercept) 2.898    1.702
##  Residual             2.547    1.596
## Number of obs: 462, groups:  isolate, 40
##
## Fixed effects:
##                   Estimate Std. Error        df t value Pr(>|t|)
## (Intercept)        0.35650    0.56983  46.78738   0.626  0.53460
## time   3.54757    0.01948 418.95477 182.103  < 2e-16 ***
## replicate2       -1.20323    0.18088 419.58573  -6.652 9.04e-11 ***
## replicate3       -0.37797    0.18394 420.47805  -2.055  0.04051 *
## lineage2           -2.46584    0.79717  36.32295  -3.093  0.00380 **
## lineage3           -2.20758    0.77054  35.53873  -2.865  0.00696 **
## lineage4           -2.28674    0.77054  35.53873  -2.968  0.00534 **
## ---
## Signif. codes:  0 '***' 0.001 '**' 0.01 '*' 0.05 '.' 0.1 ' ' 1
##
## Correlation of Fixed Effects:
##             (Intr) tmps__ rpttn2 rpttn3 ligne2 ligne3
## tmps_d_cltr -0.309
## replicate2 -0.161  0.010
## replicate3 -0.160  0.002  0.485
## lineage2     -0.627  0.004  0.011  0.022
## lineage3     -0.644  0.000  0.000  0.000  0.460
## lineage4     -0.644  0.000  0.000  0.000  0.460  0.476
#T=15°C: model fit
ggplot(T15, aes( time,diameter, color=lineage)) +
  stat_summary(fun.data=mean_se, geom="pointrange")+
  stat_summary(aes(y=fitted(m.15)),fun=mean,geom="line")
```

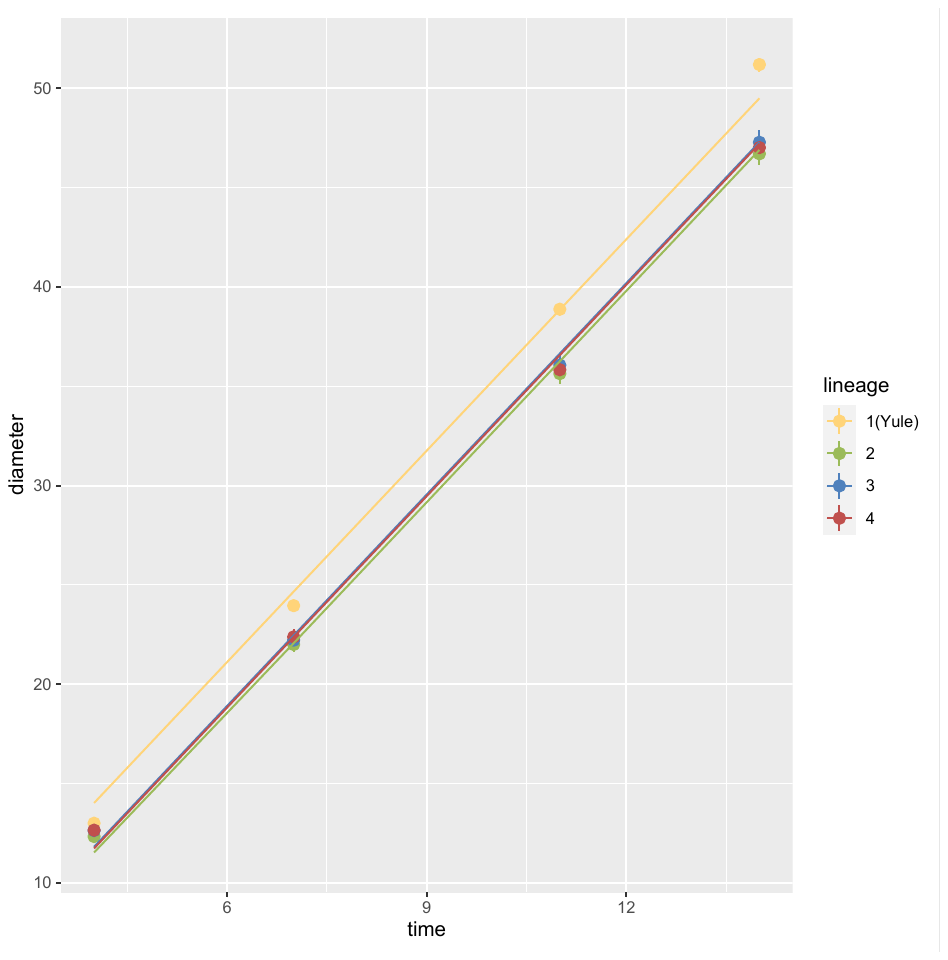

```
#T=15°C: least-squares means
LSM<-lsmeans(m.15,pairwise~lineage)
LSM
## $lsmeans
##  lineage lsmean    SE   df lower.CL upper.CL
##  1(Yule)   31.7 0.532 35.7     30.6     32.8
##  2        29.2 0.594 37.1     28.0     30.4
##  3        29.5 0.558 35.7     28.3     30.6
##  4        29.4 0.558 35.7     28.3     30.5
##
## Results are averaged over the levels of: replicate
## Degrees-of-freedom method: kenward-roger
## Confidence level used: 0.95
##
## $contrasts
##  contrast   estimate    SE   df t.ratio p.value
##  1(Yule) - 2   2.4658 0.797 36.5  3.093  0.0190
##  1(Yule) - 3   2.2076 0.771 35.7  2.865  0.0335
##  1(Yule) - 4   2.2867 0.771 35.7  2.968  0.0262
##  2 - 3       -0.2583 0.815 36.4 -0.317  0.9888
##  2 - 4       -0.1791 0.815 36.4 -0.220  0.9962
##  3 - 4        0.0792 0.789 35.7  0.100  0.9996
##
## Results are averaged over the levels of: replicate
## Degrees-of-freedom method: kenward-roger
## P value adjustment: tukey method for comparing a family of 4 estimates
cld(lsmeans(m.15,~lineage))
##  lineage lsmean    SE   df lower.CL upper.CL .group
##  2        29.2 0.594 37.1     28.0     30.4  1
##  4        29.4 0.558 35.7     28.3     30.5  1
##  3        29.5 0.558 35.7     28.3     30.6  1
##  1(Yule)   31.7 0.532 35.7     30.6     32.8   2
##
## Results are averaged over the levels of: replicate
## Degrees-of-freedom method: kenward-roger
## Confidence level used: 0.95
## P value adjustment: tukey method for comparing a family of 4 estimates
## significance level used: alpha = 0.05
#T=20°C data summary
T20<- temp_data[ which(temp_data$Temperature==20 & temp_data$diameter>0), ]
T20<-droplevels(T20)

```

#T=20°C plot
ggplot(T20, aes( time,diameter, color=lineage)) +
 stat_summary(fun.data=mean_se, geom="pointrange") *+ scale_color_manual(values=c("#FFD479", "#9BBB59", "#4F81BD","#C0504D"))*

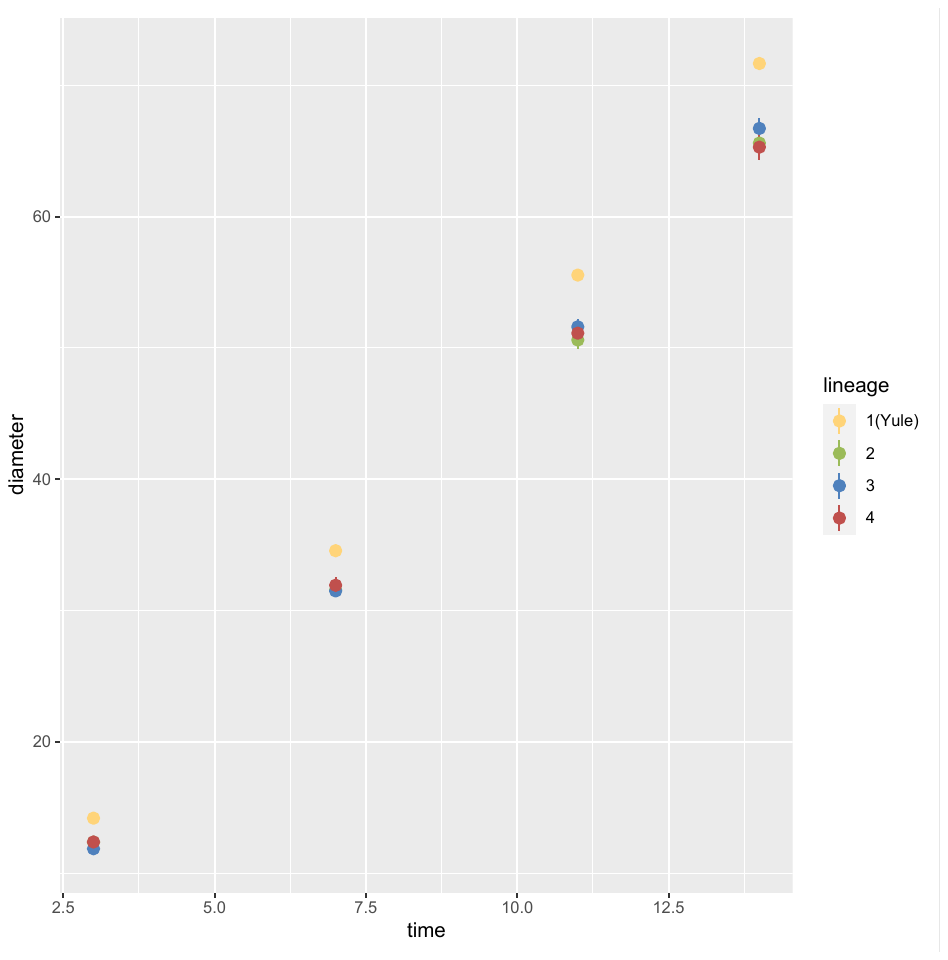

```
#T=20°C: linear mixed model
m.20 <- lmer(diameter ~ time + replicate+ lineage +(1 | isolate), data=T20, REML=T)

#T=20°C: plot residuals
plotresid(m.20)
```

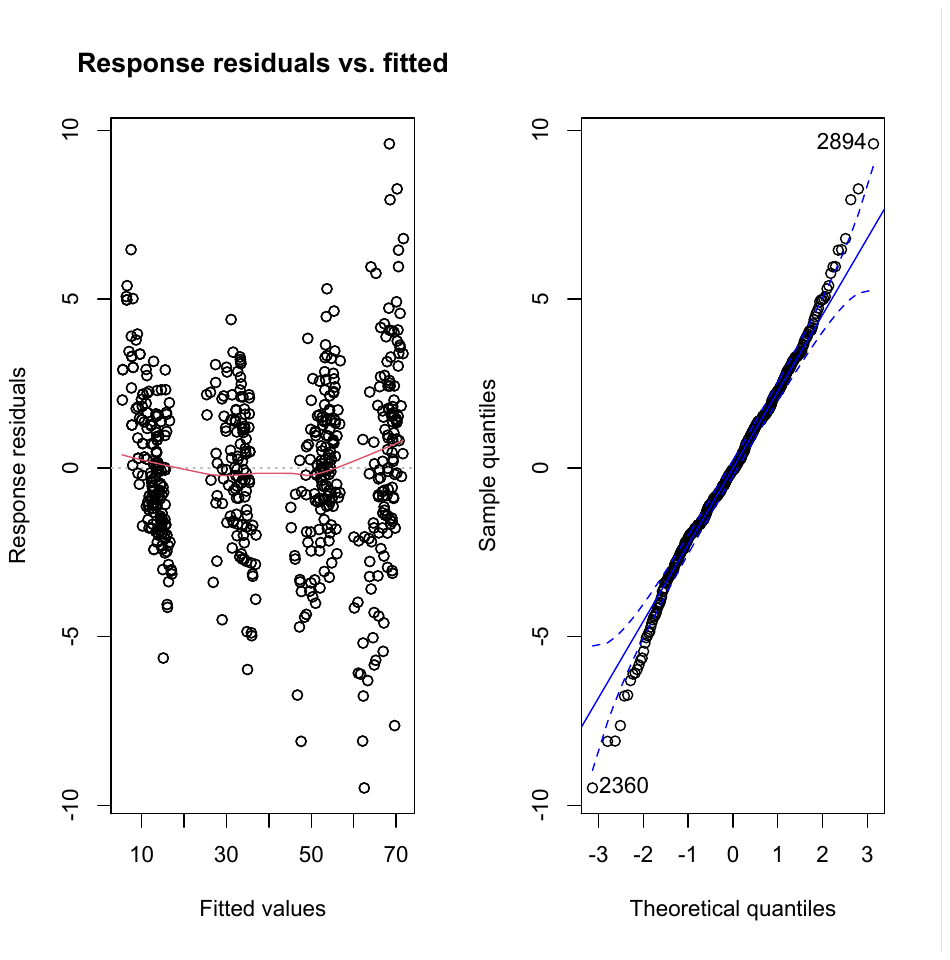

```
#T=20°C: ANOVA
anova( m.20)
## Type III Analysis of Variance Table with Satterthwaite's method
##                  Sum Sq Mean Sq NumDF  DenDF    F value    Pr(>F)
## time 261525  261525     1 542.24 40334.8449 < 2.2e-16 ***
## replicate          262      87     3 544.73    13.4525 1.763e-08 ***
## lineage              131      44     3  37.42     6.7385 0.0009527 ***
## ---
## Signif. codes:  0 '***' 0.001 '**' 0.01 '*' 0.05 '.' 0.1 ' ' 1
summary(m.20)
## Linear mixed model fit by REML. t-tests use Satterthwaite's method [
## lmerModLmerTest]
## Formula: diameter ~ time + replicate + lineage + (1 |
##     isolate)
##    Data: T20
##
## REML criterion at convergence: 2852.5
##
## Scaled residuals:
##     Min      1Q  Median      3Q     Max
## -3.7249 -0.6031 -0.0231  0.5991  3.7700
##
## Random effects:
##  Groups   Name        Variance Std.Dev.
##  isolate   (Intercept) 4.690    2.166
##  Residual             6.484    2.546
## Number of obs: 586, groups:  isolate, 41
##
## Fixed effects:
##                   Estimate Std. Error        df t value Pr(>|t|)
## (Intercept)       -0.02756    0.72532  46.84460  -0.038 0.969846
## time   4.95997    0.02470 542.23625 200.835  < 2e-16 ***
## replicate2       -0.10044    0.29087 543.55006  -0.345 0.730001
## replicate3        1.01102    0.32489 545.15609   3.112 0.001956 **
## replicate4        1.46067    0.28775 546.62482   5.076 5.29e-07 ***
## lineage2           -3.73244    0.98908  36.80540  -3.774 0.000567 ***
## lineage3           -3.51584    0.99726  37.88113  -3.525 0.001124 **
## lineage4           -3.51914    0.99135  37.14063  -3.550 0.001066 **
## ---
## Signif. codes:  0 '***' 0.001 '**' 0.01 '*' 0.05 '.' 0.1 ' ' 1
##
## Correlation of Fixed Effects:
##             (Intr) tmps__ rpttn2 rpttn3 rpttn4 ligne2 ligne3
## tmps_d_cltr -0.243
## replicate2 -0.157 -0.131
## replicate3 -0.138 -0.134  0.444
## replicate4 -0.156 -0.143  0.497  0.455
## lineage2     -0.651  0.001 -0.001 -0.003  0.001
## lineage3     -0.643  0.002 -0.001 -0.010 -0.020  0.473
## lineage4     -0.650  0.006 -0.003  0.003 -0.002  0.476  0.472
#T=20°C: model fit
ggplot(T20, aes( time,diameter, color=lineage)) +
  stat_summary(fun.data=mean_se, geom="pointrange")+
  stat_summary(aes(y=fitted(m.20)),fun=mean,geom="line")
```

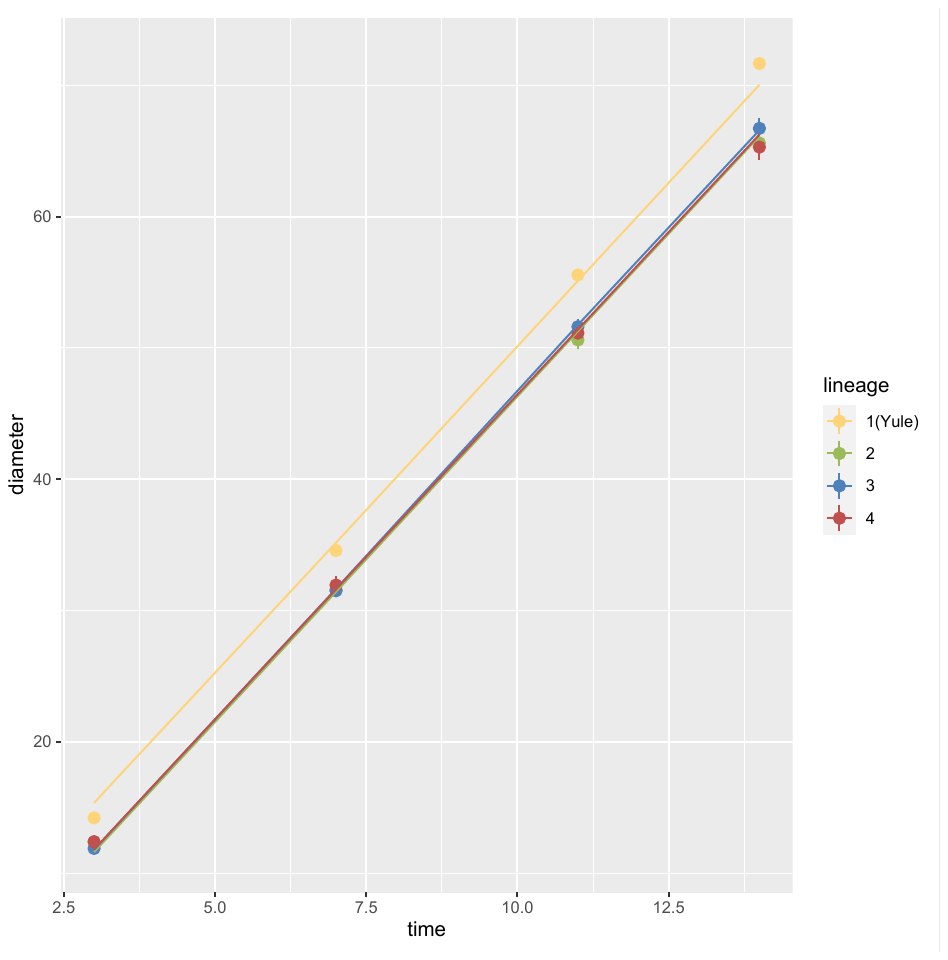

```
#T=20°C: least-squares means
LSM<-lsmeans(m.20,pairwise~lineage)
LSM
## $lsmeans
##  lineage lsmean    SE   df lower.CL upper.CL
##  1(Yule)   41.5 0.683 36.6     40.1     42.8
##  2        37.7 0.715 36.3     36.3     39.2
##  3        37.9 0.727 38.3     36.5     39.4
##  4        37.9 0.719 37.0     36.5     39.4
##
## Results are averaged over the levels of: replicate
## Degrees-of-freedom method: kenward-roger
## Confidence level used: 0.95
##
## $contrasts
##  contrast   estimate    SE   df t.ratio p.value
##  1(Yule) - 2  3.73244 0.989 36.4  3.774  0.0031
##  1(Yule) - 3  3.51584 0.997 37.4  3.525  0.0060
##  1(Yule) - 4  3.51914 0.991 36.7  3.550  0.0057
##  2 - 3      -0.21660 1.020 37.3 -0.212  0.9966
##  2 - 4      -0.21329 1.014 36.6 -0.210  0.9966
##  3 - 4       0.00331 1.022 37.6  0.003  1.0000
##
## Results are averaged over the levels of: replicate
## Degrees-of-freedom method: kenward-roger
## P value adjustment: tukey method for comparing a family of 4 estimates
cld(lsmeans(m.20,~lineage))
##  lineage lsmean    SE   df lower.CL upper.CL .group
##  2        37.7 0.715 36.3     36.3     39.2  1
##  4        37.9 0.719 37.0     36.5     39.4  1
##  3        37.9 0.727 38.3     36.5     39.4  1
##  1(Yule)   41.5 0.683 36.6     40.1     42.8   2
##
## Results are averaged over the levels of: replicate
## Degrees-of-freedom method: kenward-roger
## Confidence level used: 0.95
## P value adjustment: tukey method for comparing a family of 4 estimates
## significance level used: alpha = 0.05
#T=25°C data summary
T25<- temp_data[ which(temp_data$Temperature==25 & temp_data$diameter>0), ]
T25<-droplevels(T25)

```

#T=25°C plot
ggplot(T25, aes( time,diameter, color=lineage)) +
 stat_summary(fun.data=mean_se, geom="pointrange") *+ scale_color_manual(values=c("#FFD479", "#9BBB59", "#4F81BD","#C0504D"))*

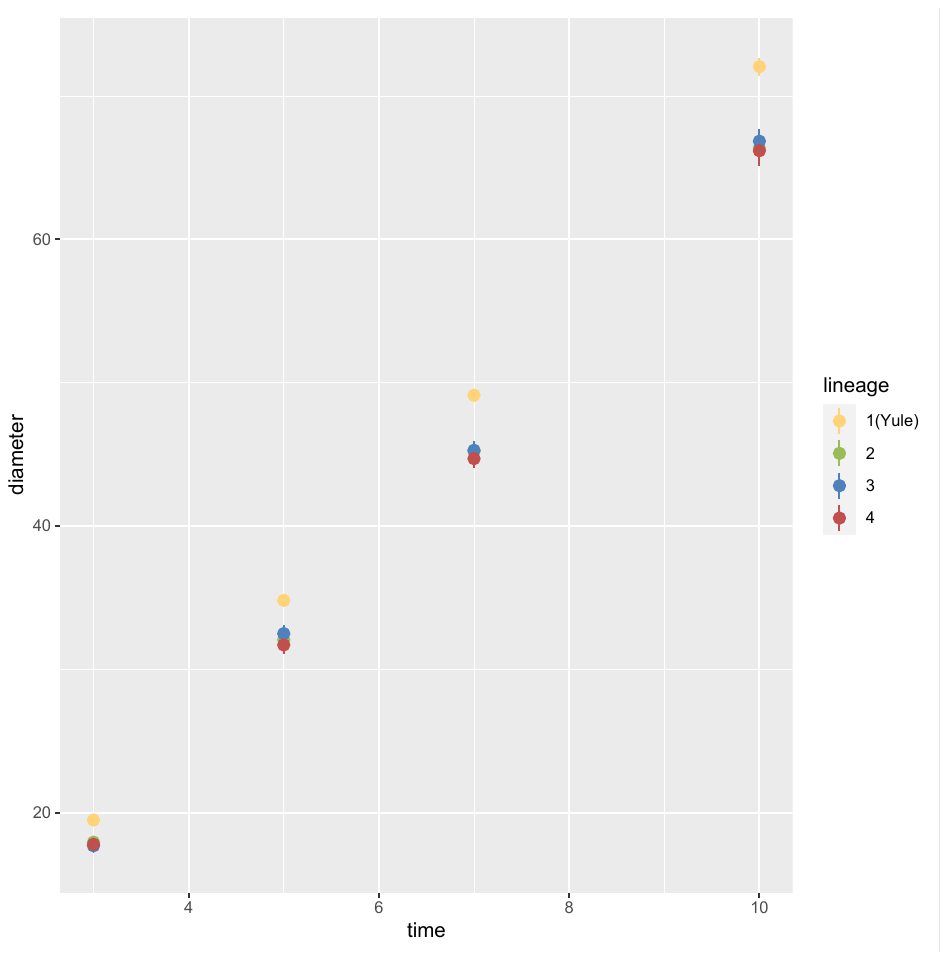

```
#T=25°C: linear mixed model
m.25 <- lmer(diameter ~ time + replicate+ lineage +(1 | isolate), data=T25, REML=T)

#T=25°C: plot residuals
plotresid(m.25)
```

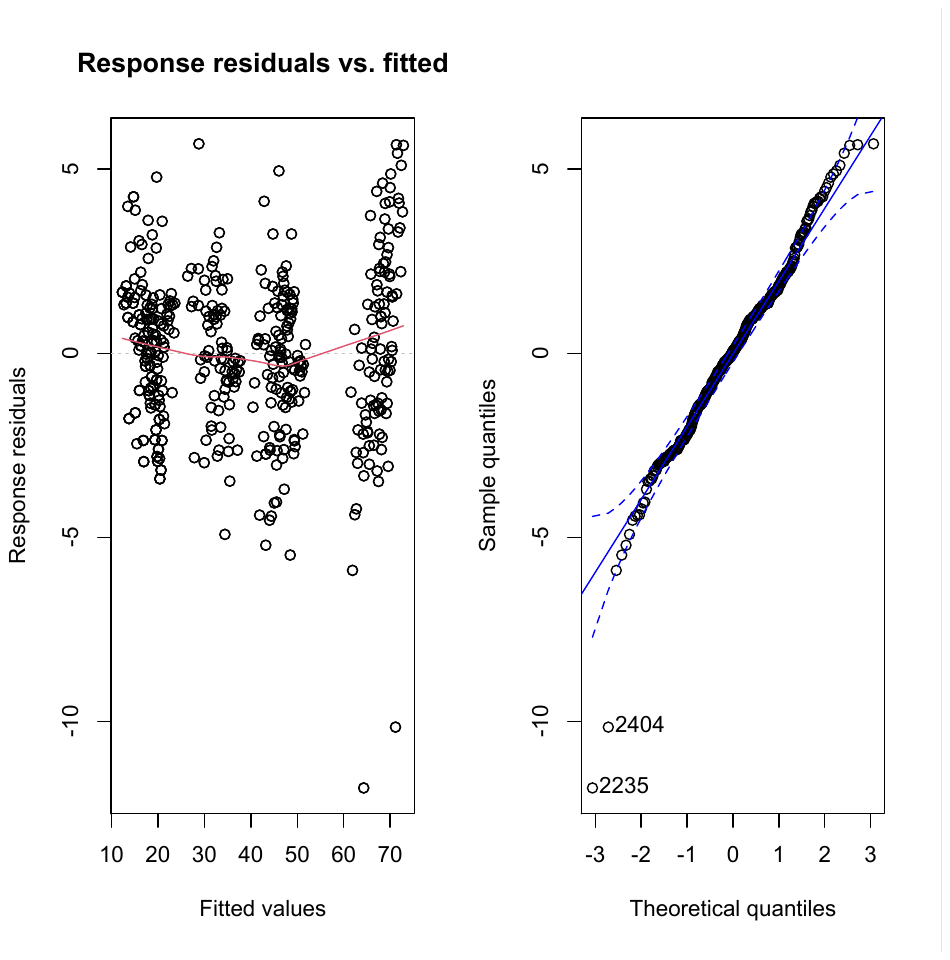

```
anova( m.25)
## Type III Analysis of Variance Table with Satterthwaite's method
##                  Sum Sq Mean Sq NumDF  DenDF    F value    Pr(>F)
## time 151922  151922     1 412.75 30155.7746 < 2.2e-16 ***
## replicate          716     358     2 413.29    71.0545 < 2.2e-16 ***
## lineage              102      34     3  36.19     6.7492 0.0009877 ***
## ---
## Signif. codes:  0 '***' 0.001 '**' 0.01 '*' 0.05 '.' 0.1 ' ' 1
summary(m.25)
## Linear mixed model fit by REML. t-tests use Satterthwaite's method [
## lmerModLmerTest]
## Formula: diameter ~ time + replicate + lineage + (1 |
##     isolate)
##    Data: T25
##
## REML criterion at convergence: 2114.1
##
## Scaled residuals:
##     Min      1Q  Median      3Q     Max
## -5.2596 -0.6017  0.0232  0.5864  2.5340
##
## Random effects:
##  Groups   Name        Variance Std.Dev.
##  isolate   (Intercept) 3.958    1.990
##  Residual             5.038    2.245
## Number of obs: 455, groups:  isolate, 40
##
## Fixed effects:
##                   Estimate Std. Error        df t value Pr(>|t|)
## (Intercept)       -1.33602    0.67969  48.02757  -1.966 0.055136 .
## time   7.03059    0.04049 412.74959 173.654  < 2e-16 ***
## replicate2        0.34003    0.27474 413.62203   1.238 0.216551
## replicate3        2.77275    0.24920 413.49296  11.126  < 2e-16 ***
## lineage2           -3.20583    0.91654  36.09631  -3.498 0.001264 **
## lineage3           -3.14836    0.94356  36.19973  -3.337 0.001970 **
## lineage4           -3.59600    0.91748  36.23012  -3.919 0.000378 ***
## ---
## Signif. codes:  0 '***' 0.001 '**' 0.01 '*' 0.05 '.' 0.1 ' ' 1
##
## Correlation of Fixed Effects:
##             (Intr) tmps__ rpttn2 rpttn3 ligne2 ligne3
## tmps_d_cltr -0.307
## replicate2 -0.091 -0.213
## replicate3 -0.121 -0.164  0.454
## lineage2     -0.643 -0.001  0.003  0.004
## lineage3     -0.625 -0.003  0.010  0.004  0.463
## lineage4     -0.640 -0.006 -0.002 -0.002  0.476  0.462
# model fit
ggplot(T25, aes( time,diameter, color=lineage)) +
  stat_summary(fun.data=mean_se, geom="pointrange")+
  stat_summary(aes(y=fitted(m.25)),fun=mean,geom="line")
```

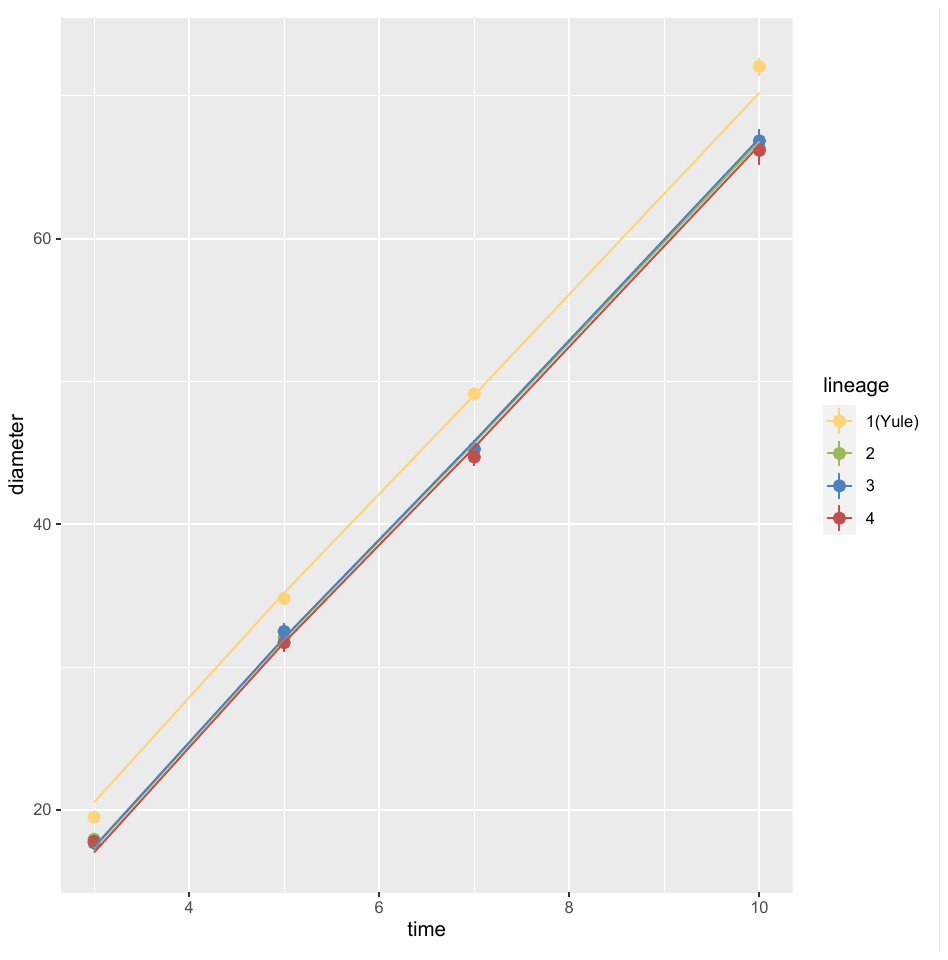

```
#T=25°C: least-squares means
LSM<-lsmeans(m.25,pairwise~lineage)
LSM
## $lsmeans
##  lineage lsmean    SE   df lower.CL upper.CL
##  1(Yule)   41.5 0.633 36.0     40.3     42.8
##  2        38.3 0.663 35.9     37.0     39.7
##  3        38.4 0.700 36.1     37.0     39.8
##  4        37.9 0.665 36.1     36.6     39.3
##
## Results are averaged over the levels of: replicate
## Degrees-of-freedom method: kenward-roger
## Confidence level used: 0.95
##
## $contrasts
##  contrast   estimate    SE   df t.ratio p.value
##  1(Yule) - 2   3.2058 0.917 35.9  3.498  0.0067
##  1(Yule) - 3   3.1484 0.944 36.0  3.337  0.0102
##  1(Yule) - 4   3.5960 0.917 36.0  3.919  0.0021
##  2 - 3       -0.0575 0.964 36.0 -0.060  0.9999
##  2 - 4        0.3902 0.939 36.0  0.416  0.9754
##  3 - 4        0.4476 0.965 36.1  0.464  0.9664
##
## Results are averaged over the levels of: replicate
## Degrees-of-freedom method: kenward-roger
## P value adjustment: tukey method for comparing a family of 4 estimates
cld(lsmeans(m.25,~lineage))
##  lineage lsmean    SE   df lower.CL upper.CL .group
##  4        37.9 0.665 36.1     36.6     39.3  1
##  2        38.3 0.663 35.9     37.0     39.7  1
##  3        38.4 0.700 36.1     37.0     39.8  1
##  1(Yule)   41.5 0.633 36.0     40.3     42.8   2
##
## Results are averaged over the levels of: replicate
## Degrees-of-freedom method: kenward-roger
## Confidence level used: 0.95
## P value adjustment: tukey method for comparing a family of 4 estimates
## significance level used: alpha = 0.05
#T=30°C data summary
T30<- temp_data[ which(temp_data$Temperature==30 & temp_data$diameter>0), ]
T30<-droplevels(T30)

```

#T=30°C plot
ggplot(T30, aes( time,diameter, color=lineage)) +
 stat_summary(fun.data=mean_se, geom="pointrange") *+ scale_color_manual(values=c("#FFD479", "#9BBB59", "#4F81BD","#C0504D"))*

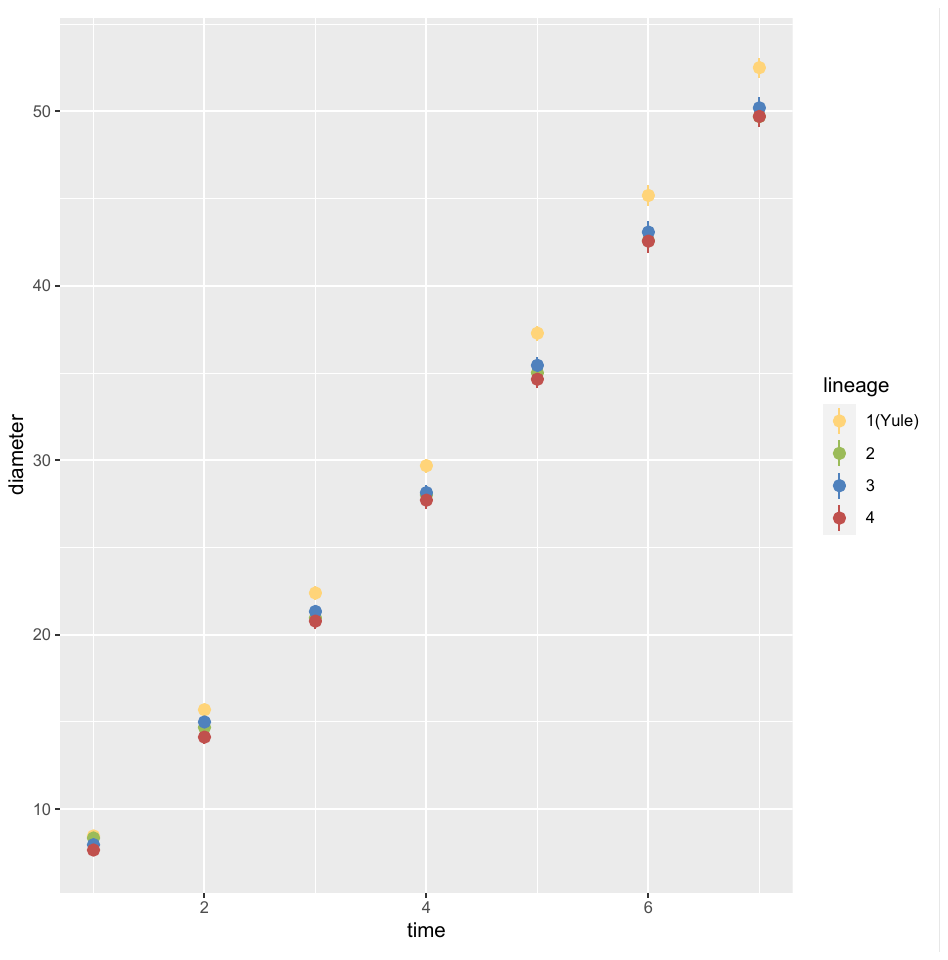

```
#T=30°C: linear mixed model
m.30 <- lmer(diameter ~ time + replicate+ lineage +(1 | isolate), data=T30, REML=T)

#T=30°C: plot residuals
plotresid(m.30)
```

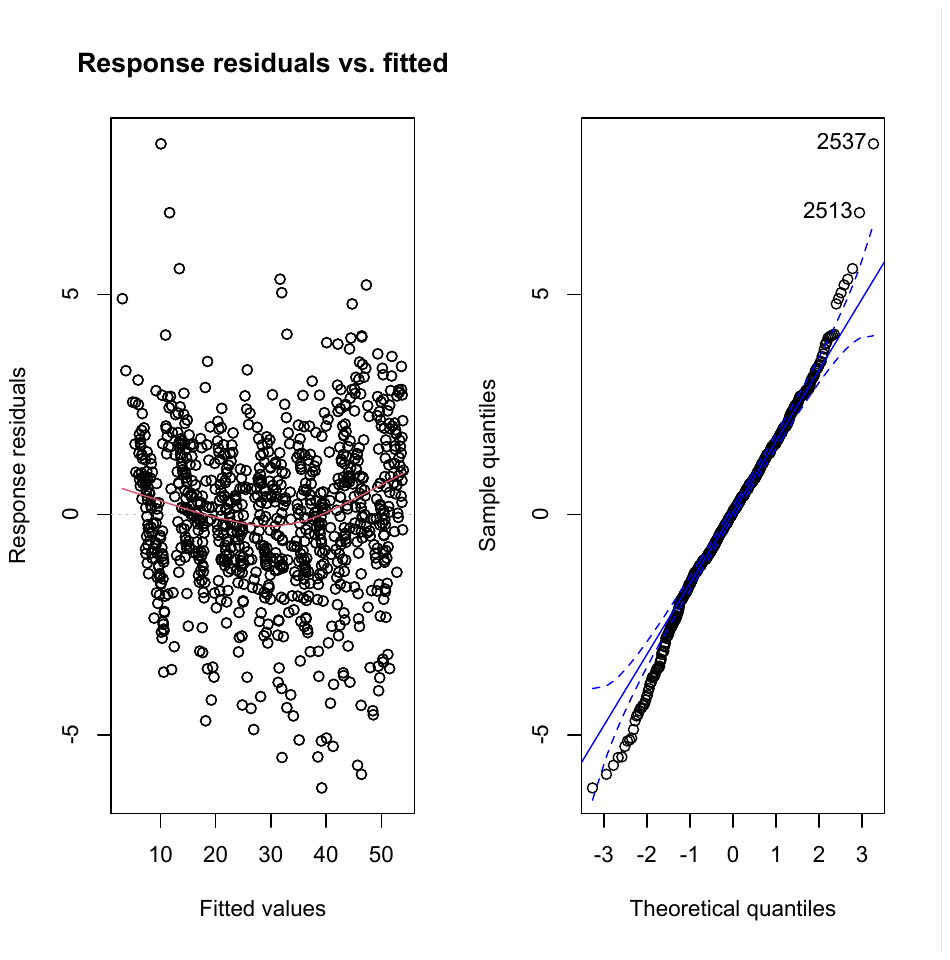

```
#T=30°C: ANOVA
anova( m.30)
## Type III Analysis of Variance Table with Satterthwaite's method
##                  Sum Sq Mean Sq NumDF  DenDF    F value  Pr(>F)
## time 178100  178100     1 870.34 52681.4995 < 2e-16 ***
## replicate         2063     688     3 881.49   203.4034 < 2e-16 ***
## lineage               39      13     3  37.07     3.8817 0.01649 *
## ---
## Signif. codes:  0 '***' 0.001 '**' 0.01 '*' 0.05 '.' 0.1 ' ' 1
summary(m.30)
## Linear mixed model fit by REML. t-tests use Satterthwaite's method [
## lmerModLmerTest]
## Formula: diameter ~ time + replicate + lineage + (1 |
##     isolate)
##    Data: T30
##
## REML criterion at convergence: 3818.9
##
## Scaled residuals:
##     Min      1Q  Median      3Q     Max
## -3.3748 -0.5624  0.0291  0.6219  4.5766
##
## Random effects:
##  Groups   Name        Variance Std.Dev.
##  isolate   (Intercept) 1.984    1.408
##  Residual             3.381    1.839
## Number of obs: 915, groups:  isolate, 41
##
## Fixed effects:
##                   Estimate Std. Error        df t value Pr(>|t|)
## (Intercept)        2.48054    0.47516  48.99784   5.220 3.62e-06 ***
## time   7.19095    0.03133 870.34024 229.525  < 2e-16 ***
## replicate2       -0.14130    0.18942 872.14201  -0.746  0.45592
## replicate3       -0.08508    0.17928 885.22910  -0.475  0.63523
## replicate4       -3.54333    0.18343 886.50289 -19.317  < 2e-16 ***
## lineage2           -1.76654    0.64236  37.14510  -2.750  0.00915 **
## lineage3           -1.55366    0.63917  36.50483  -2.431  0.02010 *
## lineage4           -1.90591    0.64205  37.15585  -2.968  0.00521 **
## ---
## Signif. codes:  0 '***' 0.001 '**' 0.01 '*' 0.05 '.' 0.1 ' ' 1
##
## Correlation of Fixed Effects:
##             (Intr) tmps__ rpttn2 rpttn3 rpttn4 ligne2 ligne3
## tmps_d_cltr -0.264
## replicate2 -0.195  0.003
## replicate3 -0.226 -0.002  0.511
## replicate4 -0.205 -0.070  0.499  0.604
## lineage2     -0.638  0.000  0.005  0.001 -0.007
## lineage3     -0.642  0.000  0.000  0.002  0.003  0.475
## lineage4     -0.635  0.001  0.000 -0.020 -0.018  0.473  0.475
#T=30°C: model fit
ggplot(T30, aes( time,diameter, color=lineage)) +
  stat_summary(fun.data=mean_se, geom="pointrange")+
  stat_summary(aes(y=fitted(m.30)),fun=mean,geom="line")
```

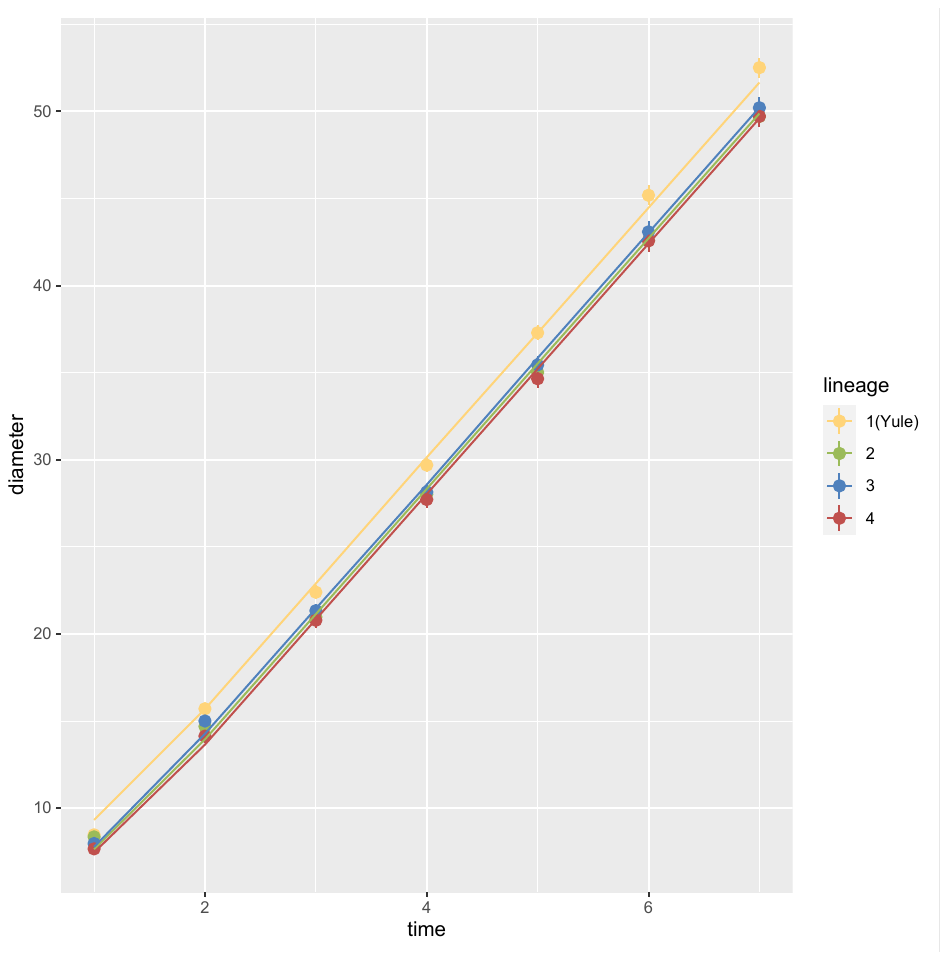

```
#T=30°C: least-squares means
LSM<-lsmeans(m.30,pairwise~lineage)
LSM
## $lsmeans
##  lineage lsmean    SE   df lower.CL upper.CL
##  1(Yule)   31.1 0.442 36.6     30.2     32.0
##  2        29.3 0.467 37.6     28.4     30.3
##  3        29.5 0.462 36.3     28.6     30.5
##  4        29.2 0.467 37.7     28.2     30.1
##
## Results are averaged over the levels of: replicate
## Degrees-of-freedom method: kenward-roger
## Confidence level used: 0.95
##
## $contrasts
##  contrast   estimate    SE   df t.ratio p.value
##  1(Yule) - 2    1.767 0.642 37.0  2.750  0.0435
##  1(Yule) - 3    1.554 0.639 36.4  2.431  0.0891
##  1(Yule) - 4    1.906 0.642 37.0  2.968  0.0257
##  2 - 3        -0.213 0.657 36.9 -0.324  0.9880
##  2 - 4         0.139 0.660 37.5  0.211  0.9966
##  3 - 4         0.352 0.657 36.9  0.536  0.9496
##
## Results are averaged over the levels of: replicate
## Degrees-of-freedom method: kenward-roger
## P value adjustment: tukey method for comparing a family of 4 estimates
cld(lsmeans(m.30,~lineage))
##  lineage lsmean    SE   df lower.CL upper.CL .group
##  4        29.2 0.467 37.7     28.2     30.1  1
##  2        29.3 0.467 37.6     28.4     30.3  1
##  3        29.5 0.462 36.3     28.6     30.5  12
##  1(Yule)   31.1 0.442 36.6     30.2     32.0   2
##
## Results are averaged over the levels of: replicate
## Degrees-of-freedom method: kenward-roger
## Confidence level used: 0.95
## P value adjustment: tukey method for comparing a family of 4 estimates
## significance level used: alpha = 0.05
```
