## Supplementary file 7 for "Ecological Differentiation Among Globally Distributed Lineages of the Rice Blast Fungus *Pyricularia oryzae*"

Supplementary file 7: Analysis of sporulation rates

We measured sporulation rate of 41 representative isolates cultured at different temperatures to test the hypothesis of adaptation to temperature. Main results are shown in Figure SF7-1.

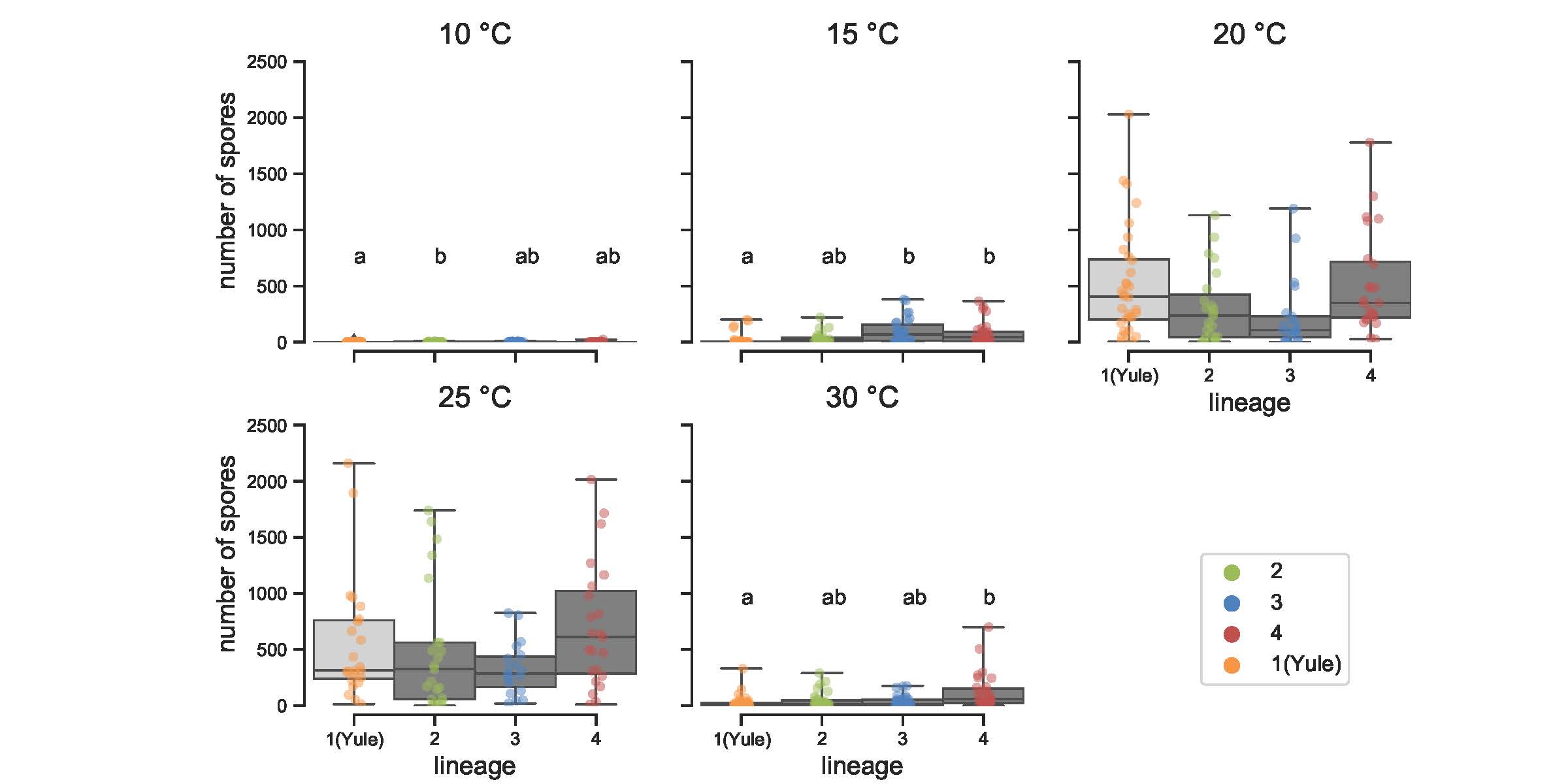

Figure SF7-1. Sporulation rates for lineages 2-4 and cluster *Yule* within lineage 1 of *P. oryzae* at five incubation temperatures, with three or four biological replicates. Each dot represents the median across three to four replicates. Letters indicate significant differences (Dunn’s non-parametric multiple comparison test, carried out only for temperatures for which Kruskal-Wallis tests were statistically significant).

In what follows, we detail the statistical processing of data:

#R packages (R version 4.0.3)
library(ggplot2)

library(MASS)

library(ggpubr)

library(RVAideMemoire)

### library(rstatix) #import data

```
data_spore <- read.csv("Supplementary_file_5.txt", sep="\t", na.strings="na")
data_spore$replicate <- as.factor(data_spore$replicate)
data_spore$lineage <- as.factor(data_spore$lineage)
data_spore$isolate <- as.factor(data_spore$isolate)


summary(data_spore)
    isolate     temperature    replicate     spores          lineage
 CH1120 : 17   Min.   :10.00   1:132     Min.   :   0.0   1(Yule):155
 US0032 : 17   1st Qu.:15.00   2:168     1st Qu.:   2.0   2      :124
 CH0999 : 16   Median :20.00   3:165     Median :  31.0   3      :131
 CH1065 : 16   Mean   :20.36   4: 74     Mean   : 189.2   4      :129
 CL0026 : 16   3rd Qu.:27.50             3rd Qu.: 222.5
 IN0072 : 16   Max.   :30.00             Max.   :2160.0
 (Other):441

#We take the median of the number of spores across replicates
data_spore_med_rep <- aggregate(data_spore$spores,data_spore[,c("temperature","lineage","isolate")], FUN=median)
colnames(data_spore_med_rep)[4] <- 'nb_spore'

#Round medians
data_spore_med_rep$nb_spore <- round(data_spore_med_rep$nb_spore)
```

#Make one dataset per temperature

data_sp_10_med <- **subset**(data_spore_med_rep, temperature**==**10)

data_sp_15_med <- **subset**(data_spore_med_rep, temperature**==**15)

data_sp_20_med <- **subset**(data_spore_med_rep, temperature**==**20)

data_sp_25_med <- **subset**(data_spore_med_rep, temperature**==**25)

data_sp_30_med <- **subset**(data_spore_med_rep, temperature**==**30)

#T=10°C: plot

ggplot(data_sp_10_med, aes( lineage,nb_spore, color=lineage))+ geom_boxplot()+ facet_wrap(~ temperature) + scale_color_manual(values=c("#FFD479", "#9BBB59", "#4F81BD","#C0504D"))+ labs(y="Number of spores", x="Lineages")

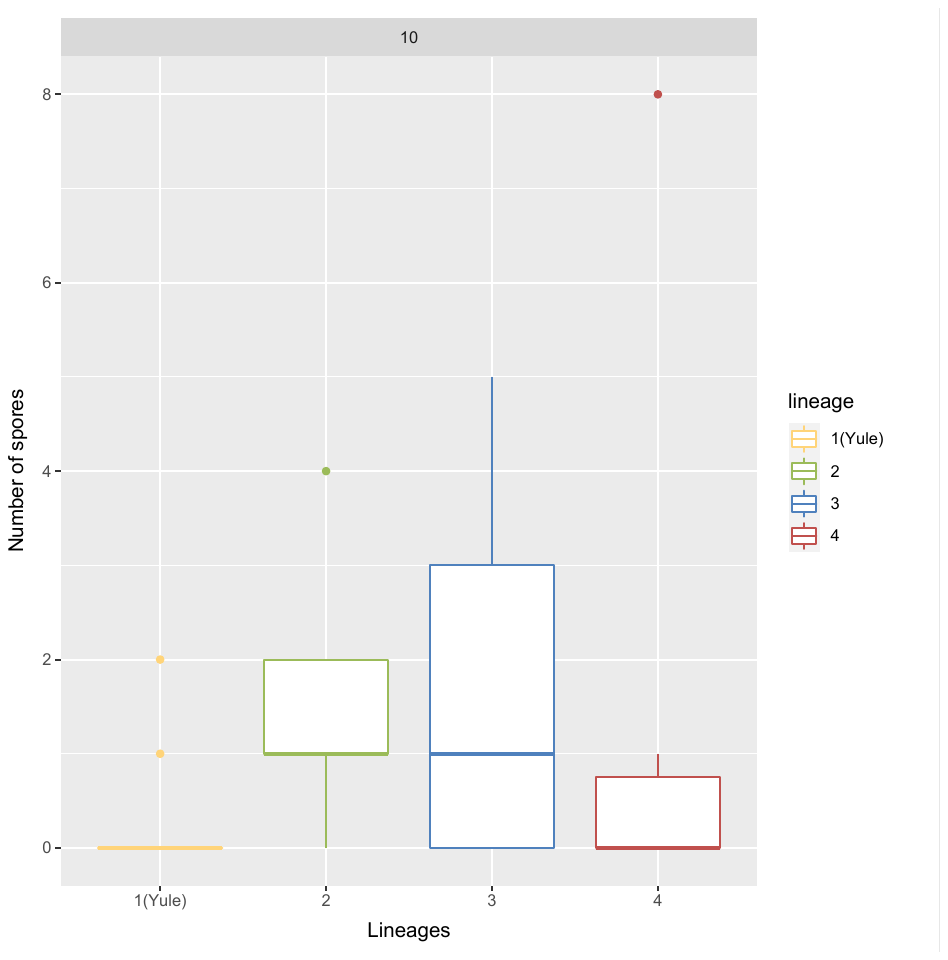

#T=15°C: plot

ggplot(data_sp_15_med, aes( lineage,nb_spore, color=lineage))+ geom_boxplot()+ facet_wrap(~ temperature) + scale_color_manual(values=c("#FFD479", "#9BBB59", "#4F81BD","#C0504D"))+ labs(y="Number of spores", x="Lineages")

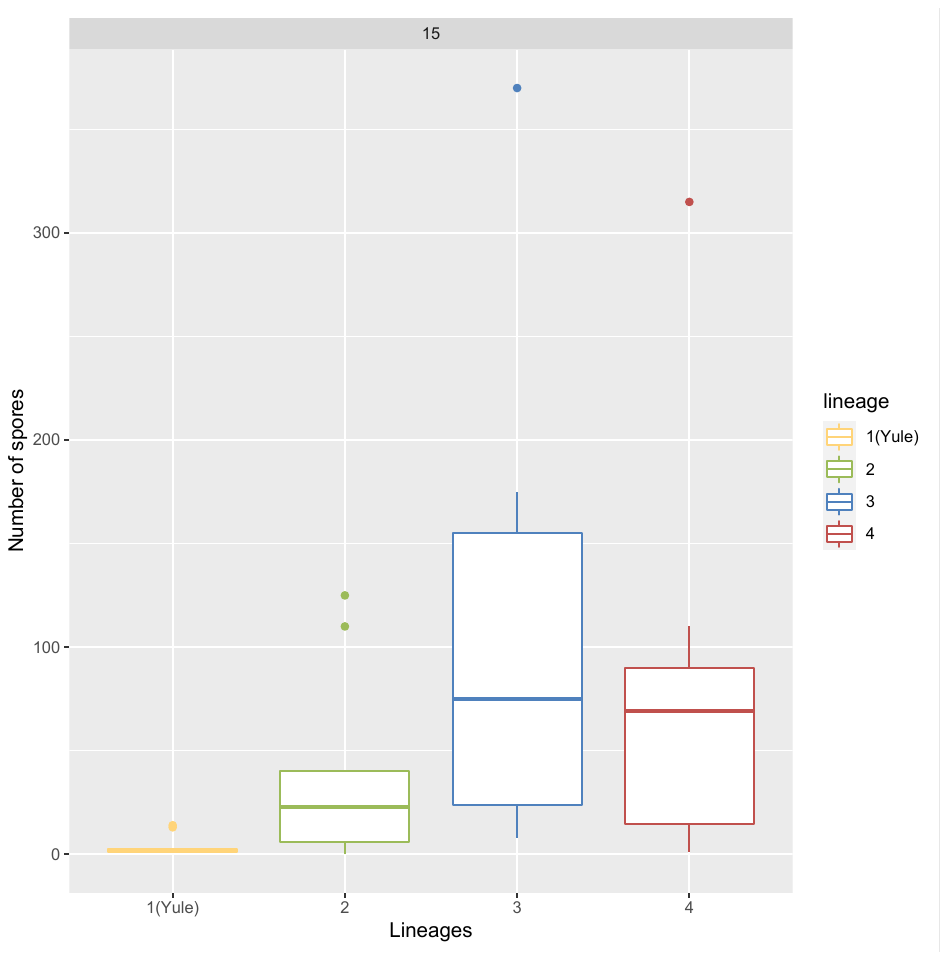

#T=20°C: plot

ggplot(data_sp_20_med, aes( lineage,nb_spore, color=lineage))+ geom_boxplot()+ facet_wrap(~ temperature) + scale_color_manual(values=c("#FFD479", "#9BBB59", "#4F81BD","#C0504D"))+ labs(y="Number of spores", x="Lineages")

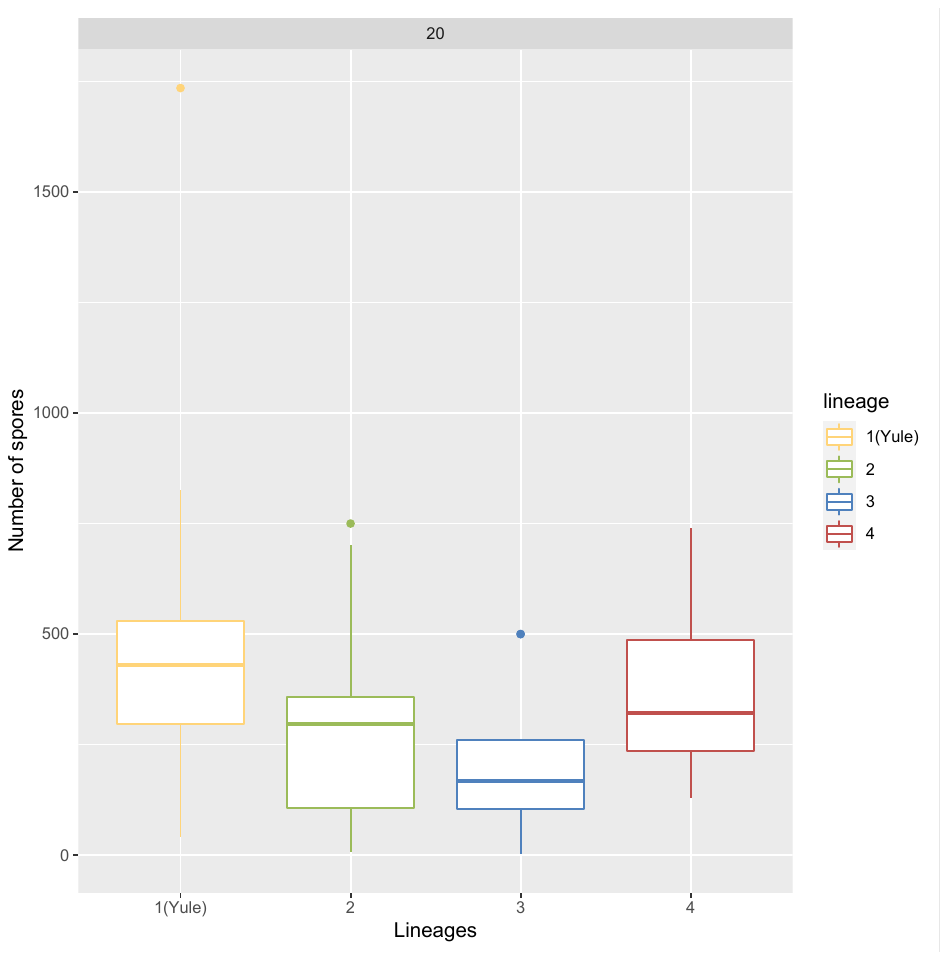

#T=25°C: plot

ggplot(data_sp_25_med, aes( lineage,nb_spore, color=lineage))+ geom_boxplot()+ facet_wrap(~ temperature) + scale_color_manual(values=c("#FFD479", "#9BBB59", "#4F81BD","#C0504D"))+ labs(y="Number of spores", x="Lineages")

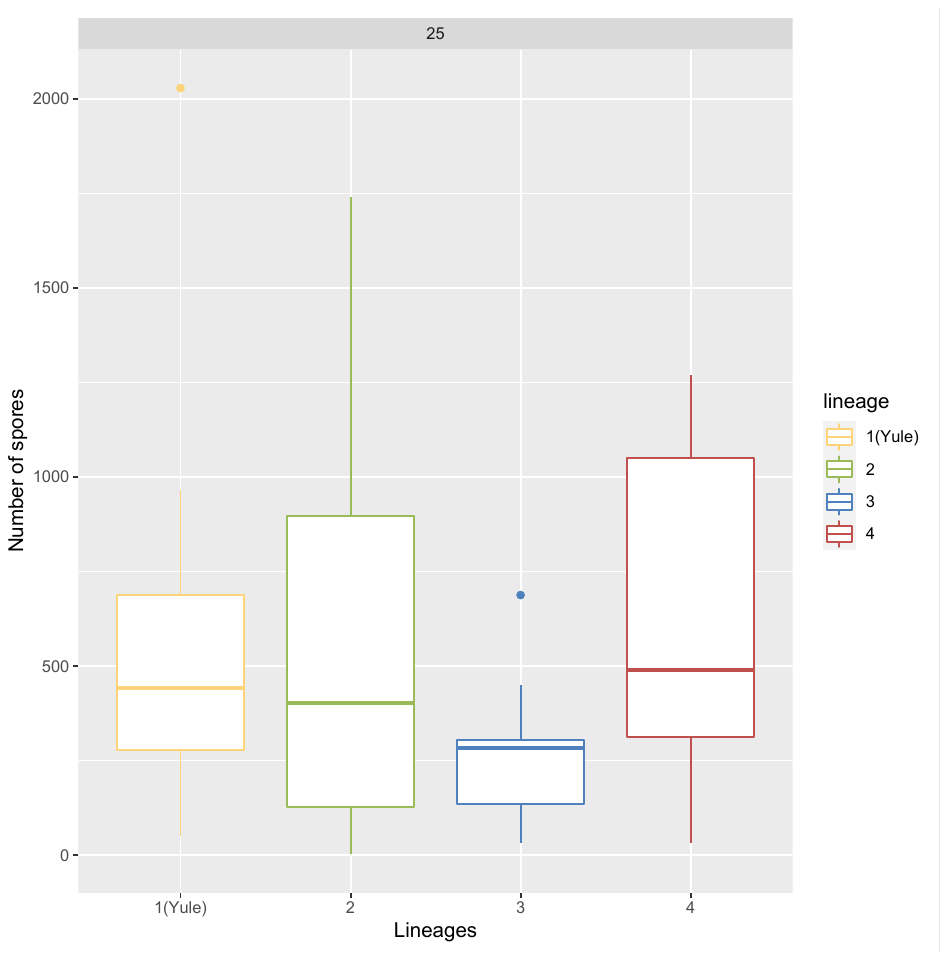

#T=30°C: plot

ggplot(data_sp_30_med, aes( lineage,nb_spore, color=lineage))+ geom_boxplot()+ facet_wrap(~ temperature) + scale_color_manual(values=c("#FFD479", "#9BBB59", "#4F81BD","#C0504D"))+ labs(y="Number of spores", x="Lineages")

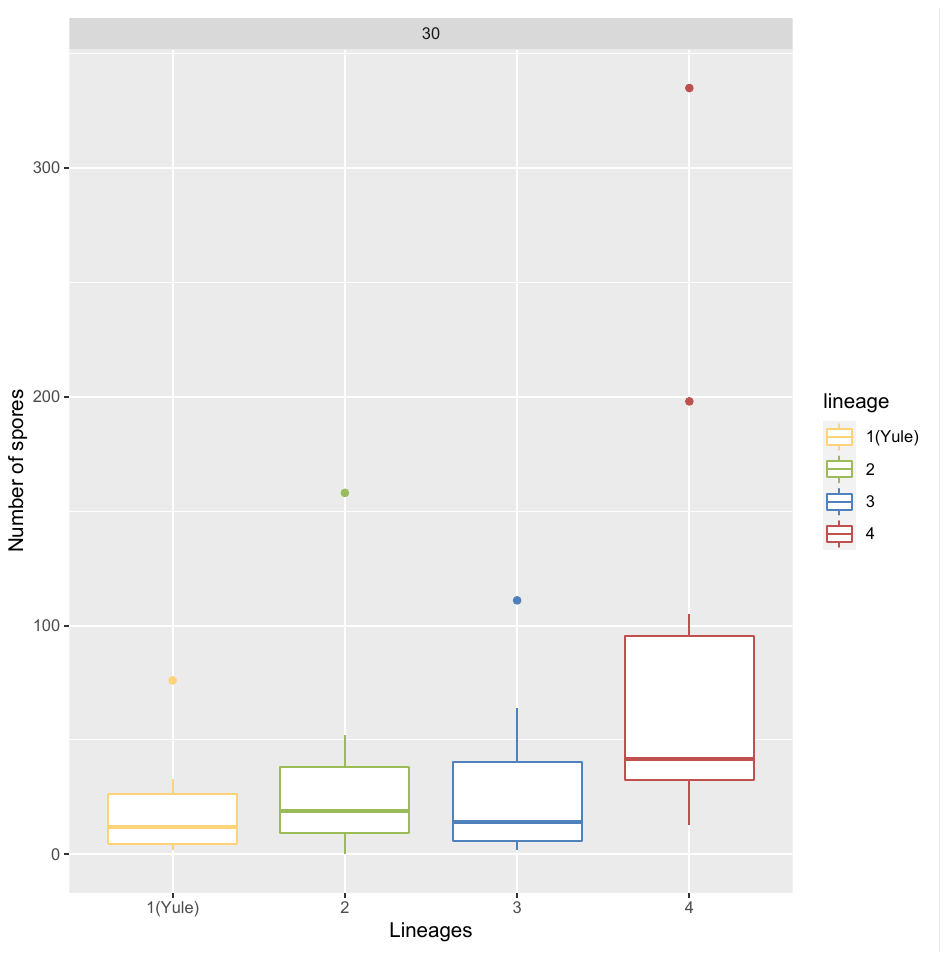

#Fit Negative Binomial Generalized Linear Models, and plot residuals

#We chose this type of model because it is suitable for over-dispersed count data.

mod.nb.pois10 <- **glm.nb**(nb_spore**~**lineage, data=data_sp_10_med)

**plotresid**(mod.nb.pois10)

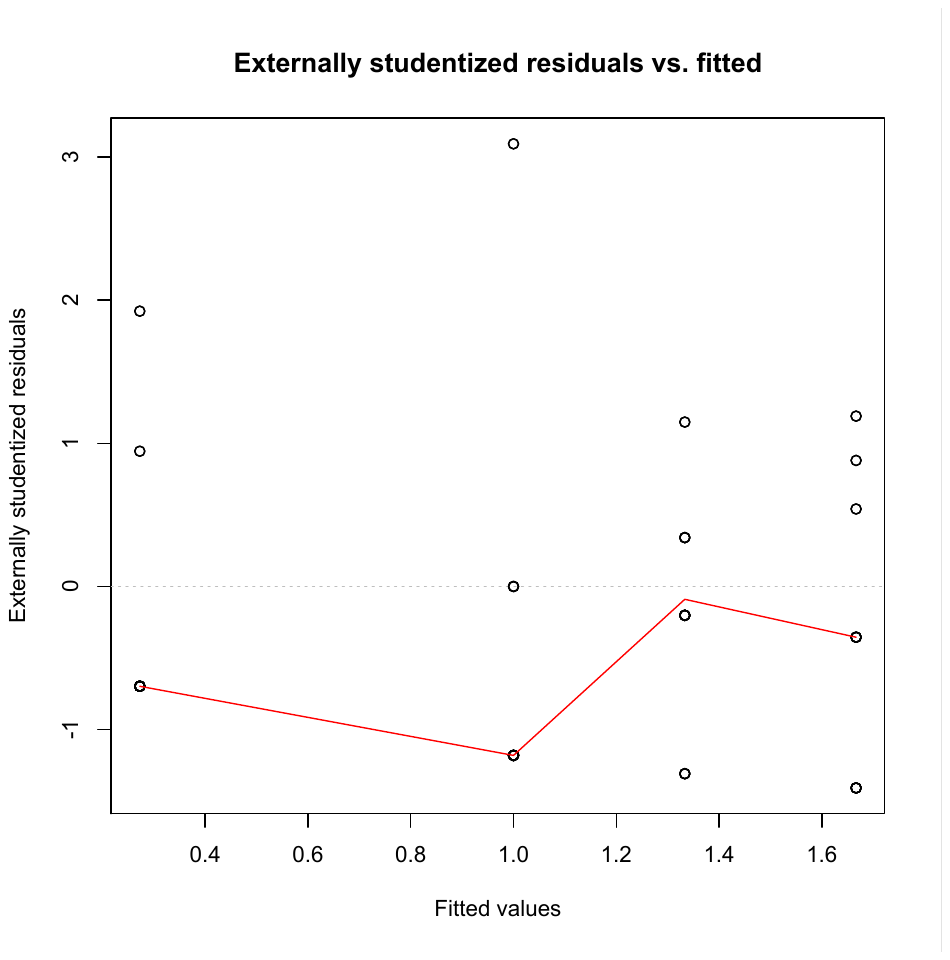

mod.nb.pois15 <- **glm.nb**(nb_spore**~**lineage, data=data_sp_15_med)

**plotresid**(mod.nb.pois15)

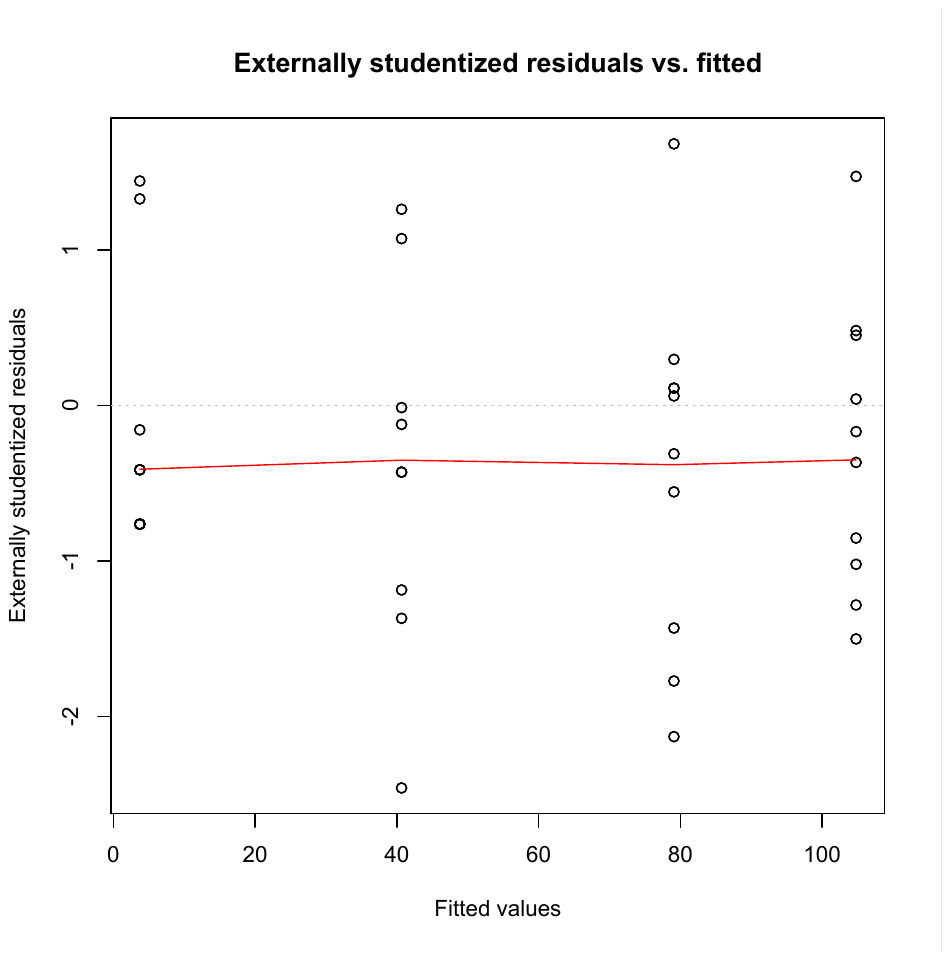

mod.nb.pois20 <- **glm.nb**(nb_spore**~**lineage, data=data_sp_20_med)

**plotresid**(mod.nb.pois20)

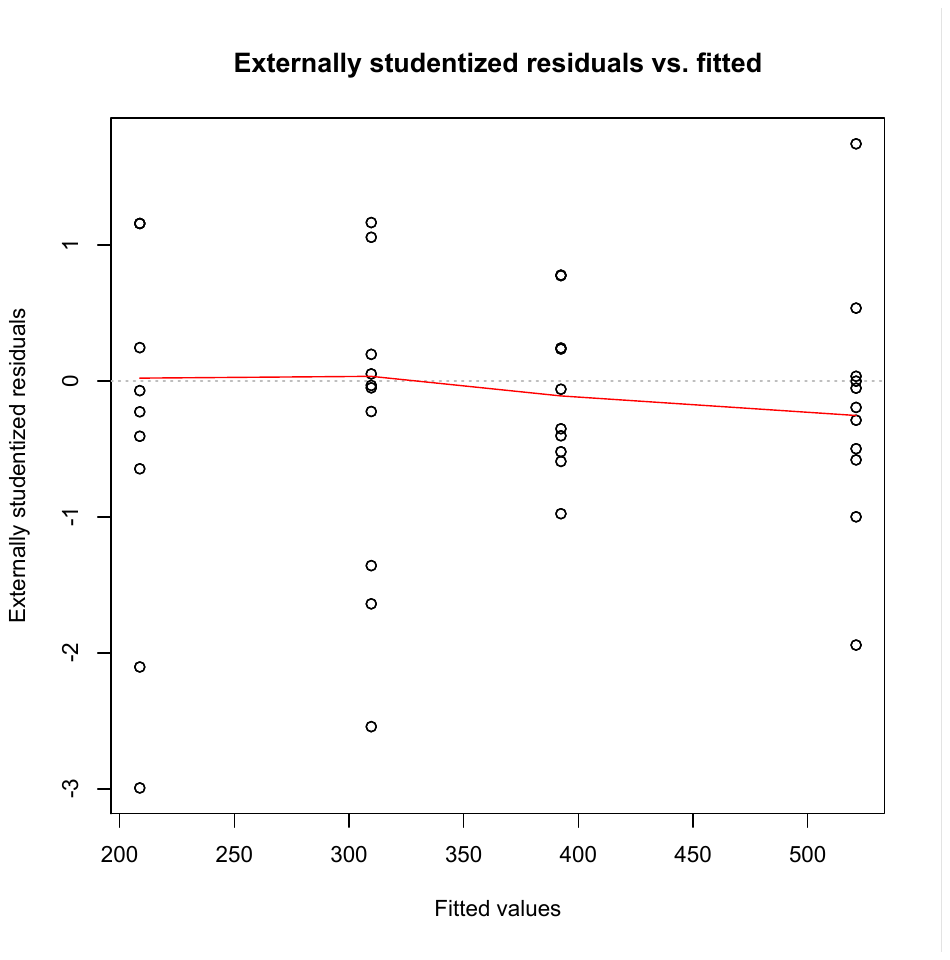

mod.nb.pois25 <- **glm.nb**(nb_spore**~**lineage, data=data_sp_25_med)

**plotresid**(mod.nb.pois25)

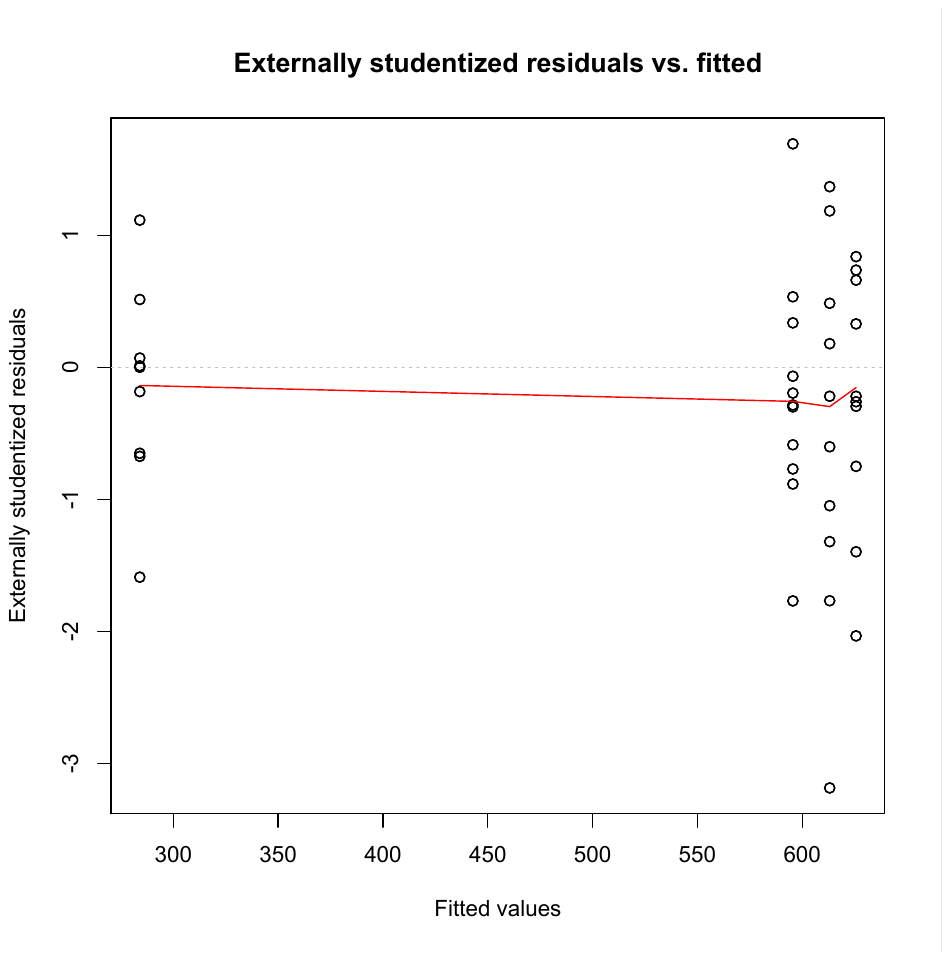

mod.nb.pois30 <- **glm.nb**(nb_spore**~**lineage, data=data_sp_30_med)

**plotresid**(mod.nb.pois30)

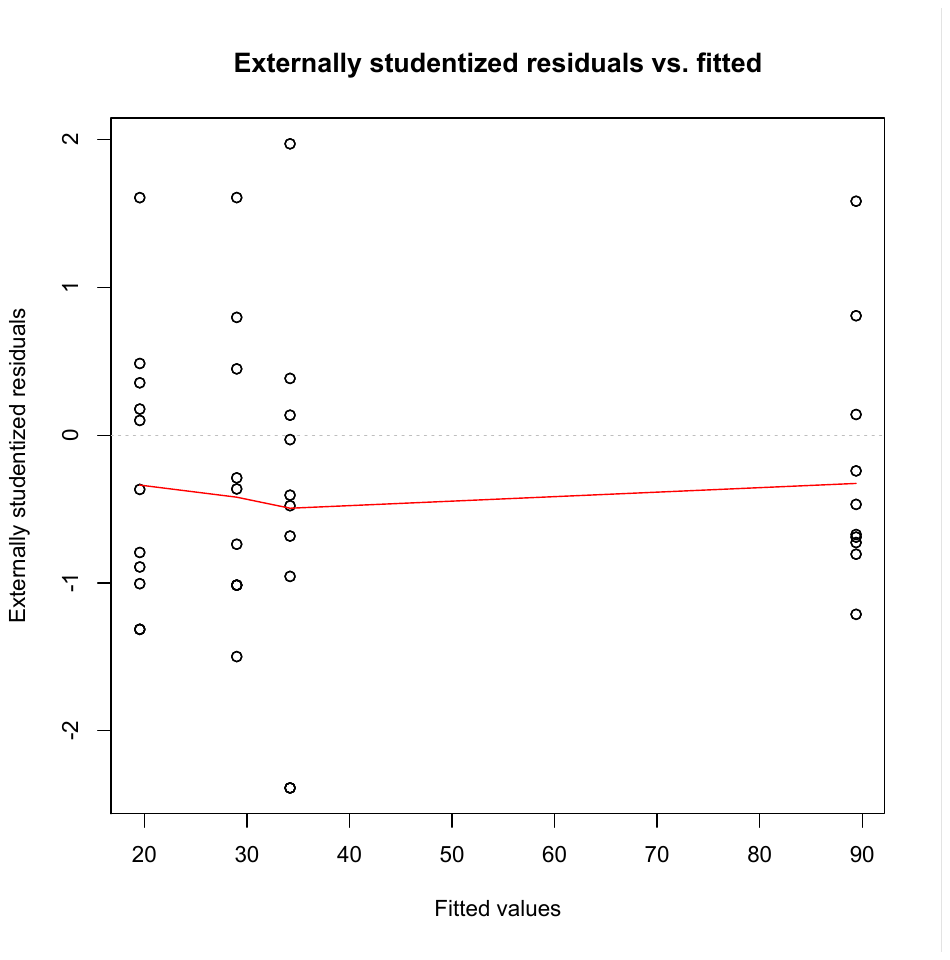

#Quantile-quantile plots to check if data are normally distributed

ggqqplot(mod.nb.pois10$residuals)

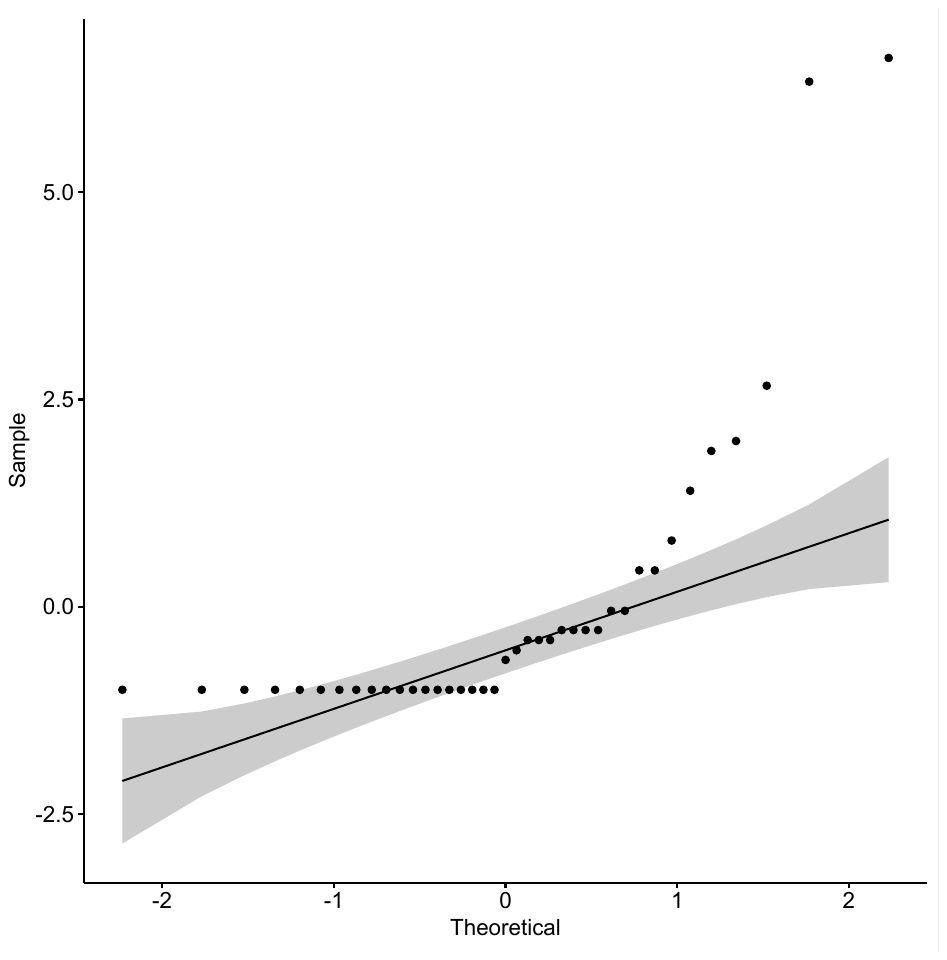

ggqqplot(mod.nb.pois15$residuals)

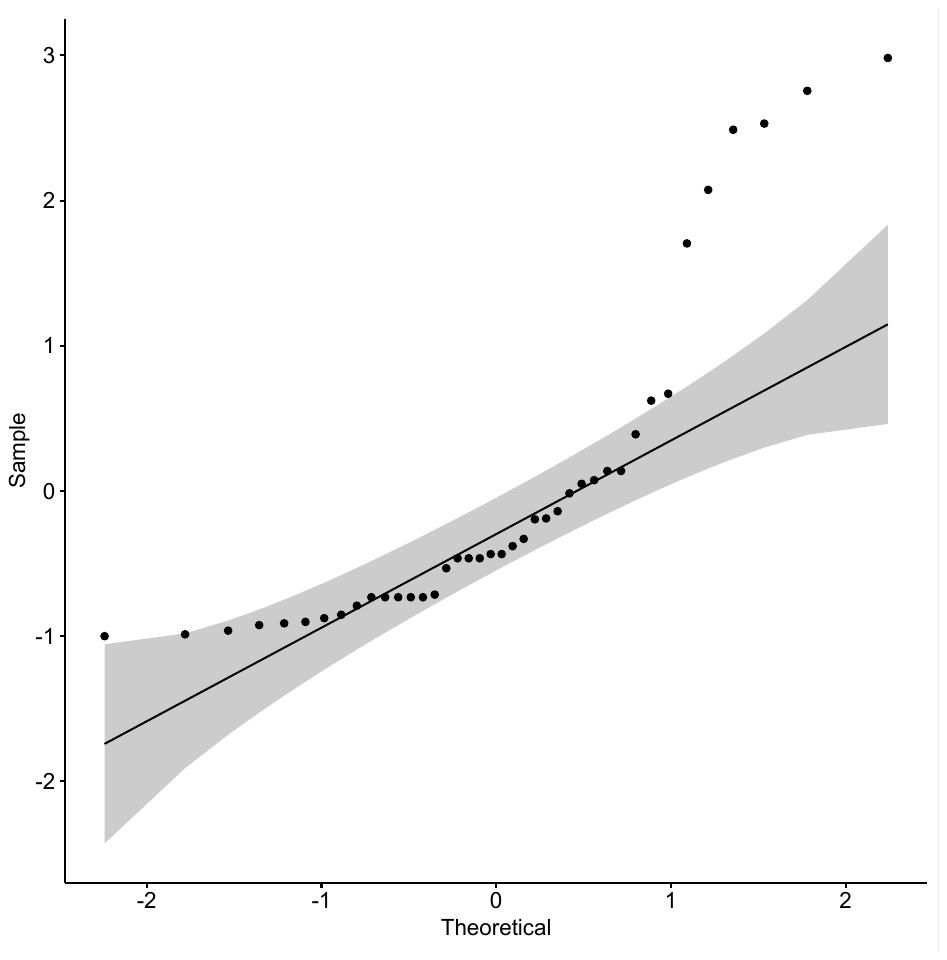

ggqqplot(mod.nb.pois20$residuals)

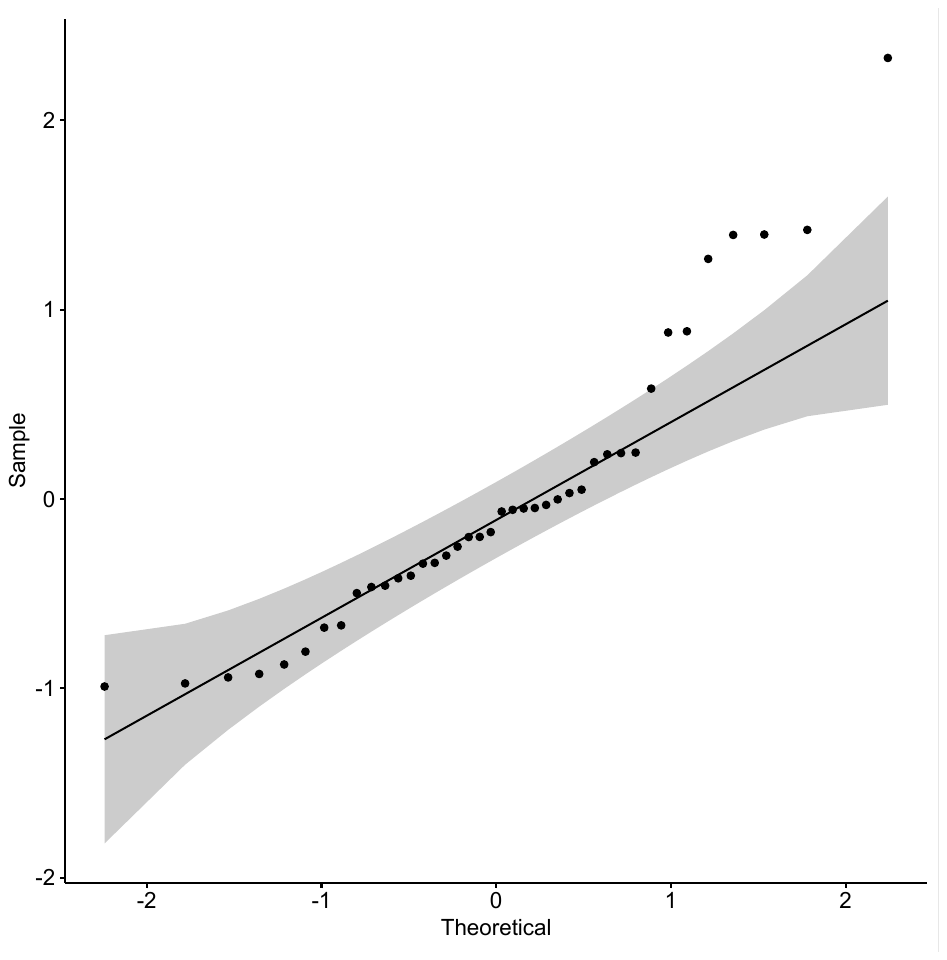

ggqqplot(mod.nb.pois25$residuals)

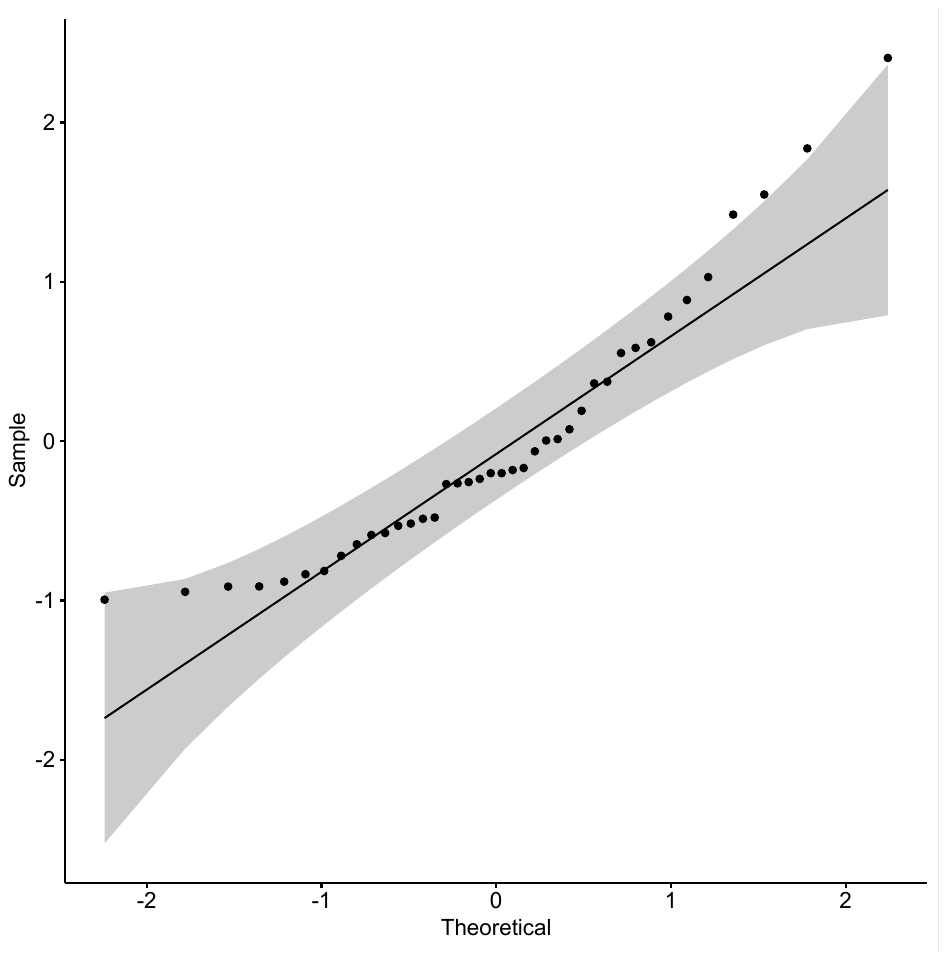

ggqqplot(mod.nb.pois30$residuals)

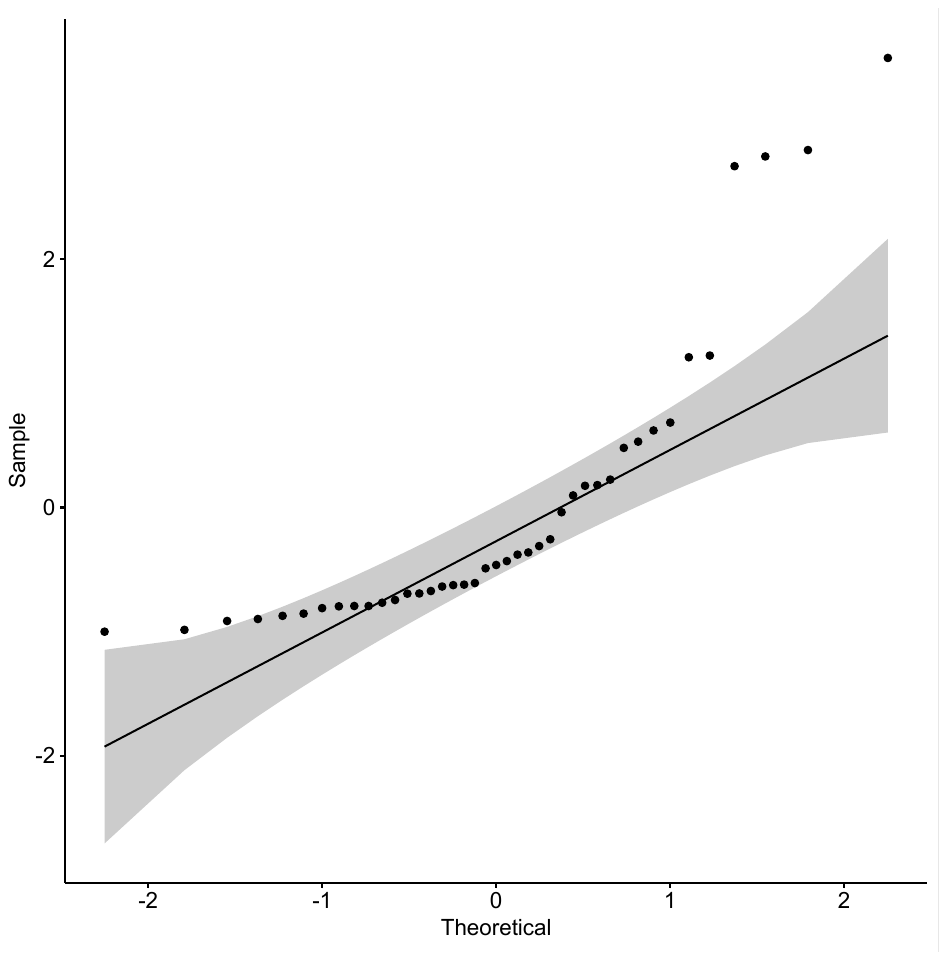

#Kruskal-Wallis test (as an alternative to ANOVA, given that data are not normally distributed)

res.kruskal <- data_sp_10_med %>% kruskal_test(nb_spore ~ lineage)

res.kruskal

\# A tibble: 1 x 6

.y. n statistic	df	p method

* <chr>	<int>	<dbl> <int> <dbl> <chr>

1 nb_spore	39	9.61	3 0.0222 Kruskal-Wallis

res.kruskal <- data_sp_15_med %>% kruskal_test(nb_spore ~ lineage)

res.kruskal

\# A tibble: 1 x 6

.y. n statistic	df	p method

* <chr>	<int>	<dbl> <int>	<dbl> <chr>

1 nb_spore	40	16.8	3 0.000761 Kruskal-Wallis

res.kruskal <- data_sp_20_med %>% kruskal_test(nb_spore ~ lineage)

res.kruskal

\# A tibble: 1 x 6

.y. n statistic	df	p method

* <chr>	<int>	<dbl> <int> <dbl> <chr>

1 nb_spore	40	6.04	3 0.11 Kruskal-Wallis

res.kruskal <- data_sp_25_med %>% kruskal_test(nb_spore ~ lineage)

res.kruskal

\# A tibble: 1 x 6

.y. n statistic	df	p method

* <chr>	<int>	<dbl> <int> <dbl> <chr>

1 nb_spore	40	3.42	3 0.331 Kruskal-Wallis

res.kruskal <- data_sp_30_med %>% kruskal_test(nb_spore ~ lineage)

res.kruskal

\# A tibble: 1 x 6

.y. n statistic	df	p method

* <chr>	<int>	<dbl> <int> <dbl> <chr>

1 nb_spore	41	8.96	3 0.0298 Kruskal-Wallis

#Dunn’s non-parametric multiple comparison test for temperatures for which Kruskal-Wallis tests were significant

PT = dunnTest(nb_spore ~ lineage,  data=data_sp_10_med)

Comparison	Z	P.unadj	P.adj

1 1(Yule) - 2 -2.7164706 0.006598206 0.03958924

2 1(Yule) - 3 -2.2508504 0.024395017 0.12197508

3	2 - 3 0.4439514 0.657077694 0.65707769

4 1(Yule) - 4 -0.7886814 0.430298220 0.86059644

5	2 - 4 1.9073416 0.056476364 0.22590546

6	3 - 4 1.4518571 0.146541356 0.43962407

PT=PT$res

cldList(comparison = PT$Comparison, p.value	= PT$P.adj, threshold = 0.05)

Group Letter MonoLetter

1 1(Yule)	a	a

2	2	b	b

3	3	ab	ab

4	4	ab	ab

PT = dunnTest(nb_spore ~ lineage,  data=data_sp_15_med)

Comparison	Z	P.unadj	P.adj

1 1(Yule) - 2 -2.3573282 0.0184069683 0.073627873

2 1(Yule) - 3 -3.8137339 0.0001368828 0.000821297

3	2 - 3 -1.3206571 0.1866157373 0.559847212

4 1(Yule) - 4 -3.1367872 0.0017081003 0.008540501

5	2 - 4 -0.6769145 0.4984602096 0.996920419

6	3 - 4 0.6613826 0.5083669692 0.508366969

PT=PT$res

cldList(comparison = PT$Comparison, p.value	= PT$P.adj, threshold = 0.05)

Group Letter MonoLetter

1 1(Yule)	a	a

2	2	ab	ab

3	3	b	b

4	4	b	b

PT = dunnTest(nb_spore ~ lineage,  data=data_sp_30_med)

Comparison	Z	P.unadj	P.adj

1 1(Yule) - 2 -0.7036481 0.481651938 1.00000000

2 1(Yule) - 3 -0.5412010 0.588369076 1.00000000

3	2 - 3 0.1587122 0.873895593 0.87389559

4 1(Yule) - 4 -2.8154612 0.004870728 0.02922437

5	2 - 4 -2.0632591 0.039088014 0.15635206

6	3 - 4 -2.2219714 0.026285240 0.13142620

PT=PT$res

cldList(comparison = PT$Comparison, p.value	= PT$P.adj, threshold = 0.05)

Group Letter MonoLetter

1 1(Yule)	a	a

2	2	ab	ab

3	3	ab	ab

4	4	b	b
