## Supplementary file 10 for "Ecological Differentiation Among Globally Distributed Lineages of the Rice Blast Fungus *Pyricularia oryzae*"

Supplementary file 10: statistical processing of pathogenicity data.

#R packages (R version 4.0.3)

library(ordinal)

library(RVAideMemoire)

library(lsmeans)

library(FSA)

library(lattice)

library(multcomp)

library(rcompanion)

#import data

patho_data <- read.csv("patho.txt",sep=";",na.strings=c("NA","na"))#patho.txt is Figure 4 – source data 1

patho_data$isolate<- as.factor(patho_data$isolate)

patho_data$clone<- as.factor(patho_data$clone)

patho_data$lineage<- as.factor(patho_data$lineage)

patho_data$ss.lineage<- as.factor(patho_data$ss.lineage)

patho_data$rice_type<- as.factor(patho_data$rice_type)

patho_data$variety<- as.factor(patho_data$variety)

patho_data$notes<-factor(patho_data$symptom_score,levels=c("1","2","3","4","5","6"),ordered=TRUE)

patho_data$typebin<-cut(patho_data$symptom_score,c(0,2,6),include.lowest=TRUE,labels = c("Incompatible","Compatible"))

summary(patho_data)

#Take the maximum symptom score across replicate

max_score<-aggregate(patho_data$symptom_score, patho_data[,c("isolate","lineage","ss.lineage","rice_type","variety")], FUN=max)

colnames(max_score)[6] <- 'symptom_score'

max_score$symptom_score<-round(max_score$symptom_score,0)

max_score$notes<-factor(max_score$symptom_score,levels=c("1","2","3","4","5","6"),ordered=TRUE)

\# fit a proportional odds model

max.clm1<-clm(data=max_score, notes~ss.lineage*rice_type,Hess=TRUE)

max.clm1

```
## formula: notes ~ ss.lineage * rice_type
## data:    max_score
##
##  link  threshold nobs logLik   AIC     niter max.grad cond.H
##  logit flexible  3080 -4083.68 8245.36 6(2)  6.26e-09 1.4e+04
##
## Coefficients:
##                        ss.lineage3                        ss.lineage4
##                          1.130758                          0.938238
##                  ss.lineageBaoshan                      ss.lineageInt
##                          1.053746                          0.780031
##                     ss.lineageLaos                     ss.lineageYule
##                          1.446904                          0.998254
##                   rice_typearomatic                       rice_typeind
##                         -0.408516                          0.250523
##                  rice_typejap_temp                   rice_typejap_trp
##                          1.727616                          0.006872
##        ss.lineage3:rice_typearomatic        ss.lineage4:rice_typearomatic
##                         -0.685654                         -0.042992
##  ss.lineageBaoshan:rice_typearomatic      ss.lineageInt:rice_typearomatic
##                         -0.062907                         -0.309470
##     ss.lineageLaos:rice_typearomatic     ss.lineageYule:rice_typearomatic
##                          0.141134                         -0.068622
##            ss.lineage3:rice_typeind            ss.lineage4:rice_typeind
##                          0.093780                         -0.397605
##      ss.lineageBaoshan:rice_typeind          ss.lineageInt:rice_typeind
##                         -0.053179                         -0.106520
##         ss.lineageLaos:rice_typeind         ss.lineageYule:rice_typeind
##                         -1.637128                          0.801246
##       ss.lineage3:rice_typejap_temp       ss.lineage4:rice_typejap_temp
##                         -0.378156                         -0.776136
## ss.lineageBaoshan:rice_typejap_temp     ss.lineageInt:rice_typejap_temp
##                         -0.864880                         -0.372044
##    ss.lineageLaos:rice_typejap_temp    ss.lineageYule:rice_typejap_temp
##                         -1.897844                         -0.526060
##        ss.lineage3:rice_typejap_trp        ss.lineage4:rice_typejap_trp
##                          0.262846                         -0.089005
##  ss.lineageBaoshan:rice_typejap_trp      ss.lineageInt:rice_typejap_trp
##                          0.145275                          0.722665
##     ss.lineageLaos:rice_typejap_trp     ss.lineageYule:rice_typejap_trp
##                         -0.541955                          0.691068
##
## Threshold coefficients:
##     1|2     2|3     3|4     4|5     5|6
## -1.1428  0.2255  0.4679  0.6320  0.9903
```

#ANOVA

anova(max.clm1)

```
## Type I Analysis of Deviance Table with Wald chi-square tests
##
##                    Df   Chisq Pr(>Chisq)
## ss.lineage           6  99.591  < 2.2e-16 ***
## rice_type            4 161.249  < 2.2e-16 ***
## ss.lineage:rice_type 24  97.114  9.288e-11 ***
## ---
## Signif. codes:  0 '***' 0.001 '**' 0.01 '*' 0.05 '.' 0.1 ' ' 1
```

#Fit a proportional odds model without interactions to carry out posthoc tests

max.clm2<-clm(data=max_score, notes~ss.lineage+rice_type,Hess=TRUE)

max.clm2

```
## formula: notes ~ ss.lineage + rice_type
## data:    max_score
##
##  link  threshold nobs logLik   AIC     niter max.grad cond.H
##  logit flexible  3080 -4133.45 8296.90 6(2)  7.18e-10 1.1e+03
##
## Coefficients:
##       ss.lineage3       ss.lineage4 ss.lineageBaoshan     ss.lineageInt
##          0.99189          0.71707          0.91894          0.76240
##    ss.lineageLaos    ss.lineageYule  rice_typearomatic      rice_typeind
##          0.75610          1.13995         -0.55516          0.01813
## rice_typejap_temp  rice_typejap_trp
##          1.01643          0.15485
##
## Threshold coefficients:
##      1|2      2|3      3|4      4|5      5|6
## -1.26229  0.07864  0.31418  0.47358  0.82169
```

#ANOVA

anova(max.clm2)

#Contrast lineages using least-squares means

```
## Type I Analysis of Deviance Table with Wald chi-square tests
##
##           Df  Chisq Pr(>Chisq)
## ss.lineage  6 103.67  < 2.2e-16 ***
## rice_type   4 163.40  < 2.2e-16 ***
## ---
## Signif. codes:  0 '***' 0.001 '**' 0.01 '*' 0.05 '.' 0.1 ' ' 1
```

LSM <- lsmeans(max.clm2,~ss.lineage,adjust="sidak")

contrast(LSM,"pairwise")

```
##  contrast       estimate    SE  df z.ratio p.value
##  2 - 3           -0.9919 0.126 Inf -7.867  <.0001
##  2 - 4           -0.7171 0.125 Inf -5.733  <.0001
##  2 - Baoshan     -0.9189 0.129 Inf -7.149  <.0001
##  2 - Int         -0.7624 0.125 Inf -6.103  <.0001
##  2 - Laos        -0.7561 0.122 Inf -6.188  <.0001
##  2 - Yule        -1.1399 0.128 Inf -8.911  <.0001
##  3 - 4            0.2748 0.130 Inf  2.106  0.3488
##  3 - Baoshan      0.0730 0.134 Inf  0.546  0.9981
##  3 - Int          0.2295 0.130 Inf  1.760  0.5751
##  3 - Laos         0.2358 0.128 Inf  1.845  0.5170
##  3 - Yule        -0.1481 0.133 Inf -1.114  0.9239
##  4 - Baoshan     -0.2019 0.133 Inf -1.519  0.7335
##  4 - Int         -0.0453 0.130 Inf -0.350  0.9999
##  4 - Laos        -0.0390 0.127 Inf -0.307  0.9999
##  4 - Yule        -0.4229 0.132 Inf -3.199  0.0233
##  Baoshan - Int    0.1565 0.133 Inf  1.179  0.9023
##  Baoshan - Laos   0.1628 0.130 Inf  1.250  0.8743
##  Baoshan - Yule  -0.2210 0.135 Inf -1.634  0.6601
##  Int - Laos       0.0063 0.127 Inf  0.050  1.0000
##  Int - Yule      -0.3775 0.132 Inf -2.859  0.0644
##  Laos - Yule     -0.3838 0.129 Inf -2.964  0.0478
##
## Results are averaged over the levels of: rice_type
## P value adjustment: tukey method for comparing a family of 7 estimates
```

cld(LSM)

```
##  ss.lineage lsmean     SE  df asymp.LCL asymp.UCL .group
##  2         0.0417 0.0851 Inf    -0.187      0.27  1
##  4         0.7588 0.0926 Inf     0.510      1.01   2
##  Laos      0.7978 0.0889 Inf     0.559      1.04   2
##  Int       0.8041 0.0925 Inf     0.556      1.05   23
##  Baoshan   0.9606 0.0972 Inf     0.700      1.22   23
##  3         1.0336 0.0940 Inf     0.781      1.29   23
##  Yule      1.1816 0.0964 Inf     0.923      1.44    3
##
## Results are averaged over the levels of: rice_type
## Confidence level used: 0.95
## Conf-level adjustment: sidak method for 7 estimates
## P value adjustment: tukey method for comparing a family of 7 estimates
## significance level used: alpha = 0.05
```

#Contrast rice-types using least-squares means

LSM <- lsmeans(max.clm2,~rice_type,adjust="sidak")

contrast(LSM,"pairwise")

```
##  contrast           estimate    SE  df z.ratio p.value
##  aus - aromatic        0.5552 0.102 Inf   5.433 <.0001
##  aus - ind           -0.0181 0.100 Inf  -0.181 0.9998
##  aus - jap_temp      -1.0164 0.110 Inf  -9.245 <.0001
##  aus - jap_trp       -0.1549 0.103 Inf  -1.506 0.5583
##  aromatic - ind       -0.5733 0.115 Inf  -4.966 <.0001
##  aromatic - jap_temp  -1.5716 0.125 Inf -12.612 <.0001
##  aromatic - jap_trp   -0.7100 0.118 Inf  -6.024 <.0001
##  ind - jap_temp      -0.9983 0.122 Inf  -8.156 <.0001
##  ind - jap_trp       -0.1367 0.116 Inf  -1.178 0.7638
##  jap_temp - jap_trp   0.8616 0.125 Inf   6.919 <.0001
##
## Results are averaged over the levels of: ss.lineage
## P value adjustment: tukey method for comparing a family of 5 estimates
```

cld(LSM)

```
##  rice_type lsmean     SE  df asymp.LCL asymp.UCL .group
##  aromatic   0.115 0.0826 Inf   -0.0974     0.327  1
##  aus       0.670 0.0603 Inf    0.5152     0.825   2
##  ind       0.688 0.0808 Inf    0.4806     0.896   2
##  jap_trp   0.825 0.0841 Inf    0.6088     1.041   2
##  jap_temp  1.686 0.0934 Inf    1.4466     1.926    3
##
## Results are averaged over the levels of: ss.lineage
## Confidence level used: 0.95
## Conf-level adjustment: sidak method for 5 estimates
## P value adjustment: tukey method for comparing a family of 5 estimates
## significance level used: alpha = 0.05
```

\# generate datasets specific to each rice-type

max_score_aus <- subset(max_score, rice_type == "aus")

max_score_aus <- droplevels(max_score_aus )

max_score_bas <- subset(max_score, rice_type == "aromatic")

max_score_bas <- droplevels(max_score_bas )

max_score_ind <- subset(max_score, rice_type == "ind")

max_score_ind <- droplevels(max_score_ind )

max_score_temp <- subset(max_score, rice_type == "jap_temp")

max_score_temp <- droplevels(max_score_temp )

max_score_trp <- subset(max_score, rice_type == "jap_trp")

max_score_trp <- droplevels(max_score_trp )

#Contrast lineages inoculated onto Aus using least-squares means

max.aus<-clm(data=max_score_aus, notes~ss.lineage,Hess=T)

anova(max.aus)

```
## Type I Analysis of Deviance Table with Wald chi-square tests
##
##           Df  Chisq Pr(>Chisq)
## ss.lineage  6 50.585  3.588e-09 ***
## ---
## Signif. codes:  0 '***' 0.001 '**' 0.01 '*' 0.05 '.' 0.1 ' ' 1
```

LSM <- lsmeans(max.aus,~ss.lineage,adjust="sidak")

contrast(LSM,"pairwise")

```
##  contrast       estimate    SE  df z.ratio p.value
##  2 - 3           -1.1071 0.218 Inf -5.085  <.0001
##  2 - 4           -0.9181 0.218 Inf -4.205  0.0005
##  2 - Baoshan     -1.0303 0.222 Inf -4.638  0.0001
##  2 - Int         -0.7628 0.216 Inf -3.534  0.0075
##  2 - Laos        -1.4213 0.218 Inf -6.532  <.0001
##  2 - Yule        -0.9764 0.217 Inf -4.498  0.0001
##  3 - 4            0.1890 0.227 Inf  0.833  0.9815
##  3 - Baoshan      0.0768 0.230 Inf  0.334  0.9999
##  3 - Int          0.3443 0.225 Inf  1.533  0.7251
##  3 - Laos        -0.3142 0.225 Inf -1.394  0.8051
##  3 - Yule         0.1307 0.226 Inf  0.579  0.9974
##  4 - Baoshan     -0.1122 0.231 Inf -0.486  0.9990
##  4 - Int          0.1553 0.225 Inf  0.689  0.9933
##  4 - Laos        -0.5032 0.226 Inf -2.224  0.2827
##  4 - Yule        -0.0583 0.227 Inf -0.257  1.0000
##  Baoshan - Int    0.2675 0.229 Inf  1.168  0.9061
##  Baoshan - Laos  -0.3909 0.230 Inf -1.702  0.6146
##  Baoshan - Yule   0.0539 0.230 Inf  0.234  1.0000
##  Int - Laos      -0.6585 0.224 Inf -2.938  0.0516
##  Int - Yule      -0.2136 0.224 Inf -0.952  0.9639
##  Laos - Yule      0.4449 0.225 Inf  1.975  0.4306
##
## P value adjustment: tukey method for comparing a family of 7 estimates
```

cld(LSM)

```
##  ss.lineage lsmean    SE  df asymp.LCL asymp.UCL .group
##  2         -0.222 0.146 Inf    -0.615     0.170  1
##  Int        0.541 0.158 Inf     0.116     0.966   2
##  4          0.696 0.162 Inf     0.262     1.129   2
##  Yule       0.754 0.160 Inf     0.325     1.183   2
##  Baoshan    0.808 0.167 Inf     0.361     1.255   2
##  3          0.885 0.161 Inf     0.454     1.316   2
##  Laos       1.199 0.160 Inf     0.769     1.629   2
##
## Confidence level used: 0.95
## Conf-level adjustment: sidak method for 7 estimates
## P value adjustment: tukey method for comparing a family of 7 estimates
## significance level used: alpha = 0.05
```

#Contrast lineages inoculated onto Aromatic using least-squares means

max.bas<-clm(data=max_score_bas, notes~ss.lineage,Hess=T)

anova(max.bas)

```
## Type I Analysis of Deviance Table with Wald chi-square tests
##
##           Df  Chisq Pr(>Chisq)
## ss.lineage  6 29.585  4.714e-05 ***
## ---
## Signif. codes:  0 '***' 0.001 '**' 0.01 '*' 0.05 '.' 0.1 ' ' 1
```

LSM <- lsmeans(max.bas,~ss.lineage,adjust="sidak")

contrast(LSM,"pairwise")

```
##  contrast       estimate    SE  df z.ratio p.value
##  2 - 3           -0.4271 0.300 Inf -1.425  0.7884
##  2 - 4           -0.8591 0.306 Inf -2.807  0.0741
##  2 - Baoshan     -0.9568 0.308 Inf -3.107  0.0312
##  2 - Int         -0.4522 0.303 Inf -1.492  0.7500
##  2 - Laos        -1.5316 0.310 Inf -4.934  <.0001
##  2 - Yule        -0.8943 0.299 Inf -2.995  0.0437
##  3 - 4           -0.4320 0.309 Inf -1.400  0.8023
##  3 - Baoshan     -0.5297 0.310 Inf -1.707  0.6112
##  3 - Int         -0.0250 0.307 Inf -0.082  1.0000
##  3 - Laos        -1.1044 0.312 Inf -3.538  0.0074
##  3 - Yule        -0.4672 0.301 Inf -1.551  0.7139
##  4 - Baoshan     -0.0977 0.315 Inf -0.310  0.9999
##  4 - Int          0.4070 0.312 Inf  1.304  0.8503
##  4 - Laos        -0.6724 0.316 Inf -2.127  0.3367
##  4 - Yule        -0.0352 0.306 Inf -0.115  1.0000
##  Baoshan - Int    0.5047 0.314 Inf  1.609  0.6766
##  Baoshan - Laos  -0.5747 0.317 Inf -1.811  0.5406
##  Baoshan - Yule   0.0625 0.308 Inf  0.203  1.0000
##  Int - Laos      -1.0794 0.316 Inf -3.421  0.0112
##  Int - Yule      -0.4422 0.305 Inf -1.451  0.7738
##  Laos - Yule      0.6372 0.309 Inf  2.062  0.3755
##
## P value adjustment: tukey method for comparing a family of 7 estimates
```

cld(LSM)

```
##  ss.lineage lsmean    SE  df asymp.LCL asymp.UCL .group
##  2         -0.624 0.211 Inf    -1.190   -0.0582  1
##  3         -0.197 0.215 Inf    -0.774    0.3793  12
##  Int       -0.172 0.220 Inf    -0.762    0.4174  12
##  4          0.235 0.222 Inf    -0.360    0.8301  123
##  Yule       0.270 0.211 Inf    -0.297    0.8372   23
##  Baoshan    0.332 0.224 Inf    -0.267    0.9324   23
##  Laos       0.907 0.226 Inf     0.301    1.5136    3
##
## Confidence level used: 0.95
## Conf-level adjustment: sidak method for 7 estimates
## P value adjustment: tukey method for comparing a family of 7 estimates
## significance level used: alpha = 0.05
```

#Contrast lineages inoculated onto Indica using least-squares means

max.ind<-clm(data=max_score_ind, notes~ss.lineage,Hess=T)

anova(max.ind)

```
## Type I Analysis of Deviance Table with Wald chi-square tests
##
##           Df  Chisq Pr(>Chisq)
## ss.lineage  6 52.523  1.463e-09 ***
## ---
## Signif. codes:  0 '***' 0.001 '**' 0.01 '*' 0.05 '.' 0.1 ' ' 1
```

LSM <- lsmeans(max.ind,~ss.lineage,adjust="sidak")

contrast(LSM,"pairwise")

```
##  contrast       estimate    SE  df z.ratio p.value
##  2 - 3            -1.161 0.299 Inf -3.886  0.0020
##  2 - 4            -0.502 0.287 Inf -1.748  0.5838
##  2 - Baoshan      -0.950 0.306 Inf -3.106  0.0313
##  2 - Int          -0.635 0.289 Inf -2.196  0.2974
##  2 - Laos          0.164 0.272 Inf  0.604  0.9967
##  2 - Yule         -1.717 0.320 Inf -5.360  <.0001
##  3 - 4             0.658 0.306 Inf  2.149  0.3240
##  3 - Baoshan       0.211 0.323 Inf  0.654  0.9949
##  3 - Int           0.526 0.308 Inf  1.710  0.6096
##  3 - Laos          1.325 0.295 Inf  4.497  0.0001
##  3 - Yule         -0.556 0.335 Inf -1.658  0.6441
##  4 - Baoshan      -0.447 0.314 Inf -1.426  0.7879
##  4 - Int          -0.132 0.298 Inf -0.444  0.9994
##  4 - Laos          0.666 0.283 Inf  2.357  0.2170
##  4 - Yule         -1.214 0.327 Inf -3.715  0.0038
##  Baoshan - Int     0.315 0.315 Inf  1.000  0.9541
##  Baoshan - Laos    1.114 0.301 Inf  3.694  0.0041
##  Baoshan - Yule   -0.767 0.342 Inf -2.241  0.2734
##  Int - Laos        0.799 0.284 Inf  2.810  0.0735
##  Int - Yule       -1.082 0.328 Inf -3.298  0.0169
##  Laos - Yule      -1.881 0.317 Inf -5.938  <.0001
##
## P value adjustment: tukey method for comparing a family of 7 estimates
```

cld(LSM)

```
##  ss.lineage  lsmean    SE  df asymp.LCL asymp.UCL .group
##  Laos      -0.1385 0.189 Inf   -0.6445     0.368  1
##  2          0.0256 0.196 Inf   -0.5007     0.552  1
##  4          0.5280 0.210 Inf   -0.0357     1.092  12
##  Int        0.6602 0.212 Inf    0.0909     1.230  12
##  Baoshan    0.9752 0.234 Inf    0.3462     1.604   23
##  3          1.1864 0.225 Inf    0.5826     1.790   23
##  Yule       1.7424 0.253 Inf    1.0640     2.421    3
##
## Confidence level used: 0.95
## Conf-level adjustment: sidak method for 7 estimates
## P value adjustment: tukey method for comparing a family of 7 estimates
## significance level used: alpha = 0.05
```

#Contrast lineages inoculated onto Temperate japonica using least-squares means

max.temp<-clm(data=max_score_temp, notes~ss.lineage,Hess=T)

anova(max.temp)

```
## Type I Analysis of Deviance Table with Wald chi-square tests
##
##           Df  Chisq Pr(>Chisq)
## ss.lineage  6 17.151   0.008743 **
## ---
## Signif. codes:  0 '***' 0.001 '**' 0.01 '*' 0.05 '.' 0.1 ' ' 1
```

LSM <- lsmeans(max.temp,~ss.lineage,adjust="sidak")

contrast(LSM,"pairwise")

```
##  contrast       estimate    SE  df z.ratio p.value
##  2 - 3           -0.7789 0.369 Inf -2.112  0.3452
##  2 - 4           -0.1727 0.339 Inf -0.510  0.9987
##  2 - Baoshan     -0.2016 0.341 Inf -0.590  0.9971
##  2 - Int         -0.4266 0.346 Inf -1.235  0.8807
##  2 - Laos         0.4725 0.313 Inf  1.511  0.7384
##  2 - Yule        -0.4916 0.349 Inf -1.407  0.7984
##  3 - 4            0.6062 0.375 Inf  1.615  0.6729
##  3 - Baoshan      0.5773 0.378 Inf  1.527  0.7287
##  3 - Int          0.3523 0.382 Inf  0.923  0.9690
##  3 - Laos         1.2514 0.353 Inf  3.545  0.0072
##  3 - Yule         0.2872 0.385 Inf  0.746  0.9897
##  4 - Baoshan     -0.0289 0.349 Inf -0.083  1.0000
##  4 - Int         -0.2539 0.353 Inf -0.720  0.9915
##  4 - Laos         0.6452 0.321 Inf  2.009  0.4088
##  4 - Yule        -0.3190 0.357 Inf -0.894  0.9735
##  Baoshan - Int   -0.2250 0.356 Inf -0.633  0.9958
##  Baoshan - Laos   0.6741 0.324 Inf  2.079  0.3649
##  Baoshan - Yule  -0.2900 0.359 Inf -0.807  0.9843
##  Int - Laos       0.8991 0.329 Inf  2.737  0.0893
##  Int - Yule      -0.0650 0.363 Inf -0.179  1.0000
##  Laos - Yule     -0.9641 0.333 Inf -2.898  0.0577
##
## P value adjustment: tukey method for comparing a family of 7 estimates
```

cld(LSM)

```
##  ss.lineage lsmean    SE  df asymp.LCL asymp.UCL .group
##  Laos        1.17 0.214 Inf     0.599      1.74  1
##  2           1.64 0.241 Inf     0.998      2.29  12
##  4           1.82 0.253 Inf     1.139      2.50  12
##  Baoshan     1.85 0.257 Inf     1.158      2.53  12
##  Int         2.07 0.262 Inf     1.367      2.78  12
##  Yule        2.14 0.268 Inf     1.417      2.86  12
##  3           2.42 0.293 Inf     1.637      3.21   2
##
## Confidence level used: 0.95
## Conf-level adjustment: sidak method for 7 estimates
## P value adjustment: tukey method for comparing a family of 7 estimates
## significance level used: alpha = 0.05
```

#Contrast lineages inoculated onto Tropical japonica using least-squares means

max.trp<-clm(data=max_score_trp, notes~ss.lineage,Hess=T)

anova(max.trp)

```
## Type I Analysis of Deviance Table with Wald chi-square tests
##
##           Df  Chisq Pr(>Chisq)
## ss.lineage  6 44.117  7.009e-08 ***
## ---
## Signif. codes:  0 '***' 0.001 '**' 0.01 '*' 0.05 '.' 0.1 ' ' 1
```

LSM <- lsmeans(max.trp,~ss.lineage,adjust="sidak")

contrast(LSM,"pairwise")

```
##  contrast       estimate    SE  df z.ratio p.value
##  2 - 3           -1.5538 0.318 Inf -4.882  <.0001
##  2 - 4           -0.9725 0.310 Inf -3.137  0.0284
##  2 - Baoshan     -1.3442 0.322 Inf -4.171  0.0006
##  2 - Int         -1.6580 0.323 Inf -5.134  <.0001
##  2 - Laos        -1.0223 0.309 Inf -3.311  0.0162
##  2 - Yule        -1.8528 0.334 Inf -5.548  <.0001
##  3 - 4            0.5813 0.317 Inf  1.831  0.5268
##  3 - Baoshan      0.2095 0.328 Inf  0.639  0.9955
##  3 - Int         -0.1043 0.328 Inf -0.318  0.9999
##  3 - Laos         0.5315 0.316 Inf  1.680  0.6294
##  3 - Yule        -0.2990 0.338 Inf -0.884  0.9751
##  4 - Baoshan     -0.3718 0.322 Inf -1.154  0.9111
##  4 - Int         -0.6856 0.322 Inf -2.126  0.3372
##  4 - Laos        -0.0498 0.310 Inf -0.161  1.0000
##  4 - Yule        -0.8803 0.333 Inf -2.643  0.1132
##  Baoshan - Int   -0.3138 0.333 Inf -0.942  0.9657
##  Baoshan - Laos   0.3220 0.321 Inf  1.003  0.9536
##  Baoshan - Yule  -0.5086 0.343 Inf -1.482  0.7557
##  Int - Laos       0.6358 0.321 Inf  1.978  0.4286
##  Int - Yule      -0.1947 0.343 Inf -0.567  0.9977
##  Laos - Yule     -0.8305 0.332 Inf -2.502  0.1583
##
## P value adjustment: tukey method for comparing a family of 7 estimates
```

cld(LSM)

```
##  ss.lineage lsmean    SE  df asymp.LCL asymp.UCL .group
##  2         -0.284 0.215 Inf   -0.8609     0.293  1
##  4          0.689 0.221 Inf    0.0946     1.283   2
##  Laos       0.739 0.220 Inf    0.1486     1.328   2
##  Baoshan    1.060 0.238 Inf    0.4225     1.699   2
##  3          1.270 0.232 Inf    0.6478     1.892   2
##  Int        1.374 0.239 Inf    0.7343     2.014   2
##  Yule       1.569 0.253 Inf    0.8901     2.248   2
##
## Confidence level used: 0.95
## Conf-level adjustment: sidak method for 7 estimates
## P value adjustment: tukey method for comparing a family of 7 estimates
## significance level used: alpha = 0.05
```
