## Supplementary file 12 for "Ecological Differentiation Among Globally Distributed Lineages of the Rice Blast Fungus *Pyricularia oryzae*"

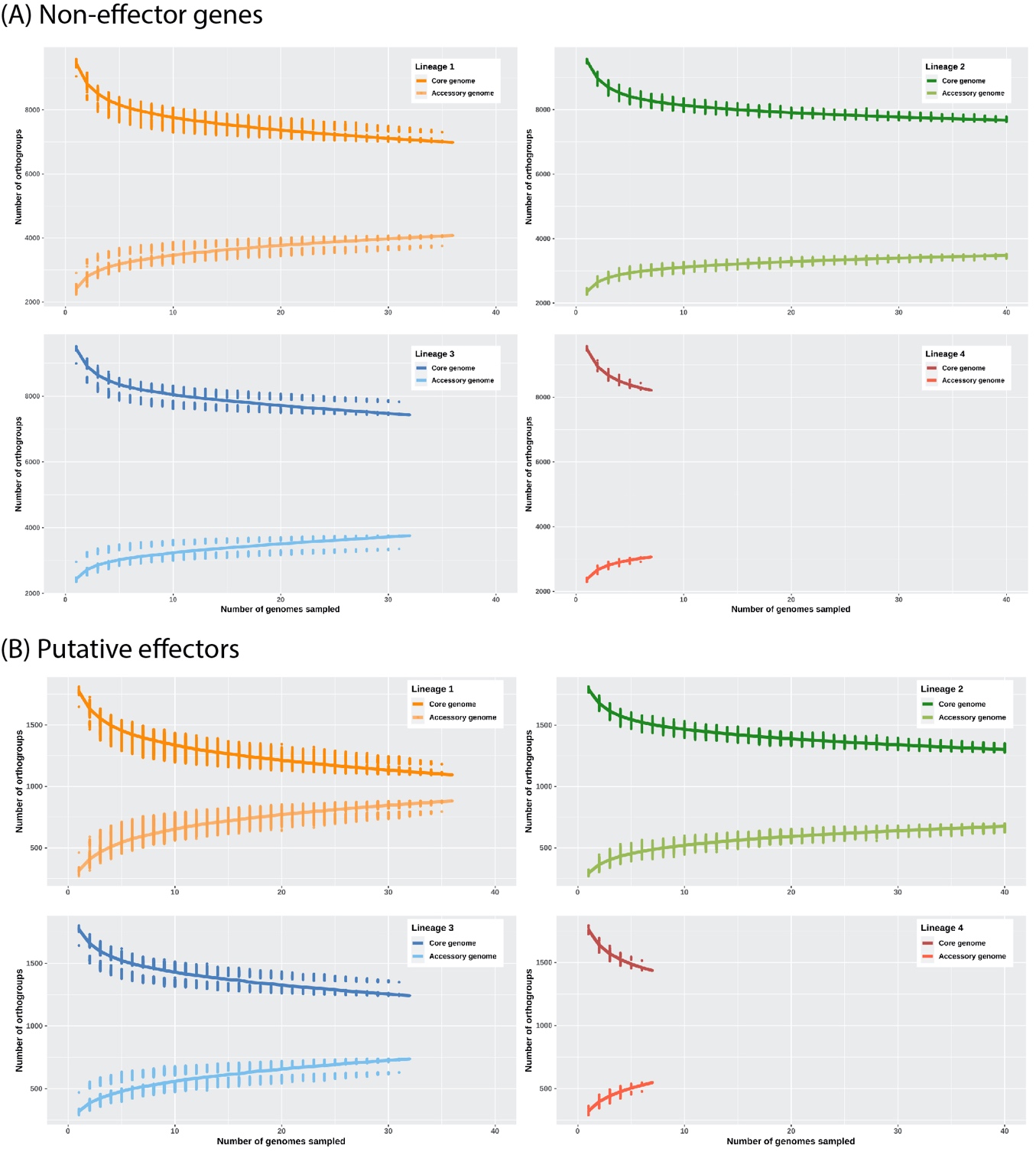

Supplementary file 12. Estimating the size of the core and accessory genome using a rarefaction approach. For each lineage, genomes were resampled in <=2000 combinations of N-1 genomes (N being the sample size). (A) Non-effector genes. (B) Putative effectors.
