## Supplementary file 13 for "Ecological Differentiation Among Globally Distributed Lineages of the Rice Blast Fungus *Pyricularia oryzae*"

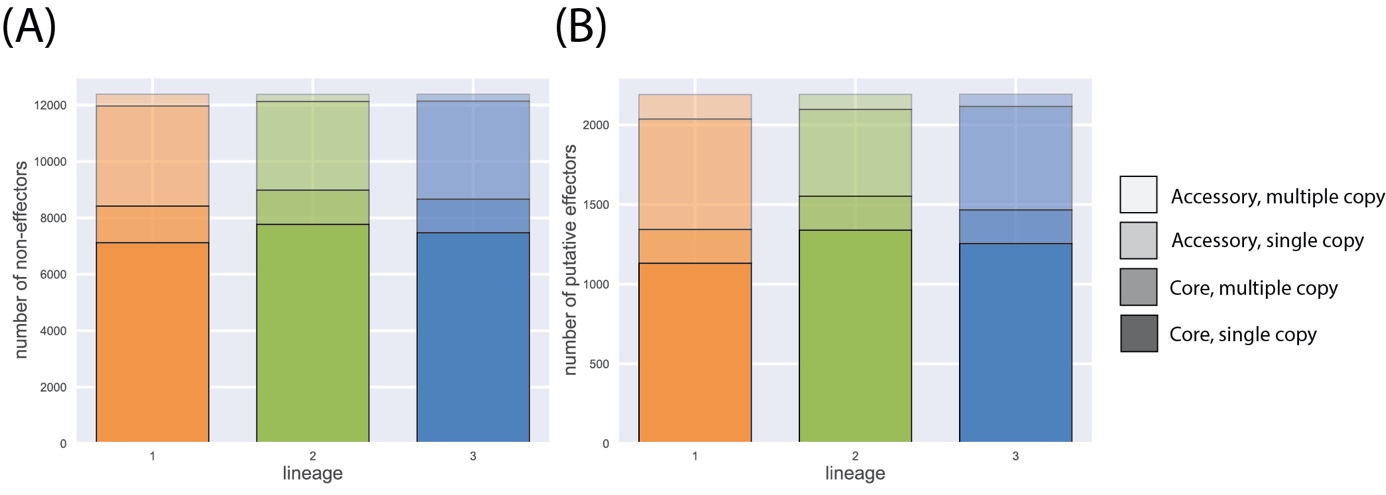

Supplementary file 13. Size of core and accessory genomes for non-effector proteins (A) and putative effectors (B) estimated using a rarefaction approach with a pseudo-sample size of n=30 genomes. Lineage 4 was not included due to small sample size.
