## Supplementary file 15 for "Ecological Differentiation Among Globally Distributed Lineages of the Rice Blast Fungus *Pyricularia oryzae*"

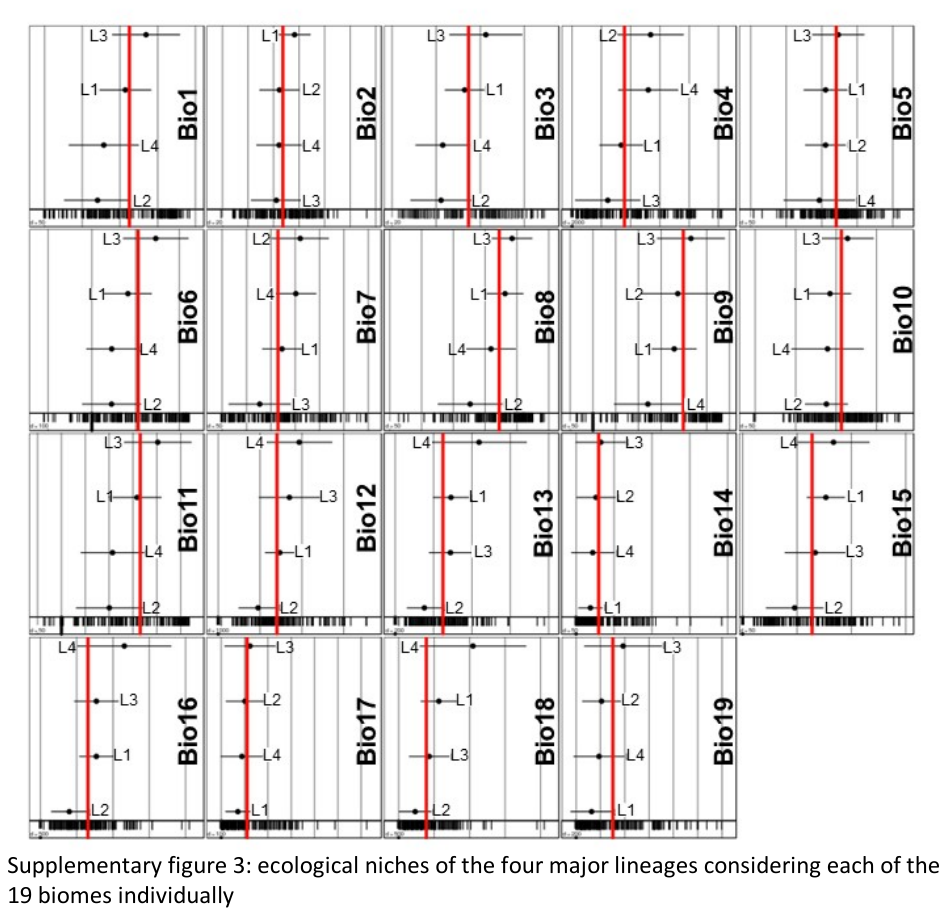

Supplementary file 15: ecological niches of the four major lineages (referred to as L1 to L4) considering each of the 19 biomes individually (referred to as Bio1 to Bio19). The x-axis represents the outlying mean index (OMI), which measures the distance between the mean habitat conditions used by a lineage and the mean habitat conditions used by the entire species, to test the hypothesis that different lineages are distributed in regions with different climates.
